# Genetic interactions shaping evolutionary trajectories in an RNA virus population

**DOI:** 10.1101/2020.01.16.908129

**Authors:** Chang Chang, Simone Bianco, Ashley Acevedo, Chao Tang, Raul Andino

**Affiliations:** Center for Quantitative Biology, Schools of Physics and Peking-Tsinghua Center for Life Sciences, Peking University, Beijing 100871, China; Industrial and Applied Genomics, IBM Accelerated Discovery Laboratory, IBM Research – Almaden, 650 Harry Road, San Jose, CA 95120-6099, USA; Department of Microbiology and Immunology, University of California, San Francisco, San Francisco, CA 94158, USA

## Abstract

A quantitative understanding of evolution rests on the analysis of the mutation accumulation process in biological populations, but is largely limited to high-frequency mutations due to the resolution of conventional sequencing technologies. Here, we examine the mutation composition of a poliovirus population over multiple passages using a highly-accurate sequencing strategy, that enables detection of up to 99% of all possible mutations, most of which are present at low-frequency. This data informs a mathematical model describing trajectory patterns of individual mutations to understand the type of interactions shaping population dynamics. We identify mutations consistent with a locus-independent behavior, and others deviating from that simple model by interactions. Clonal interference, followed by hitchhiking, appear to be the most prevalent interactions in the virus population. Epistasis, while presents, but does not significantly affect the distribution of mutational fitness on the short time scale examined in our study. Our study provides a comprehensive analysis of the allelic composition and how mutation rate, fitness, epistasis, clonal interference and hitchhiking influence population dynamics and evolution.

---

Understanding how mutations accumulate in biological populations is a central task in evolutionary biology. Factors such as mutation-selection balance^1–4^, genetic drift^5–7^, epistasis^8–13^, clonal interference^14–22^ and hitchhiking^17,23^, together determine the frequencies and dynamics of mutations in asexual populations. Accordingly, examining the role of these factors is critical to understand evolution^16,19,22,24–27^. However, conventional sequencing technologies have so far limited researchers to consider only high-frequency mutations^8–11,16,17^, ignoring the contribution and dynamics of minor alleles.

To reveal the principles of virus population dynamics and evolution across the entire spectrum of mutational abundances, we measured the genetic composition of 37 poliovirus populations obtained by serial passage using a highly-accurate sequencing technique. Totally 22,118 genetic variants are identified, most of which are of low abundance. A mathematical model is developed to understand and identify genetic interactions controlling distinct mutational trajectories. The overall impact of these genetic interaction factors on fitness landscape is also assessed.

## Results

### A highly-accurate sequencing strategy enables the detection of both high and low abundance genetic variants that follow complex dynamic patterns

To systematically investigate mutation dynamics in asexual populations we used CirSeq, a population sequencing method that enables reliable frequency estimations of both high and low abundance genetic variants^28,29^. We examined the genetic composition of 37 poliovirus populations obtained by serial passage in HeLa S3 cells at low multiplicity of infection (m.o.i. 0.1) (Supplementary Fig. 1) and large population size (10^6^). Given the high mutation rates per locus (~10^-6^-10^-4^), it was possible to calculate the frequency of a large number of genetic variants (22,118 from a total of 22,320 possible variants) (Methods). The majority of undetected variants mapped to the ends of the genome where CirSeq yielded low coverage. Based on the frequency of individual mutations we distinguished different classes of substitutions. First, a small number of mutations (~15) were observed at relatively high frequency (>0.04) (Fig. 1a). Within this high-frequency variant group some mutations follow a steady increase in frequency, consistent with the expected behavior of beneficial mutations or hitchhikers. Genetic hitchhiking^17,23^ occurs when a mutation changes frequency not because it is itself under natural selection, but because it is in the same genome with another mutation that is favorable under selection. Other mutations decrease in frequency after an apparent increase, a behavior consistent with clonal interference, when two (or more) mutations arise independently in different genomes, and if they are not combined in a single genome, their competition typically leads to a decrease in their frequencies and disappearance of some alleles from the population.

**Figure 1.**
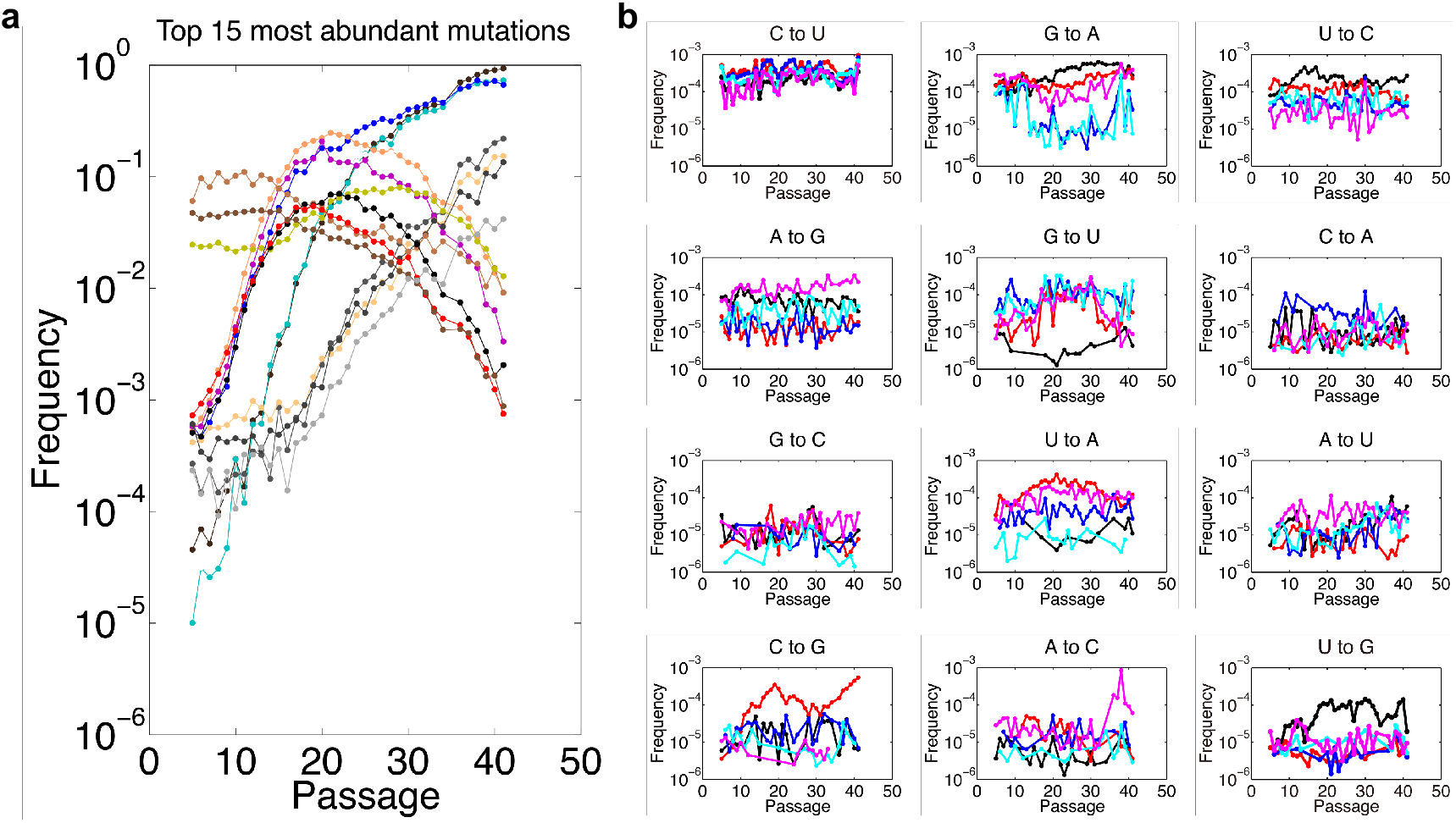
Evolutionary determinants of mutation trajectories. **a,** Trajectories of 15 highest frequency mutations. **b,** Samples of randomly selected mutation trajectories grouped by mutation type. Most randomly selected mutants shown similar order of abundance for each mutation type. **a,b,** Each color represents one particular mutation. Passage 35 is removed from data analysis during quality control of data processing (Methods).

Under our experimental conditions, the vast majority of mutations are present at very low abundances, and they follow complex dynamic patterns (Fig. 1b). We thus set out to understand the genetic interactions controlling distinct mutational trajectories across the entire spectrum of mutational abundances to uncover basic principles of virus population dynamics and evolution. Accordingly, we developed a mathematical model of mutation dynamics. Our experimental design, low m.o.i. and large population size at each passage, was designed to minimize genetic drift^5^, complementation^30^ and recombination^31,32^ between individuals in the population. Given that the change in frequency of a given variant over time is determined by mutation rate and selection (“unlinked mutations”, Fig. 2a), deviations from the expected pattern may stem from its interactions with other mutations (Fig. 2b).

**Figure 2.**
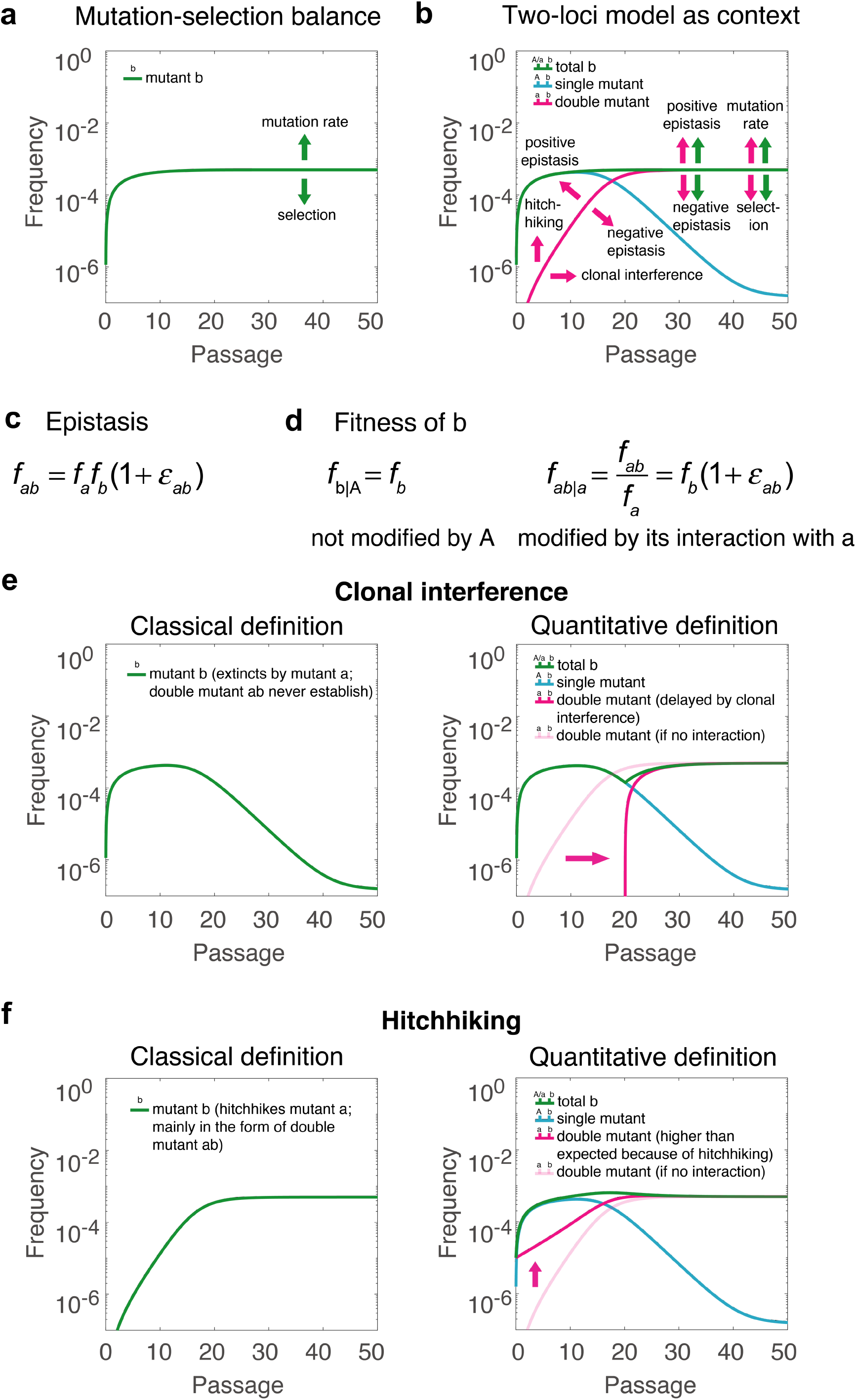
Multiple evolutionary factors shape mutation trajectories. A detrimental mutation is employed as an example to explain the effects or definitions of different evolutionary factors. **a,** In the context of a one-locus model, mutation rate and selection determine the trajectory of a detrimental mutation. The sample numerical trajectory is plotted with fitness 0.8 and mutation rate 10^−4^. **b,** In the context of a two-loci model, we give a numerical trajectory of a detrimental mutation *b* also with fitness 0.8 and mutation rate 10^−4^. The fitness of the dominant mutation *a* is 1.5, with a same mutation rate 10^−4^. The green curve represents the total frequency of mutation *b* in the population (observable data). The blue curve indicates the frequency of the single detrimental mutant, *b*, and the purple curve is the frequency of the double mutant *ab*, which will take over the single mutant as the major form of mutant *b* when the abundance of the dominant mutation *a* becomes large at its locus. (In this panel, only trajectories related to mutant *b* are shown. See Supplementary Fig. 3 for all numerical trajectories of mutants *a* and *b*). The fitness of the double mutant *ab* in this simulation is simply the product of the fitness of the single mutant *a* and *b* (no epistasis). The arrows represent the factors that could affect the trajectory of the mutation. An increase of the effect leads to a deformation of the curve along its direction. Clonal interference shifts the purple curve to the right (delayed formation of the double mutant), hitchhiking shifts it upwards (non-zero initial frequency of *b* in double mutant) (see their quantitative definitions in panels *e* and *f*), thus would not occur simultaneously on the same mutation trajectory. Positive and negative epistasis change the slope of the accumulation rate, as well as the mutation-selection balance, as they change the selection coefficient. See also Supplementary Figs. 4-6 for the detailed effects of different factors on trajectories of detrimental, neutral and beneficial mutations. **c,** The definition of epistasis. *f_a_* and *f_b_* represent the fitness of mutants *a* and *b*, and is the epistasis. **d,** The definitions of the relative fitness of mutant *b* with genetic background *A*(f_b|A_) and genetic background *a* (f_ab|a_) in the context of the two-loci model. In the case of *ε_ab_*=0, f_b|A=_f_ab|a_. **e,** The left panel illustrates the classical definition of clonal interference in the context of the two-loci model, i,e, the double mutant is unable to form and mutant *b* as a single mutant finally extinct. The right panel shows a quantitative, broader definition of clonal interference used in our paper. Using the trajectory of the double mutant predicted by the model without interaction as the frame of reference (the light purple curve; note that a one-locus model is equivalent to a two-loci model without epistasis, clonal interference or hitchhiking for a detrimental mutation), if the formation of the double mutant is delayed, we say it is under clonal interference. This definition includes the traditional definition of clonal interference as a limiting case of infinite delay. It also includes the situations when the extinction of mutant *b* is rescued by the delayed formation of the double mutant. **f,** The left panel illustrates the classical definition of hitchhiking in the context of the two-loci model, i,e, the formation of the double mutant. Under this definition, by the per locus high mutation rate in our data, all mutations would eventually hitchhike. The right panel shows a quantitative, narrower definition of hitchhiking used in our paper. Similar to that for clonal interference, using the trajectory of the double mutant predicted by the model without interaction as the frame of reference (the light purple curve), if the trajectory of the double mutant is higher than the one predicted by the one-locus model from the very beginning, we say it is under hitchhiking. **b,e,f,** The only observable variable in the two-loci model is the total frequency of mutant *b*. All other variables are latent variables. **c,e,f,** Those panels give the genetic interactions that are able to drive mutation trajectories departure from the simple mutation-selection model as shown in panel *b*.

### A mathematical model of mutation dynamics

We considered a virus population of genome length *L* with two possible alleles at a locus *k*, wild type and mutant, whose frequencies are denoted as *z_k_*(*t*) and *y_k_*(*t*) at passage *t*, respectively. The number of the possible genotypes containing each allele at this locus is 2^*L*−1^, and the abundance of these genotypes would change with time, thus making the total allele frequency and the fitness associated with each allele time-dependent. Forward and reverse mutation rates at this locus are indicated as 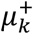 and 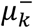, respectively (Supplementary Fig. 2a). The change in allele frequencies over time can be written as:

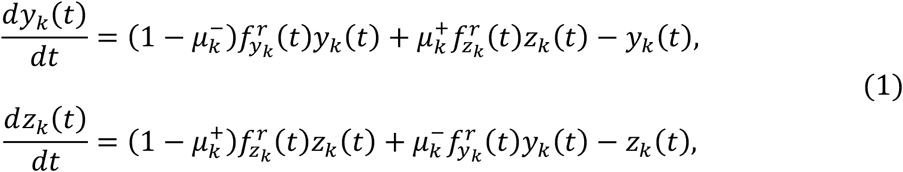

where 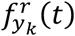 and 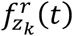 are the relative fitness of *y_k_*(*t*) and *z_k_*(*t*) over all possible genotypes containing each allele (Supplementary Fig. 2b). The dynamics of *y_k_*(*t*) are governed by mutant offspring produced by *y_k_*(*t*), mutants produced by *z_k_*(*t*) and the dilution term that keeps the population size constant. Although the model assumes two alleles at each locus, it can naturally be applied to situations with multiple alleles.

#### One-locus model for locus independence behavior

Previous studies have suggested that mutations with “unlinked” trajectories predicted by one-locus model may appear only for unrealistically large populations over a long time scale^33^. However, under our experimental conditions, we observed a number of mutations which behavior can be described by a single locus model, i.e., mutations behave independent and unlinked (Supplementary Fig. 2c and Methods). In these examples, neutral mutations accumulate linearly, beneficial mutations grow exponentially before saturating and asymptotically reaching fixation and detrimental mutations accumulate until reaching mutation-selection balance (Fig. 3a).

**Figure 3.**
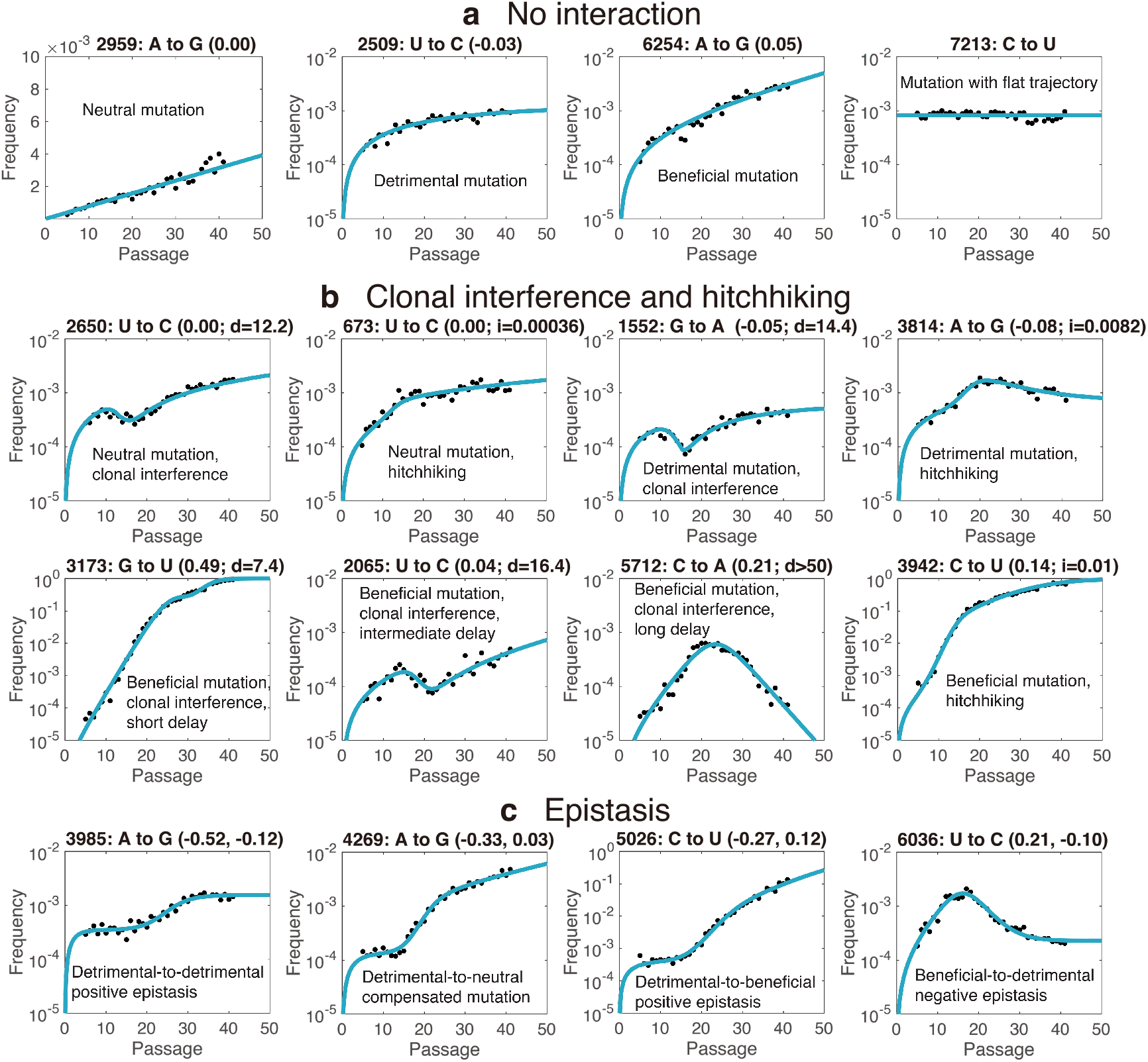
Mathematical model recapitulates the mutational behaviors observed in the poliovirus population. **a,** Examples of mutations showing no interactions (locusindependence): neutral, beneficial and detrimental mutations. The number in each pair of parentheses is the inferred selection coefficient of this mutation. For mutation with flat trajectory, inference of selection coefficient and mutation rate from the data is not possible. It belongs to no interaction under unlinked assumption, and hypermutation under linked assumption. Note that the plot for the neutral mutation A2959G is in linear space. Its semilog plot is available in Supplementary Fig. 11a. **b,** Neutral, detrimental and beneficial mutations under clonal interference or hitchhiking. The effect of clonal interference moves the curve of the double mutant toward right (Fig. 2b,e), which delayed the accumulation of the total abundance of the mutant. The effect of hitchhiking moves the curve of the double mutant toward upside (Fig. 2b,f), leading to a faster accumulation in the initial region, and even causing an abundance higher than the balance it should have. *d* is the delay between emergence and establishment of the mutation (see text), and *i* is initial frequency of [*ab*|*a*]. **c,** Epistasis between mutations can affect mutation frequency and the accumulation speed. The left three subpanels show that the trajectory of a detrimental mutation can be greatly influenced by interaction between beneficial and detrimental mutations, which can significantly alter the mutation-selection balance (3985: A to G), or allows it to accumulate at high frequency in the population (4269: A to G as compensated mutation, and 5026: C to U as detrimental-to-beneficial mutation). The right panel shows a beneficial mutation becoming detrimental following a strong negative epistasis event can significantly decrease in frequency (beneficial-to-detrimental mutation). The pair of numbers in the parentheses are the selection coefficients for f_b|A_ and the f_ab|a_ of this mutation. **a,b,c,** Detailed information for all these mutations is available at Supplementary Table 7. Also see Supplementary Figs. 4-6 for the effects of these evolutionary factors.

The assumption of mutations behaving independently from each other is reasonable during early passages, when the probability of more than one mutation per genome is low^28^. However, as time progresses, multiple mutations are likely to accumulate in single genomes^28^. In such a situation, a trajectory predicted by the locus-independent model will be still valid for this mutation if it does not have interactions with other mutations (C. Chang, *et al*., manuscript in preparation). On the other hand, epistasis between individual mutations, defined based on the effect of one allele on the fitness of another, is considered to be one of the mechanisms responsible for deviations from the independent locus model (Fig. 2b,c). Epistatic interactions have been proposed to play an important role in evolution^8–11^ but their effects on minor alleles and on the overall population dynamics have not been experimentally examined in detail.

#### Two-loci interaction model

To examine the contribution of interactions between mutations to virus population composition, we developed a two-loci interaction model (Supplementary Fig. 2c and Methods). We initially focused on epistasis between two beneficial mutations and between a beneficial and a detrimental mutation. Epitasis between two detrimental mutations is difficult to detected from the present data, since the abundance of detrimental mutations is usually low. To detect epistatic effects on a given mutation (*b*), its “modifier” (the interacting mutation, *a*) needs to be at high enough frequency for the double mutant (*ab*) to be generated and be detected over background, thus *a* should be at higher frequency than that expected for a detrimental mutation (Supplementary Fig. 7a). Accordingly, the frequency of the mutation *b, y_b_* (*t*), is given by the sum of the frequency of *b* when locus *a* is wild type (*A*) plus the frequency of the double mutant when locus *a* is mutant (*a*), thus giving

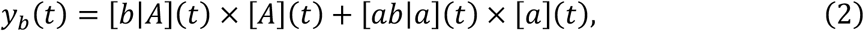

where [*b*|*A*] and [*ab*|*a*] are the conditional frequencies of *b* in the single and double mutants, and all terms on the right hand of the equation cannot be detected under our experimental conditions (i.e., latent variables). Mathematically, either of these above-mentioned two terms of conditional frequencies could be viewed as a variation of the one-locus model and thus equations for the latter could be directly used for the former, the frequencies of which can be determined by the relative fitness of mutant *b* in each genetic background (C. Chang, *et al*., manuscript in preparation). For example, the relative fitness of mutant *b* with genetic background *A* (f_b|A_) is defined as the fitness of the single mutant *b*, which determines the frequency of [*b*|A] (Fig. 2d), without any effect from other modifying mutations (“modifiers”; e.g. *a*). On the other hand, the relative fitness of mutant *b* with genetic background *a* (fab|a) is defined as the fitness of double mutant *ab* divided by the fitness of mutant *a*, with the effect from modifier *a* that epistasis has been implicitly considered (determining the frequencies of [*ab*|*a*]; Fig. 2d). Considering the trajectory of *b*, starting from a frequency dictated by the mutation rate at which *b* is acquired, fitness of individual mutations dominates the trajectory of *b* until the double mutant *ab* is high enough in the population. After a transition stage, where the abundance of the single mutant *b* and double mutant *ab* are at the same order (see the crosses of cyan and pink curves in Fig. 2b and in Supplementary Figs. 4-6), the fitness of *b* will be modified and dominated by the interaction with *a*. If no epistasis is present, f_b|A_ is expected to be equal to f_ab|a_ (Fig. 2d). For a detrimental mutation, the locus-independent model approximates well in the presence of a second mutation on the same genome (Supplementary Fig. 8a). On the other hand, for a neutral or a beneficial mutation, its trajectory could be slightly modified by another (beneficial) mutation on the same genome because of non-negligible higher-order terms of the hitchhiking effect in a deterministic numerical trajectory (Supplementary Fig. 8b,c; also C. Chang, *et al*., manuscript in preparation). These higher-order terms of hitchhiking, however, may not present in stochastic simulations in finite-sized population (Supplementary Fig. 8e), reducing the two-loci model to locus independent model in such a situation. Regardless the fitness, if the mutation frequency is modified by epistasis, it would violate the locus independence model and its frequency will deviate from the expectation without epistasis (Supplementary Figs. 4-6).

### Epistasis between beneficial and detrimental mutations

Our model describes, and our data fits, several distinct types (definitions in Supplementary Table 3) and strengths of epistasis (Fig. 3c). Pairwise strong epistasis between beneficial and detrimental mutations lead to trajectory shifting of the detrimental mutation from the mutation-selection balance that it is expected to reach in the case of no mutational interaction (compare Fig. 3a, U to C at position 2509 with Fig. 3c, left three panels). Epistasis modifies the trajectories of several mutations that do not longer fit to a one-locus model. We find examples of detrimental mutations under positive epistasis reaching a frequency higher than its original mutationselection balance (Fig. 3c, A to G at position 3985), or that even behave as neutral (Fig. 3c, A to G at position 4269, compensated mutation) or beneficial (Fig. 3c, C to U at position 5026). Conversely, a negative interaction leads to a lower balance, although is not observed in our data (an example numerical trajectory in the lower right panel of Supplementary Fig. 4; however, we also have real-data examples that be fitted by the clonal interference model as well as the model of detrimental mutation with negative interaction, with the fitting of the former better than the latter in these cases; see Supplementary Figs. 13-14).

To further examine the two-loci interaction model we conducted a number of simulations where parameters and variables can be thoroughly examined. We found that genetic interactions for a detrimental mutation with different beneficial mutations could lead to similar trajectories to each other (Supplementary Fig. 9). However, given this complexity, we are able to assess with sufficient confidence the strength of effective epistasis exerted at individual loci rather than obtaining specific pairs of interacting mutations (Methods).

### Clonal interference and hitchhiking

In addition to epistasis, clonal interference and hitchhiking are important and pervasive factors in asexual evolution^17^. In fact, we identified clonal interference and hitchhiking based on the specific characteristics of anomalous mutational trajectories, which also deviates from the locus independent mutational behavior (Fig. 2e,f and Supplementary Figs. 4-6). For instance, following a characteristic delay, clonal interference may reduce the frequency of mutations that are competing with other, fitter or more abundant, variants. In one example, the abundance of C5712A increase initially and then decrease over time (Fig. 3b). Our two-loci model (Fig. 2e, left panel) and simulations (“Clonal interference II” in Supplementary Fig. 5 and “Clonal interference III” in Supplementary Fig. 6), describe this trajectory considering that the double mutant *ab* never appeared in the population, which is one of the basic assumptions for clonal interference. However, we also observed trajectories in which some mutation frequency increase following an initial decrease (see Fig. 3b, neutral: U to C at position 2650; detrimental: G to A at position 1552; beneficial: G to U at position 3173 and U to C at position 2065). Our model suggests that in this case the double mutant *ab* is formed late or formed on time or early but its frequency is initially low due to clonal interference, thus a decrease is initially observed, but the formation of the double mutant eventually corrects that trajectory (right panel of Fig. 2e, pink arrow; and Supplementary Figs. 4-6). Accordingly, to identify mutations modified by clonal interference, we consider these two scenarios by introducing in our model a term describing a “delayed effect” of the formation of double mutant *ab*. Thus, as clonal interference has the effect of delaying the emergence of the double mutant, we replaced in equation (2) the conditional frequency [*ab*|*a*](*t*) with [*ab*|*a*](*t – delay*), to include an explicit time delay, with the time unit being number of passages. In the experimental data, we are able to observe the increase-decrease-increase pattern benefiting from the exceptionally high per-locus mutation rate.

On the other hand, hitchhiking occurs when a mutation that is not under natural selection links to another selectively favorable mutation. Besides neutral mutations, beneficial or detrimental mutations can also change frequencies by linking to another beneficial mutation (Supplementary Figs. 4-6; also see McDonald, Rice and Desai^32^). Such a mutation can accumulates faster or reach higher frequency than it would have if it were only subject to its own selection, or even be able to be fixed in the population. In the context of the two-loci model, hitchhiking is equivalent to a nonzero frequency of double mutant *ab* (left panel of Fig. 2f). For a large-sized asexual population with minimized recombination, once mutant *a* reach high abundance, hitchhiking is unavoidable since the double mutant *ab* is natural to happen because of high per-locus mutation rate. It makes most, if not all, mutation trajectories examples of hitchhiking. For example, the trajectory of a mutation with no interaction seems to be unlinked (the green curve in Fig. 2b), but its genetic constitution is initially the single mutant (cyan curve) and then the double mutant (the purple curve). Thus, this linked mutation can have an “unlinked” trajectory because of hitchhiking. Similarly, mutations with epistasis or even clonal interference can be examples of hitchhiking, as long as they have nonzero frequencies of double mutant *ab* (see the purple curves in Supplementary Figs. 4-6). The only situation explicitly not belonging to hitchhiking is that clonal interference completely prevent the present of the double mutant. Thus, it would be improper to use hitchhiking to describe all co-occurrence phenomena because its effect is so broad. We adopt a narrower and more specific definition of hitchhiking in our paper. For a given mutant, if its trajectory can be better explained by a hitchhiking model of which its double mutant frequency (the purple curve in Fig. 2f) is higher than that predicted by the model without interaction (the light purple curve), it will be considered under hitchhiking. The initial frequency of [*ab*|*a*] in the two-loci model is the key parameter to describe and approximate the effect of hitchhiking on a mutation trajectory that deviates from an “unlinked” behavior (Methods). In the data, we find examples of hitchhiking for neutral (U673C; semi-log plot in Fig. 3b and linear plot in Supplementary Fig. 11b), beneficial (C3942U in Fig. 3b) and detrimental mutations (A3814G in Fig. 3b).

We further expect that many epistasis events related to beneficial mutations to be masked by clonal interference or hitchhiking, and thus are unlikely to be identified (Supplementary Table 11). Beneficial-to-beneficial mutations (*f*_b|A_ > 1, *f*_ab|a_ > 1), both positive and negative epistasis, are vulnerable to the effect of clonal interference and can have deformation in trajectories when such an effect is present (Supplementary Fig. 10a). Even if the effect of clonal interference is not introduced, beneficial-to-beneficial mutations with negative epistasis or beneficial-to-lethal mutations (*f*_b|A_ > 1, *f*_ab|a_ ≈ 0), can have similar trajectories to that of non-epistasis mutations under clonal interference or hitchhiking in simulation (Supplementary Fig. 10b,c). However, we were able to detect beneficial-to-detrimental mutations (*f*_b|A_ > 1, *f*_ab|a_ < 1; Fig. 3c, U to C at position 6036 is an example; also see Supplementary Table 8b).

In general, clonal interference and hitchhiking are not thought to act on one mutation simultaneously, since the former would slow or even prevent the accumulation of the double mutant, but the latter, on the other hand, would improve the accumulation (Fig. 2b). In spite of these general considerations, as is mentioned above, it may happen in some conditions, that trajectories under clonal interference, hitchhiking or epistasis may appear similar. Thus, whenever a mutation can be classified, distinguishing between clonal interference, hitchhiking and epistasis is critical to understand the mechanisms controlling virus population dynamics.

### Model selection to determine the best interaction model for a mutation trajectory

For this purpose, we have devised a model selection procedure based on one-locus and two-loci models to determine the best interaction model for each mutation’s trajectory (Methods). This consists of a scoring procedure of models with increasing complexity, where complexity here is measured by the type of the genetic interaction and the number of parameters allowed in the model (we use 6 levels of complexity, see Supplementary Table 1). Every possible interaction (no interaction, clonal interference, hitchhiking, and positive/negative epistasis) was considered for mutations at every locus. Model fitting was performed in both linear and log space to properly consider the weights of both high and low frequency data. For models of the same level of complexity, the best model is selected based on goodness of fitting in both linear and log space in order to properly consider data of both high and low abundance. For models at different complexity levels, a Bayesian Information Criterion^34,35^ is also used to assess significance of model selection. This method tends to penalize a more complex model (with more parameters) unless it fits significantly better than a less complex one. Notably, in order to guarantee the accuracy of our epistasis predictions, if a mutational trajectory can be better explained by clonal interference or hitchhiking, it will be classified as non-epistatic even in the case where the fit to either of these two models is not statistically different from the fit to the epistatic model, although the numbers of parameters are the same (Supplementary Table 1). To exemplify our procedure let us consider the mutant at position 4269 (A to G; Fig. 3c). By fitting in both linear and log space, the best candidate model of each level of complexity is obtained (Fig. 4 and Supplementary Fig. 12). Among those, the detrimental-to-beneficial two-loci epistatic model stands out as the one that fits the data best (significantly better than the competitive “neutral mutation under hitchhiking” model), with its f_ab|a_ very close to 1, thus indicating that it is a compensated mutation (Supplementary Table 3).

**Figure 4.**
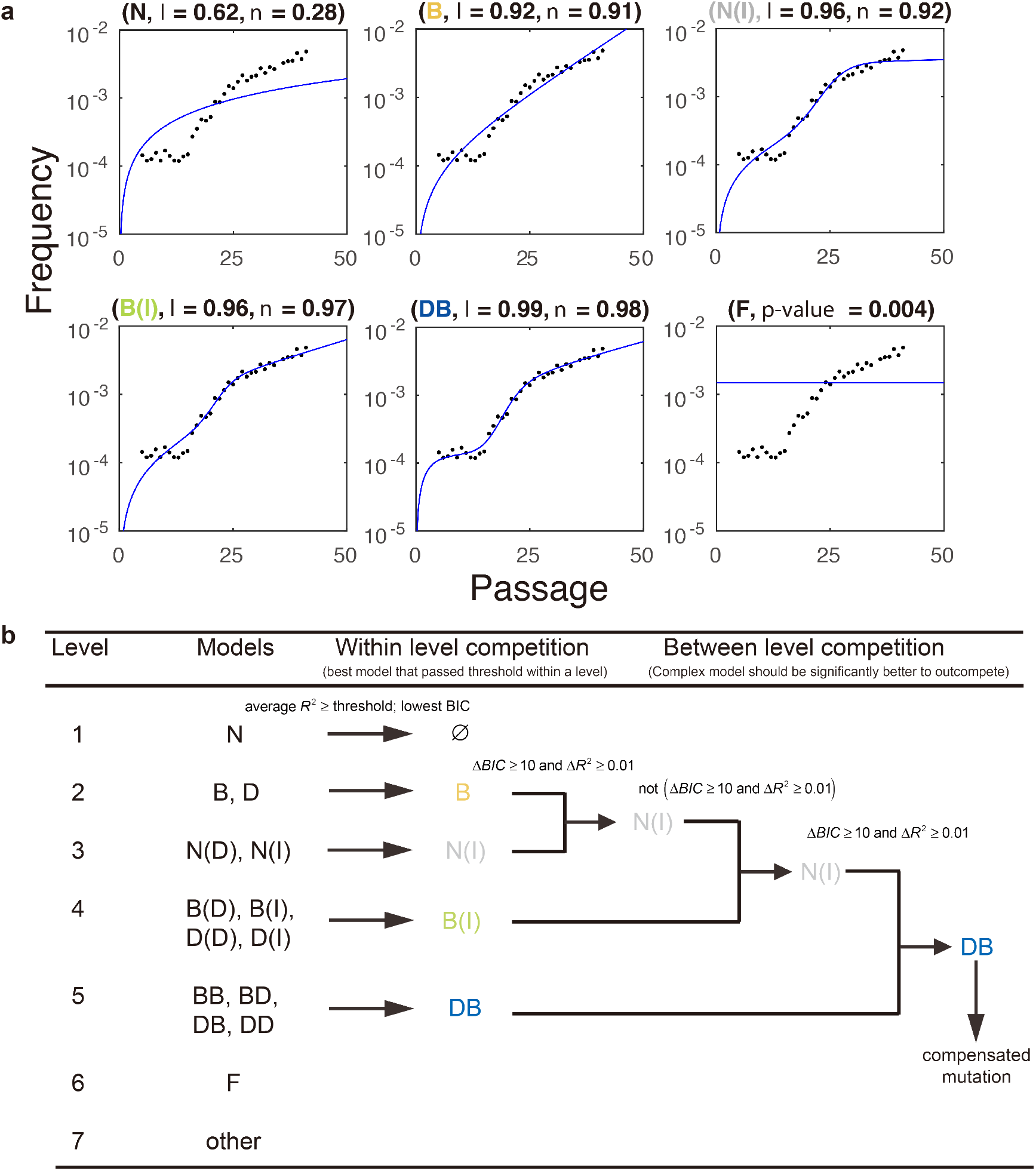
An example of model selection. **a,** In this plot, different panels represent the fitting of different models to the trajectory of the mutation A to G at position 4269 (Fig. 3c). We only included several representative models in this plot, according to the fitting results illustrated in the panel *b*. The full version is available at Supplementary Fig. 12. In titles of each panel, one-letter code represents one-locus model, two-letter code is a two-loci epistasis model (the first letter represents the model for f_b|A_, and the second letter represents the model for f_ab|a_). For instance, *DB* is the detrimental-to-beneficial two-loci epistasis model. The numbers following *l* and *n* are R^2^ in log space and normal space. For each letter of a code, *N* is neutral, *D* represents detrimental, and *B* means beneficial. *D* or *I* in the parentheses means this model is the version of clonal interference or hitchhiking. An exception is *F*, which represents mutation with flat trajectory. Note that in this *F* model, a larger p-value suggest a better fit (*i.e*., the data could be explained by the null model under larger p-value). Also see Supplementary Table 1 for the code. In the model selection process, every one-locus and two-loci models are used to fit the data, but only those models fit well enough in log and normal spaces are considered. For models in each level (Supplementary Table 1), the best one is selected (with its model code being marked with color). The best model that can explain the data is chosen from best models in each level according to the procedure described in *Methods* and here illustrated in the panel *b*. **b,** A flow chart of model selection. For each level of models, the best model with average R^2^ passing threshold is selected. The threshold is set to be 0.8 for non-epistatic models (model levels 1-4) and 0.9 for epistatic models (model level 5). For models in different levels, the more complex model should be significantly better to outcompete other models, which requires better average R^2^ and better BIC.

### Landscape

Under the assumptions of the model, our classification uncovers the underlying evolutionary dynamics for the majority of mutation trajectories of the data (Supplementary Table 5). For mutations having a trajectory that allows for assessment of fitness (N=1226), non-epistatic mutations count for the majority (N=1198). Half of these mutations are under clonal interference (N=644), the majority of which are beneficial or neutral mutations (Fig. 5a,b). Most of the inferred hitchhikers (N=329) are either neutral or deleterious variants, with a few of them being beneficial (Fig. 5a,b). Most of the inferred locus-independent mutations have a fitness near neutral (Fig. 5a).

**Figure 5.**
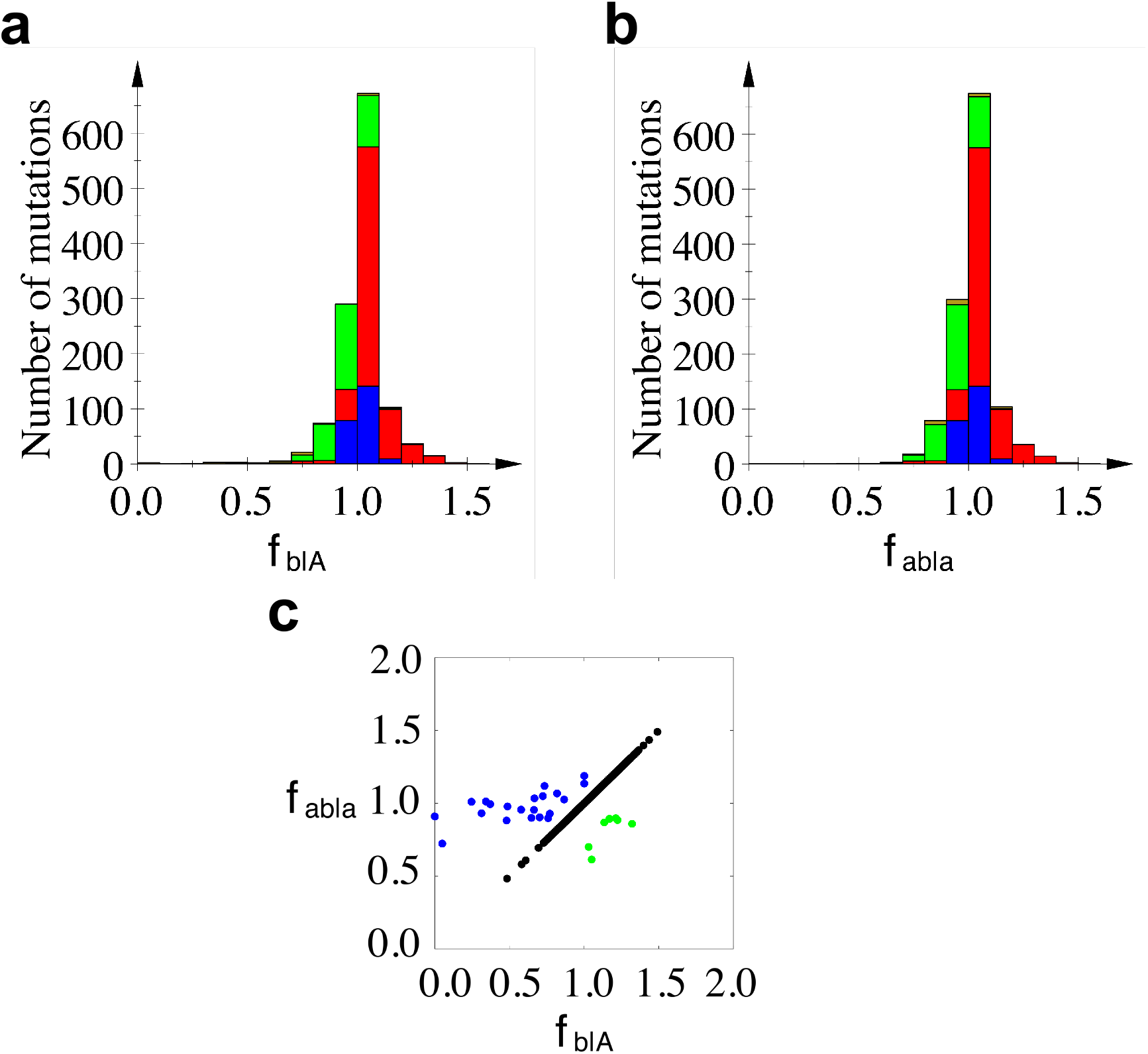
The fitness landscape. The distributions of **(a)** f_b|A_ and **(b)** f_ab|a_. **a,b,** Colors represent different subtypes of mutations: no epistasis (blue), clonal interference (red), hitchhiking (green) and epistasis (brown). **c,** Scatter plot of f_b|A_ *vs*. f_ab|a_. Mutations are color coded by the type of epistasis (black for no interaction, blue for positive epistasis, green for negative epistasis).

Mutations under epistasis represent a relatively small proportion of the total (N=28; 2.3%), mostly being positive interactions (Supplementary Fig. 15d). Analysis of fitness landscapes of f_b|A_ and f_ab|a_ showed that the overall distribution of mutational effects is not significantly affected by epistasis in the time scale of our experiments (Fig. 5a,b), during which several mutations almost fixed (Fig. 1a), suggesting that epistasis is not a fast time-scale factor for changing fitness landscape in evolution. Interestingly, we also observed a compensatory effect of epistasis, resulting in an increase in fitness for many detrimental mutants (Fig. 5c).

## Discussion

Over 60% of all mutations were classified as mutation with flat trajectory, those that show low frequency and no observable initial accumulation phase (Fig 3c, right panel), which are predicted as detrimental mutations with low fitness under the locus independence assumption (≤0.6; estimated by Eq. (7)). This includes a large number of mutations that are considered to be with high mutation rates (Supplementary Table 6; the magnitude of mutation rates of different mutation types is available in Acevedo *et al*.^28^). Under the locus independence assumption, we show that both mutation rate (peaked at 10^−5^ in simulation) and strong negative selection could contribute to the formation of a mutation with flat trajectory, which could be further enhanced by the presence of clonal interference (Supplementary Table 14). However, for a detrimental mutation, when the mutation rate is high (≥ 10^−5^), clonal interference solely cannot explain a mutation with flat trajectory unless its negative selection is strong enough (Supplementary Table 14), indicating loci with high mutation rates are generally under strong negative selection. Alternatively, hypermutation could introduce multiple C to U or G to A mutations (those with the highest mutation rates in Acevedo *et al*.^28^) to the same genome, thus largely increasing the likelihood for the aggregated mutations being detrimental and could lead to a flat trajectory if the overall fitness is in the parameter range that no initial accumulation phase is observed. Considering that the observation of mutations with flat trajectory is not limited to C to U or G to A (Supplementary Table 6), both mechanisms could contribute to the observation of pervasive mutations with flat trajectories. Interestingly, we observed the enrichment of mutations with flat trajectory in coding regions of capsid proteins (Supplementary Fig. 17).

In our study, the number of loci in the interaction model is set to be two. Though higher-order epistasis involving multiple loci is possible, we expect this higher dimension interactions to be infrequent under our experimental conditions (e.g. 37 passages), given that these mutations need to be of high abundance (Supplementary Fig. 7b).

Overall, our analysis revealed the enormous dynamical range of complex interactions of a viral population. This is even more striking considering the weakly-selective nature of the experimental environment. While low selection allows for wide exploration of the genetic landscape with little or no penalty, our work highlights the exceptional potential of genetic interactions that can occur during an infection. Hence clonal interference appears from our analysis to be the most prevalent means of interaction between mutations, followed by genetic hitchhiking. Epistasis is present, but not pervasive, and does not significantly affect the distribution of mutational fitness on the short time scale examined here.

We have shown, using a combination of CirSeq data analysis and mathematical modeling, that a variety of mutational dynamics can be assessed through analysis of the trajectories of individual mutants. Despite the fact that pairs of interacting mutations are elusive, by taking into account the major factors including mutation rate, fitness, epistasis, clonal interference and hitchhiking (Fig. 2), information about the evolutionary forces acting on individual mutations can be revealed.

## Acknowledgements

This work was supported by NIH (R01, AI36178, AI40085, P01 AI091575) and the University of California (CCADD), and DoD-DARPA Prophecy to R.A.; and by Chinese Ministry of Science and Technology (2015CB910300), National Natural Science Foundation of China (91430217). A.A. was supported by NSF GRF. C.C. acknowledges the support from the PhD Student’s Short-term Overseas Research Program of Graduate School of Peking University. We thank Hao Li, Xin Wang, Xiaojing Yang, and Minghua Deng for helpful discussions, and Xiang Liu and Hao Shen not only for helpful discussions, but also for checking the mathematics.

## Author contributions

C.C., S.B., C.T. and R.A. conceived the study. C.C. built the mathematical models, carried out simulations and data analysis, and prepared the initial manuscript. S.B. helped the design of simulations that lead to the concept of trajectory shifting under epistasis, and extensively improved the manuscript. A.A. and R.A. designed the experiments. A.A. performed the experiments. C.T. and R.A. directed the project. All authors improved and approved the manuscript.

## METHODS

### Passaging of poliovirus population

Poliovirus type 1 Mahoney populations were obtained by passaging through HeLa S3 cells as described in Acevedo *et al*.^28^. Specifically, a single wild type poliovirus clone was isolated by plaque purification (passage 0) and amplified to obtain a high titer stock (P1). For all subsequent passages, 10^6^ plaque forming units (p.f.u.) were used to infect a HeLa S3 monolayer at a multiplicity of infection (m.o.i.) 0.1 for 8 hours (approximately one replication cycle). Virus was harvested by three freeze-thaw cycles and clearing of the supernatant by centrifugation.

### Production and sequencing of CirSeq libraries

As described in Acevedo *et al*.^28^, each passage was amplified in HeLa S3 by a single high m.o.i. infection. At 6 hours post infection, total cellular RNA was isolated by TRIzol extraction and virus RNA was enriched by poly(A) purification. Libraries were produced as described in Acevedo and Andino^29^. Briefly, virus RNA was fragmented, circularized and reverse-transcribed to generate cDNAs containing tandemly-repeated virus sequences. These cDNAs were further cloned by second strand synthesis, end repair, dA-tailing and ligation of Illumina TruSeq indexed adaptors. Libraries were sequenced on either the MiSeq or HiSeq 2500 platforms for 300-323 cycles.

### Quality control of the data

Mutations which show long periods of absence (that is, >10 passages) have been removed from the analysis. This brings the number of mutations down to from 22,118 to 21,031. We also removed all data of passage 35 from analysis because it has unreasonable bias.

### Establishment and observability of a mutant

According to the theoretical analysis of Desai and Fisher^3^, a beneficial mutation with number larger than *n*~1/*s* can be established in a population with a sufficiently small mutation rate (*s* is the selection coefficient). Under the effect of genetic drift, a lineage with size *n* would need *n* generations to change by order *n*. A beneficial mutation with selection coefficient *s* on average can generate *ns* more offspring per generation, and *n*^2^*s* for *n* generations. Thus, a beneficial mutation can establish when *n*^2^*s* >n or *n* > 1/*s*.

In our disease model, differently from the work of Desai and Fisher, mutation rate is not negligible and needs to be explicitly considered as follows. For a beneficial mutation, the number of *de novo* alleles obtained at mutation rate *μ* for *n* generations would be *μPn*, where *P* is the total number of individuals in the population. Considering both contributions by mutation rate and by selection, *μPn* + *n*^2^*s* > n is needed for establishment, i.e., 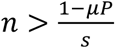. Within our parameters range, once a beneficial mutation has emerged, the effect of genetic drift is limited, since *μP* ≥ 1. Thus, the contribution by mutation rate along is sufficient for the establishment and the observability of a beneficial mutation. We further consider the stochasticity of generation by *de novo* mutations by assuming that the number of offspring contributed by mutation rate is Poisson distributed with average *μP*. When *μP* = 3, the probability of generating no offspring is about 5%. This implies that, for mutation rate larger than 3 × 10^-6^, a beneficial mutation will always be present in the population and escape from the fate of extinction by genetic drift. This analysis does not consider the occurrence of extinction by clonal interference.

For a neutral mutation to establish in the population, one also needs to have *μP* > 1. This condition needs to hold in order for a detrimental and even lethal mutation to survive and present in a population, and helps explaining how the whole spectrum of mutations, including strongly deleterious, can be observed in our data.

For a double mutant, it is able to establish and be observed in the population if the abundance of genomes with the “dominant” mutation(s) (denoted as *A*) and the mutation rate of the “subordinated” mutation (still denoted as *μ*) is large. Consider that *A* would be smaller than the total population size *P, μ* needs to be higher than that required by the single mutant analysis.

### A mathematical model to capture mutation dynamics

For each passage of the serial passage experiments, 10^6^ p.f.u. were used to infect HeLa cells with 0.1 m.o.i., minimizing genetic drift, complementation and recombination between individuals in the population. Within one passage, when a virus successfully infects a cell, it can generate tens of thousands of offsprings at the end of this passage. However, only a tiny fraction would be sampled to infect cells in the next passage, given that the p.f.u. to infect each passage is fixed. In evolution literatures, the definition of fitness is based on the number of reproducible offsprings. Thus, in this system, the average fitness of the population could be normalized to 1, and the fitness of a specific mutant or genotype needs to be normalized by the total number of offsprings generated by all viruses during this passage.

Let *y_k_*(*t*) and *z_k_*(*t*) denote, respectively, the frequency of mutant and wild type alleles at locus *k* at passage *t*. Their time-dependent average fitness are 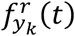 and 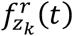. The forward and reverse mutation rates at locus *k* are 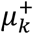 and 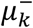. Thus, in passage *t* + 1, the allele frequency could be given by

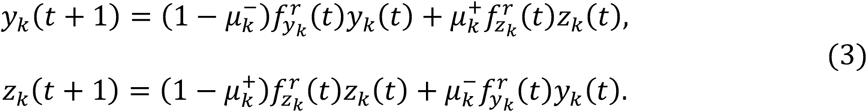

By subtracting the frequency at passage *t* for each equation in Eq. (3) and use a continuum approximation, we could easily get Eq. (1) in the main text.

Although Eq. (1) is too complex to be solved analytically, model reduction techniques allow for approximate solution (Supplementary Fig. 2c).

### One-locus model

When a mutation at locus *k* is independent, i.e. it does not interact with any other mutation at other loci, it is not a hitchhiker and it is not under clonal interference, the ratio between passage-dependent fitness can be simplified as 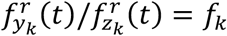, i.e., the mutant’s own fitness. Also, backward mutations are neglected in our analysis. Under locus independence, the full model can be reduced to one-locus model. This reduction holds even when the population size is finite (C. Chang, *et al*., manuscript in preparation).

A locus-independent neutral mutation does not change the fitness of the genome. Thus, its accumulation is linear in time and its frequency is approximatively equal to the forward mutation rate times the time, or:

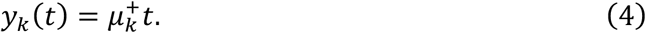

A locus-independent beneficial mutation accumulates as follows

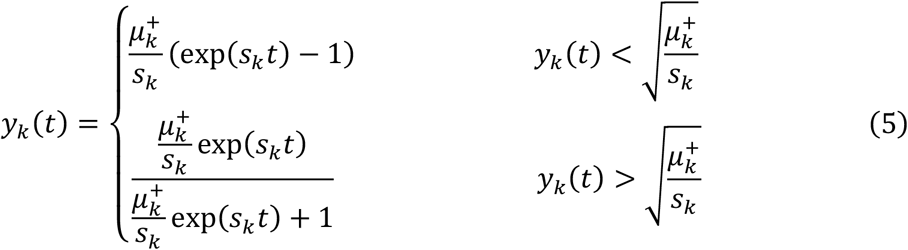

where *s_k_* = *f_k_* – 1 is the selection coefficient. By convention, a positive selection coefficient indicates a beneficial mutation 0 is neutral, and negative a detrimental mutation.

The frequency of a locus-independent detrimental mutation in time is given by

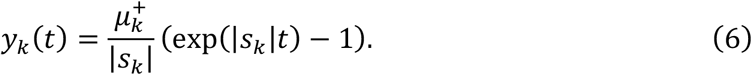

If a mutation is observed at a frequency in the population at a given time, we can estimate the number of passages that it takes it to reach its mutation-selection balance 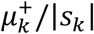

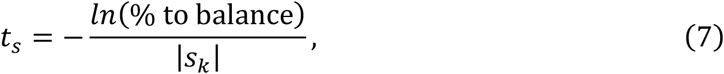

where the numerator represents the natural logarithm of the ratio between observed mutations selection balance and the asymptotic one.

### Two-loci model: epistasis

Our two-loci model captures all three kinds of interactions: epistasis, clonal interference, and hitchhiking (Fig. 1c). Numerical simulations displaying the several dynamic behaviors that characterize the interactions are shown in Supplementary Figs. 4-6.

Let us consider first the case of epistatic interactions, without clonal interference or hitchhiking. Epistasis can occur through combinations of beneficial (positive selection coefficient) and deleterious (negative selection coefficient) mutations. From a modeling perspective, we consider epistasis involving neutral mutations as a special case of epistasis between detrimental or beneficial mutations.

We use a multiplicative definition of epistasis^36^. Mathematically, the frequency of a “dominant” mutation at locus *a* can be directly obtained by Eq. (5), which holds even when *a* is simultaneously interacting with other mutations besides mutation *b* (C. Chang, *et al*., manuscript in preparation). Eqs. (4), (5) or (6) can be used to obtain [*b*|*A*] and [*ab*|*a*] (Supplementary Table 1). The mathematical form above allows for enumeration of all possible cases of pairwise epistasis.

### Two-loci model: clonal interference and hitchhiking

Clonal interference and hitchhiking are reported to be pervasive in asexual evolution^17^. In simulations, even only one beneficial mutation with large fitness in the genome is enough to induce substantial hitchhiking (Supplementary Fig. 21). Concurrently, clonal interference becomes an important effect when multiple beneficial mutations with large fitness appear in the population (Supplementary Fig. 20). It is known that the ultimate consequence of clonal interference is extinction of a beneficial mutation through competition^16–19^. However, clonal interference can lead to various mutation dynamics when the per locus mutation rate is extremely high. This is due to the aforementioned effective time delay of establishment of double or multiple mutations. Either of these effects may severely affect mutation trajectories.

Using the curve of double mutants predicted by the model without interaction as the frame of reference, clonal interference and hitchhiking shift the curve to different directions, but these two effects are unlikely to happen simultaneously (Fig. 1c). In general, we observe that hitchhiking shifts the frequency of the double mutant upwards. This effect is akin to assigning the conditional frequency [*ab*|*a*] a non-zero initial condition. Clonal interference has the effect of delaying the emergence of the double mutant (rightward). This is similar to replacing the conditional frequency [*ab*|*a*](*t*) with [*ab*|*a*](*t – delay*), that is, to include an explicit time delay, with the time unit being the passage. If the delay is long enough, it would lead to the classical phenomenon of clonal interference, with mutations under this effect eventually lost in the evolution. On the other hand, if the delay is finite, we would observe the rescue of mutations with an initial decrease as the cases observed in our data.

In the context of the two-loci model, f_b|A_ and f_ab|a_ would still be the same under the effect of clonal interference or hitchhiking, if epistasis is not present.

### Model selection

We use a model selection procedure to identify which of the considered models best describes the behavior of each mutation (Fig. 4 and Supplementary Fig. 12), and classify mutations into different subtypes (Supplementary Table 3). We consider the simplified one-locus and two-loci models derived from the full model to fit the data. The structure of the models reveals six potential classes (Supplementary Table 1). Classification of mutation behaviors is done by iterating over models of increasing complexity, and priority is given to models of lower complexity (Fig. 4b). In general, the one-locus model is preferred over two-loci model, since a two-loci model can recapitulate the one-locus model in certain conditions. Analogously, one-locus neutral will have priority over every other model, since it is a limiting case of all models. Moreover, in our model clonal interference and hitchhiking take precedence over epistasis in order to guarantee the accuracy of predicting epistasis. We do not consider a “second order” combination of epistasis, clonal interference and hitchhiking, which would lead to higher false positive rate for predicting epistasis. If none of the above-mentioned models can capture the dynamics of a mutant, we use one sample t-test to determine if a mutation is mutation with flat trajectory (we use the log of abundances in the test). We use R^2^ and log (R^2^) to estimate model goodness of fit, while we adopt Bayesian information criterion (BIC) for quantitative model selection^34, 35^. Simulation suggests that when the sum of the squares of the residuals is very small (i.e., R^2^ is very close to 1), a small improvement in R^2^ leads to a very large increase of BIC. In this situation, the simpler model is actually good enough to explain the data. Thus, both BIC and R^2^ are used in concert to determine the significance for model selection (a complex model is significant if Δ*BIC* ≥ 10 and Δ*R*^2^ ≥ 0.01).

### Determine start passage

Although the first sequenced passage is marked as P2 (i.e., second passage) in experiments, it is unclear what is the actual or effective start passage for the data. In our model, the passage is an independent variable. Thus, the starting passage number should be considered as a global parameter in the inference process.

Intuitively, neutral or near neutral mutations in the population can help us to infer this parameter. By linear fit of mutations that are very likely to be neutral, we could get their intercepts with x-axis (passage axis), which would be the position of passage 0. In simulation, we found that, given the population size of 10^6^, this method is unable to infer the start passage when the mutation rate is lower than 10^-5^, so we only count mutations above this mutation rate. Furthermore, weighted average of the inferred start passages using 1/R^2^ as weight can improve the accuracy of this method. The simulation results are available in Supplementary Table 4 and the inferred start passage for the real data is 4.26 (N=69).

Besides the inference method by neutral mutations, we also developed a probabilistic method to infer the start passage, by which we try to maximize the probability of having a start passage given the data (maximum a posteriori estimation). The posteriori probability is given by

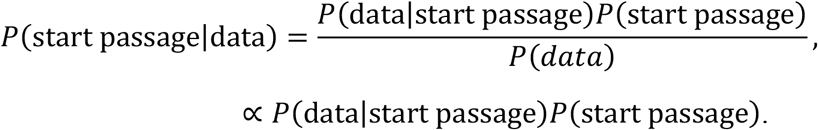

Since we have no preference on the start passage, a flat prior for the start passage is used. This optimization problem converts to maximize P(dataļstart passage), thus

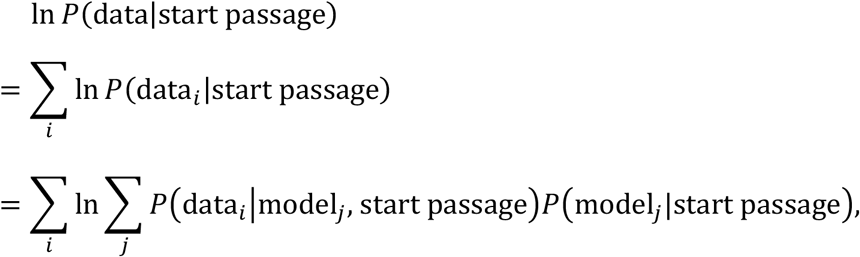

where *i* and *j* are the indices for data and model, assuming that the observed mutation trajectories could be generated from the models we derived. For *P*(model_*j*_|start passage), again we have no preference, so we use the average of all models we derived. *P*(data_i_|model_j_, start passage) could be directly calculated from the BIC of model_j_^37^. In simulations, we found that directly using the start passage that maximize ln *P*(data|start passage) tends to overestimate the actual start passage, if several neighboring start passages could have a similar ln *P*(data|start passage). In such a situation, a smaller start passage is chosen to avoid over-estimation if its ln P(data|start passage) is not less than the maximum minus a given threshold (set to be 0.2). In simulations, the inferred start passage numbers agree with the real start passages (Supplementary Table 4). In the inference of the real data, we only use mutations that are not classified as mutation with flat trajectory or “other” subtype by any start passages of 1 to 8 in the inference process, since they would not offer effective information about the actual start passage. The start passage in real data is inferred as 5.

Since the probabilistic method is more accurate than the neutral mutation method, and is applicable even to mutations with low mutation rate, it would be reasonable to consider 5 as the start passage, which is also very close to 4.26 predicted by the later.

### Simulation setup

The fitness of a genome is given by 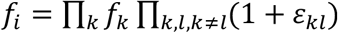, where *f_k_* and *ε_kl_* are, respectively, the fitness of a mutation at locus *k* and epistasis between mutations at loci *k* and *l*. We assume that the number of offspring of a genome is Poisson distributed around its fitness. Any locus of an offspring may mutate during replication according to locus-specific mutation rates. The population is diluted to its initial size (10^6^) by random sampling at end of every generation (5×10^6^). To introduce different levels of clonal interference effects into simulation, the number of beneficial mutations with large fitness is specified from 1 to 5. The performance of our model selection procedure in simulations is reported in Supplementary Figs. 18-19 and Supplementary Tables 8-13.

For numerical investigations, we find their deterministic numerical approximations based on the ordinary differential equations of the one-locus or two-loci models.

### Ambiguity in assessment of mutation interactions

Exact prediction of pairs of interacting mutation from frequencies data would be a challenging task. Different genetic interactions could lead to similar trajectories (Supplementary Fig. 9).

However, it is possible to assume that, in the case of a detrimental mutation subordinated to a beneficial mutation, the roster of candidate “dominant” mutations can be reduced to a small number of high frequency ones. Assume, for example, that a deleterious mutation stays detrimental or neutral even after epistatically interacting with a beneficial mutation. By calculation, if a beneficial mutation cannot reach an abundance over 0.025 in the late passages, this detrimental mutation is unable to have an abundance twice higher than its original mutation-selection balance (0.10 for fifth times higher than the original balance). In the data, there are 9 mutations with abundance higher than 2.5% at the last passage, 7 out of which is larger than 10%.

## References

1. Burger, R. Mathematical properties of mutation-selection models. Genetica 102/103, 279–298 (1998).

2. Rouzine, I.M., Rodrigo, A. & Coffin, J.M. Transition between stochastic evolution and deterministic evolution in the presence of selection: general theory and application to virology. Microbiol Mol Biol Rev 65, 151–85 (2001).

3. Desai, M.M. & Fisher, D.S. Beneficial mutation selection balance and the effect of linkage on positive selection. Genetics 176, 1759–98 (2007).

4. Goyal, S. et al. Dynamic mutation-selection balance as an evolutionary attractor. Genetics 191, 1309–19 (2012).

5. Nielsen, R. & Slatkin, M. An introduction to population genetics: theory and applications, (Sinauer Associates Sunderland, MA, 2013).

6. Hallatschek, O., Hersen, P., Ramanathan, S. & Nelson, D.R. Genetic drift at expanding frontiers promotes gene segregation. Proc Natl Acad Sci U S A 104, 19926–30 (2007).

7. Ewens, W.J. Mathematical Population Genetics 1: Theoretical Introduction, (Springer Science & Business Media, 2012).

8. Chou, H.H., Chiu, H.C., Delaney, N.F., Segre, D. & Marx, C.J. Diminishing returns epistasis among beneficial mutations decelerates adaptation. Science 332, 1190–2 (2011).

9. Khan, A.I., Dinh, D.M., Schneider, D., Lenski, R.E. & Cooper, T.F. Negative epistasis between beneficial mutations in an evolving bacterial population. Science 332, 1193–6 (2011).

10. Woods, R.J. et al. Second-order selection for evolvability in a large Escherichia coli population. Science 331, 1433–6 (2011).

11. Weinreich, D.M., Delaney, N.F., DePristo, M.A. & Hartl, D.L. Darwinian evolution can follow only very few mutational paths to fitter proteins. science 312, 111–114 (2006).

12. Good, B.H. & Desai, M.M. The impact of macroscopic epistasis on long-term evolutionary dynamics. Genetics 199, 177–90 (2015).

13. Szendro, I.G., Schenk, M.F., Franke, J., Krug, J. & De Visser, J.A.G. Quantitative analyses of empirical fitness landscapes. J Stat Mech Theory Exp 2013, P01005 (2013).

14. Gerrish, P.J. & Lenski, R.E. The fate of competing beneficial mutations in an asexual population. Genetica 102-103, 127–44 (1998).

15. Park, S.C. & Krug, J. Clonal interference in large populations. Proc Natl Acad Sci U S A 104, 18135–40 (2007).

16. Levy, S.F. et al. Quantitative evolutionary dynamics using high-resolution lineage tracking. Nature 519, 181–6 (2015).

17. Lang, G.I. et al. Pervasive genetic hitchhiking and clonal interference in forty evolving yeast populations. Nature 500, 571–4 (2013).

18. Maddamsetti, R., Lenski, R.E. & Barrick, J.E. Adaptation, Clonal Interference, and Frequency-Dependent Interactions in a Long-Term Evolution Experiment with Escherichia coli. Genetics 200, 619–31 (2015).

19. Lang, G.I., Botstein, D. & Desai, M.M. Genetic variation and the fate of beneficial mutations in asexual populations. Genetics 188, 647–61 (2011).

20. Kao, K.C. & Sherlock, G. Molecular characterization of clonal interference during adaptive evolution in asexual populations of Saccharomyces cerevisiae. Nat Genet 40, 1499–504 (2008).

21. Miralles, R., Gerrish, P.J., Moya, A. & Elena, S.F. Clonal interference and the evolution of RNA viruses. Science 285, 1745–7 (1999).

22. Strelkowa, N. & Lassig, M. Clonal interference in the evolution of influenza. Genetics 192, 671–82 (2012).

23. Smith, J.M. & Haigh, J. The hitch-hiking effect of a favourable gene. Genet Res 23, 23–35 (1974).

24. Hegreness, M., Shoresh, N., Hartl, D. & Kishony, R. An equivalence principle for the incorporation of favorable mutations in asexual populations. Science 311, 1615–7 (2006).

25. Luksza, M. & Lassig, M. A predictive fitness model for influenza. Nature 507, 57–61 (2014).

26. Wiser, M.J., Ribeck, N. & Lenski, R.E. Long-term dynamics of adaptation in asexual populations. Science 342, 1364–7 (2013).

27. Good, B.H., McDonald, M.J., Barrick, J.E., Lenski, R.E. & Desai, M.M. The dynamics of molecular evolution over 60,000 generations. Nature 551, 45–50 (2017).

28. Acevedo, A., Brodsky, L. & Andino, R. Mutational and fitness landscapes of an RNA virus revealed through population sequencing. Nature 505, 686–90 (2014).

29. Acevedo, A. & Andino, R. Library preparation for highly accurate population sequencing of RNA viruses. Nat Protoc 9, 1760–9 (2014).

30. Vignuzzi, M., Stone, J.K., Arnold, J.J., Cameron, C.E. & Andino, R. Quasispecies diversity determines pathogenesis through cooperative interactions in a viral population. Nature 439, 344–8 (2006).

31. Xiao, Y. et al. RNA Recombination Enhances Adaptability and Is Required for Virus Spread and Virulence. Cell Host Microbe 19, 493–503 (2016).

32. McDonald, M.J., Rice, D.P. & Desai, M.M. Sex speeds adaptation by altering the dynamics of molecular evolution. Nature 531, 233–6 (2016).

33. Rouzine, I.M., Wakeley, J. & Coffin, J.M. The solitary wave of asexual evolution. Proc Natl Acad Sci U S A 100, 587–92 (2003).

34. Schwarz, G. Estimating the dimension of a model. Ann Stat 6, 461–464 (1978).

35. Kass, R.E. & Raftery, A.E. Bayes factors. Journal of the american statistical association 90, 773–795 (1995).

36. Mani, R., St Onge, R.P., Hartman, J.L.t., Giaever, G. & Roth, F.P. Defining genetic interaction. Proc Natl Acad Sci U S A 105, 3461–6 (2008).

37. Raftery, A.E. Bayesian model selection in social research. Sociological methodology, 111–163 (1995).

