## Supplementary figures and tables for "Genetic interactions shaping evolutionary trajectories in an RNA virus population"

**Chang Chang[1†‡], Simone Bianco[2†], Ashley Acevedo[3†], Chao Tang[1*], Raul Andino[3*]**

[1]Center for Quantitative Biology, Schools of Physics and Peking-Tsinghua Center for Life Sciences, Peking University, Beijing 100871, China

[2]Industrial and Applied Genomics, IBM Accelerated Discovery Laboratory, IBM Research – Almaden, 650 Harry Road, San Jose, CA 95120-6099, USA

[3]Department of Microbiology and Immunology, University of California, San Francisco, San Francisco, CA 94158, USA

[†] Equal contribution authors

[‡] Present address: Department of Molecular and Cellular Biology, Harvard John A. Paulson School of Engineering and Applied Sciences, Harvard University, Cambridge, MA 02138, USA

**Supplementary figure and table legends**

**Supplementary Figure 1. Genetic compositions of serially passaged poliovirus populations were accurately sequenced using CirSeq. a,** Schematic of CirSeq sequencing method[28]. **b,** CirSeq is able to accurately measure the genetic composition of an RNA virus. **c,** Serial passage experiments were performed for poliovirus, for a total of 37 passages being sequenced for population genetic composition.

**Supplementary Figure 2. Mathematical model of population mutation dynamics. a,** Model settings. The asterisk stands for any of the two possible alleles: wild type, *w*, or mutant, *m*. **b,** Definitions of total frequency and average relative fitness of wild type and mutant alleles. **c,** A whole genome model (full model) requires a large number of parameters. Model reduction leads to simpler one-locus and two-loci models that are able to connect to the data, which enables extraction of information from mutation trajectories. In the absence of interaction between mutations, the full model reduces to the one-locus model. If epistasis, clonal interference or hitchhiking is introduced, the full model can be reduced to the two-loci model. Notably, epistatic interactions are usually asymmetric. For example, a “dominant” mutation with higher abundance at locus *a* affects the trajectory of the “subordinate” mutation with lower abundance at locus *b*, but usually not *vice versa*. Since back mutation is neglected in our analysis, the number of parameters for mutation rate are equal to those for fitness. The interaction parameter in two-loci model refers to the effect of epistasis, clonal interference, or hitchhiking.

**Supplementary Figure 3. Numerical trajectories in the context of a two-loci model.** All simulation settings are the same as in Fig. 2. **a,** Numerical trajectories. **b,** The table of fitness. Fitness of single and double mutants are set in the simulation. $f_{b|A}$ and $f_{ab|a}$ are important parameters of the inference for studying mutational trajectories (Methods). In this example, since there is no epistasis, the $f_{ab|a}$ of mutant *b* is still 0.8, which is identical to $f_{b|A}$. If there is positive epistasis, $f_{ab|a} > f_{b|A}$; if it is negative epistasis, $f_{ab|a} < f_{b|A}$. See Supplementary Figs. 4-6 for examples.

**Supplementary Figure 4. Evolutionary determinants of the frequency of a detrimental mutation in the context of two-loci model.** Numerical investigation of the dynamics of a detrimental mutation in a two-loci genome model. The fitness of the dominant mutation is 1.5 and the fitness of the subordinate mutation is 0.8. The mutation rate is $10^{-4}$ unless otherwise mentioned. Arrows are color coded to the curve changing under the considered effect. In the absence of interactions, detrimental mutations follow a one-locus detrimental model, asymptotically reaching mutation-selection balance (top row, left to right). When clonal interference is present but the double mutation can be established (delay=20), double mutants (the purple curve) rescues the frequency of detrimental mutation after its frequency is starting to decline. The time at which this dip is observed corresponds to the delay in establishing the double mutation in the population. When hitchhiking is present ($[ab]_0 = 10^{-5}$), the curve for double mutant moves upward, leading to another deformation of the trajectory of total abundance. Both positive epistasis (I: 0.8 to 0.95; II: 0.8 to 1; III: 0.8 to 1.05) and negative epistasis coefficients (0.8 to 0.5) can lead to a trajectory shifting of this mutation from its original mutation-selection balance.

**Supplementary Figure 5. Evolutionary determinants of the frequency of a neutral mutation in the context of two-loci model.** Numerical investigation of the dynamics of a neutral mutation in a two-loci genome model. Fitness of the dominant mutation is 1.5, while the subordinated mutation has fitness 1. The mutation rate is $10^{-4}$ unless otherwise mentioned. Arrows are color coded to the curve changing under the considered effect. In the absence of interactions, the frequency of neutral mutations would increase linearly in a very long-time scale, as predicted by the one-locus neutral mutation model. Different mutation rate ($10^{-5}$) would yield a different increasing slope. When clonal interference is present (delay=20, clonal interference I), the curve for double mutant (purple) moves to the right, leading to a dip in the middle part of the trajectory of total abundance. In the limiting case of infinite delay, the mutation may go extinct (cyan/green) (using delay=50 as an example, clonal interference II). When hitchhiking is present (I: $[ab]_0 = 10^{-5}$ and II: $[ab]_0 = 10^{-3}$), the curve for double mutant moves to the upper side, and the abundance of this neutral mutation could be very large in some parameter ranges. Both positive epistasis (1 to 1.1) and negative epistasis (1 to 0.9) can deform the trajectory of this mutation in the later phase.

**Supplementary Figure 6. Evolutionary determinants of the frequency of a beneficial mutation in the context of two-loci model.** Numerical investigation of the dynamics of a beneficial mutation in a two-loci genome model. Fitness of the dominant mutation is 1.5, that of the subordinate mutation is 1.2, and mutation rate is $10^{-4}$ unless otherwise mentioned. The color of an arrow is same as the curve that a factor could affect in a panel. Under the ordinary situation, this mutation would follow the trajectory predicted by the one-locus beneficial model (part of the trajectory would be linear in the log space). Different mutation rate ($10^{-5}$) would moves the trajectory. When clonal interference is present (delay=10 and 20, clonal interference I and II), the curve for double mutant (purple) moves to the right, leading to deformation in the middle part of the trajectory of total abundance (could be a dip). In the limiting case of infinite delay, the mutation can even extinct (using delay=50 as an example, clonal interference III). When hitchhiking is present ($[ab]_0 = 10^{-5}$), the curve for double mutant moves to the upper side, and this mutation could be fixed faster. Both positive epistasis (1.2 to 1.1) and negative epistasis (1.2 to 1.5) can deform the trajectory of this mutation in the later phase, leading to an earlier or a later fixation of this mutation.

**Supplementary Figure 7. Mutation(s) need to be at high abundance to affect the trajectory of an interacting mutation. a,** An example for two-loci epistasis. The initial abundance of the background beneficial mutation is set to be $10^{-3}$ (low abundance) and $10^{-1}$ (high abundance) in the left and right panels, respectively, with all other simulation settings being same in both panels. The fitness of this background beneficial mutation is 1.1. For another mutation, two separate numerical simulations are performed, one is without epistasis (fitness: 0.8), and another has epistasis with the background beneficial mutation ($f_{b|A}$: 0.8; $f_{ab|a}$: 0.95). For both mutations, the mutation rates are set to be $10^{-4}$. Left panel: When the abundance of the background mutation is low, the effect on its interacting mutation's trajectory is also low. Right panel: When the abundance of the background mutation is high, the effect on its interacting mutation's trajectory is not negligible. **b,** An example for three-loci epistasis. The initial abundance of the background beneficial mutation *1* is set to be 0.5. The initial abundance of the background beneficial mutation *2* is set to be $10^{-3}$ (low abundance) and $10^{-1}$ (high abundance) in the left and right panels, respectively, with all other simulation settings being same in both panels. The fitness of these two background beneficial mutations are both 1.1. For the third mutation, two separate numerical simulations are performed, one is without epistasis (fitness: 0.8), and another has three-loci epistasis with background beneficial mutations *1* and *2* ($f_{b|A}$: 0.8; $f_{ab|a}$: 0.95). For all mutations, the mutation rates are set to be $10^{-4}$. Left panel: When not all the background mutations have high enough abundance, the effect on their interacting mutation's trajectory is low. Right panel: When all the background mutations have high enough abundance, the effect on its interacting mutation's trajectory is not negligible. **a,b,** For a background beneficial mutation in any plot, its interacting mutation is of quite low abundance and only has negligible effect on their trajectories. Thus, in each pair of simulations (with and without epistasis), the trajectories of the same background beneficial mutation overlap well. In general, to affect the trajectory of an interacting mutation in a *N*-loci interaction, *N-1* mutations need to be of high abundance.

**Supplementary Figure 8. The comparison of trajectories of one-locus model and two-loci model without epistasis. a,b,c,d,** In each panel, the number in the parentheses is the selection coefficient of this mutation. Blue dots represent the numerical solution from the one-locus model, and the black dots are the numerical solution from the two-loci model. Examples of **a,** detrimental mutation, **b,** neutral mutation (left panel: log space; right panel: linear space), and **c,** beneficial mutation. **a,b,c,** In the two-loci model, the fitness of the interacting mutation is set to be 1.5. No epistasis, clonal interference, or hitchhiking is included. The curve of the two-loci model for the detrimental mutation overlaps extremely well with that of the one-locus model, but these of neutral or beneficial mutations differ from the trajectories of their one-locus model because of non-negligible higher-order terms of the hitchhiking effect from a beneficial mutation with larger fitness. **d,** On the other hand, the hitchhiking effect is much small for a mutation with smaller fitness to the trajectory of a mutation with larger fitness. The fitness of the interacting mutation in both panels is set to be 1.2. The plot is in linear space to make the differences between the curves easier to be observed. The hitchhiking effect to the mutation with fitness 1.5 is much smaller than that to the mutation with fitness 1.3, suggesting mutation with larger fitness is less affected.

**Supplementary Figure 9. Hard to determine the interacting partner of a detrimental mutation with epistasis.** Each panel corresponds to one simulation. In all simulations, the fitness of mutation m1-m3 are 1.2 (black), 1.1 (blue), and 0.8 (red), and all mutation rates are set to be $10^{-4}$. Epitasis is given in the title of each panel, with the subscripts indicating the interacting loci. Multiple combinations of epistasis in three-loci numerical simulations can lead to similar trajectory shifting for a detrimental mutation with epistasis, thus it is hard to determine this mutation's interacting partner from its trajectory. See the left upper panel vs. the right upper panel as an example. Epistasis between m3 and m1 is 0.25 in the left upper panel, and that between m3 and m2 is 0.3, both of which lead to similar trajectories for mutation m3. Thus, not only it is hard to determine which mutation is interacting with mutation m3, but also we do not know the actual epistasis effect. We can infer the "effective" epistasis effect in the context of the two-loci model.

**Supplementary Figure 10. Some epistasis effects related to beneficial mutations can be masked by clonal interference or hitchhiking. a,** Examples of beneficial mutation trajectories that are vulnerable to clonal interference. In each panel, the numbers in the parentheses are the selection coefficients of this mutation (a mutation with a pair of number is with epistasis, otherwise it is without epistasis). The black line represents the numerical solution using the selection coefficients specified in the title (thus is without clonal interference), and the blue dots are an example trajectory in simulation that is under the effect of clonal interference (simulation genome length = 20; simulation settings are the same to that in Supplementary Table 11). All mutation rates are set to be $10^{-4}$. (Upper panels) Two examples of beneficial-to-beneficial mutation with positive epistasis. (Lower left panel) One example of beneficial mutation. (Lower right panel) An example of beneficial-to-beneficial mutation with negative epistasis. **b,** Trajectories of beneficial mutations with epistasis can be similar to these without epistasis. In each panel, the black dots are the numerical solution specified by given selection coefficients (with epistasis), and the purple curve gives the best fitting model (for all examples in this panel, the best models are without epistasis, and their names are shown in title), suggesting that the trajectories of some beneficial mutations with epistasis can be similar to mutations without epistasis but with a different set of parameters. **c,** Trajectories of beneficial mutations without epistasis can be similar to these with epistasis. The left panel gives the numerical solution of a beneficial mutation with fitness 1.05 (black line) and an example of trajectory that has deformation in simulation (blue dots), and the right panel gives the best fitting model (the purple curve) to this example trajectory. **b,c,** The example in the upper right panel of panel *b* is classified as beneficial mutation under clonal interference, but its *delay* parameter is unable to be estimated. The reason is that the best fitting model here is actually beneficial-to-beneficial two-loci model. However, as suggested by panel *c*, trajectories of beneficial mutations without epistasis can be similar to these with epistasis. To control the false positive rate of predicting epistasis, mutations that are best fit with beneficial-to-beneficial two-loci model, if do not meet additional constraints, will be classified as beneficial mutation under clonal interference.

**Supplementary Figure 11. Different visualization of some panels in Fig. 3. a,** Semi-log plot of the neutral mutation A2959G. Its linear plot is available in Fig. 3a. **b,** Linear plot of the neutral mutation under hitchhiking U673C. Its semi-log plot is available in Fig. 3b.

**Supplementary Figure 12. An example of model selection. a,** In this plot, different panels represent the fitting of different models to the trajectory of the mutation at position 4269 (A to G; Fig. 3c). The meaning of the codes in each subtitle is the same as these in Fig. 4a. **b,** Detailed descriptions for best models in each level of model complexity. Note that the BIC is calculated based on the fit in the log space. Thus, although the average $R^2$ of model B(I) is larger comparing to the model N(I), it does not have a lower BIC.

**Supplementary Figure 13. A mutation trajectory example that fits detrimental mutation under clonal interference best but also can be fitted by detrimental mutation with negative epistasis.** In this example (A2683G), the best model is detrimental mutation under clonal interference. Notably, detrimental-to-detrimental negative epistatic model can also fit its trajectory (both models' codes are marked with blue). The reason why this example was not classified as detrimental mutation with negative epistasis is because it does not meet the $\Delta BIC \geq 10$ criterion.

**Supplementary Figure 14. More similar examples to that in Supplementary Fig. 13.** More examples that fit detrimental mutation under clonal interference best but also can be fitted by detrimental mutation with negative epistasis are given. The models, i.e., cyan curves given in each plot, are detrimental mutation under clonal interference. Notably, some of the examples list here do not pass the $\Delta R^2 \geq 0.9$ criterion for the detrimental-to-detrimental negative epistatic model.

**Supplementary Figure 15. Characterizing the poliovirus population at locus level by distributions of major evolutionary factors. a,** The distribution of mutation rate. **b,** The distributions of $f_{b|A}$ (left panel) and **c,** $f_{ab|a}$ (right panel). **d,** The sign of epistasis in the population. Most mutations that can be tested using our model show no sign of epistatic interaction. A small but non-negligible fraction of mutations are subject to epistatic interactions. **e,** The distribution of delay caused by the effect of clonal interference. **f,** The distribution of initial frequency of $[ab|a]$ caused by the effect of hitchhiking. **a,b,c,d,e,f,** Only mutations with estimable fitness and other evolutionary factors are plotted. Thus, mutation with flat trajectory and mutations of "other" are not included.

**Supplementary Figure 16. Scatter plots between major evolution factors. a,** Scatter plots of the effect of epistasis versus $f_{b|A}$ (left panel) and $f_{ab|a}$ (right panel). **b,** Scatter plots of mutation rate versus $f_{b|A}$ (left panel) and $f_{ab|a}$ (right panel). **c,** Scatter plots of delay caused by the effect of clonal interference versus $f_{b|A}$ (left panel) and $f_{ab|a}$ (right panel). **d,** Scatter plots for initial frequency of $[ab\,|\,a]$ as caused by hitchhiking versus $f_{b|A}$ (left panel) and $f_{ab|a}$ (right panel). **e,** Scatter plots of mutation rate versus the effects of epistasis, clonal interference and hitchhiking.

**Supplementary Figure 17. Genomic positional distribution and enrichment analysis of**

**mutation subtypes. a,** Genomic positional distribution for each mutation subclass. Red represents mutations of beneficial subtypes, black is neutral mutation, and blue is for detrimental subtypes. Dot represents mutations without epistasis, and other sharps are for these with epistasis. Red cross is neutral-to-beneficial, and red square is beneficial-to-detrimental. Blue upper triangle, blue cross, blue star represent detrimental mutation with positive epistasis, compensated mutation, and detrimental-to-beneficial. **b,** Enrichment analysis for beneficial mutations, neutral mutations, and negative mutations in genomic coding regions, with their false discovery rate (FDR) being calculated. Those significant FDR values are marked with blue.

**Supplementary Figure 18. Performance of the model selection procedure in non-epistasis simulation for predicting epistasis.** Fitness and epistasis type are estimated from mutation trajectories. In all simulations, genome length is set to be 1000, and the per locus mutation rate is $10^{-4}$. The initial population size is set to $10^6$, and the population is allowed to grow until it reaches the saturation population size ($5 \times 10^6$). To investigate the effect of clonal interference on the model selection procedure for predicting epistasis, we set the number of beneficial mutations with large fitness (1.5) be 1 to 5. In each plot, the x-axis indicates the $f_{b|A}$, and the y-axis indicates the $f_{ab|a}$. Color represents the predicted mutation type: a black mutation is of no epistasis, a blue one is with positive epistasis, and a green one is with negative epistasis. The sharp of beneficial mutations with large fitness are set to be a cross, and that of all other mutations is point. Note that hitchhiking phenomenon could easily be observed for the neutral simulation, since the number of inferred beneficial mutations is larger than the actual number. False positive rates of the simulations, defined as the fraction of non-epistasis mutations being predicted with epistasis, are available at Supplementary Table 8a.

**Supplementary Figure 19. Performance of the model selection procedure in epistasis simulation for predicting epistasis.** Simulation settings are same as these in Supplementary Fig. 18, expect that the effect of epistasis is included. For each group of simulation, roughly half of the mutations are under negative epistasis, and another half are under positive epistasis. Simulations for beneficial mutations under epistasis or neutral mutation with positive epistasis would be greatly suffer from the effect of clonal interference, and are shown in Supplementary Table 11. True positive rates of the simulations, defined as the fraction of epistasis being correctly predicted**,** are available at Supplementary Table 8b.

**Supplementary Figure 20. Estimated delay of non-epistasis mutations caused by clonal interference in simulations. a,** These panels shown the distributions of estimated delay in same simulations of Supplementary Fig. 18. **b,** Average delay of each mutation type is given, with the number in the pair of parentheses being its standard deviation.

**Supplementary Figure 21. Estimated effects of hitchhiking of non-epistasis mutations in simulations. a,** These panels shown the distributions of estimated initial frequencies of $[ab|a]$ in same simulations of Supplementary Fig. 18. **b,** Average initial value of double mutant of each mutation type is given, with the number in the pair of parentheses being its standard deviation.

**Supplementary Table 1. Models in model selection.** One-locus and two-loci models obtained in model reduction are included in the model selection procedure. These models were classified into different levels according to their complexity. Notably, although the number of parameters for two-loci clonal interference or hitchhiking models are the same as these of the two-loci epistasis models, the former is prior the latter in the model selection procedure to guarantee the accuracy of our epistasis predictions. The numbers in 3rd to 5th columns for two-loci models are equation indices that would be used in Eq. (2).

**Supplementary Table 2. Parameter ranges in the model selection process.** In the model selection process, multiple rounds of fitting with different initial parameter values are performed, with one round begins with initial values shown in the table, and other rounds with random initial values (N=5). Notably, for clonal interference, the upper limit of delay in inference is larger than the number of passages we have in the experiments. If the inferred delay is a large number, it corresponds to the classical phenomenon of clonal interference (Fig. 2e, left panel), of which mutation(s) extinct, at least its frequency not increase again after the decrease in the time window of our experiments.

**Supplementary Table 3. Mutation subtype description**. The relationship between models listed in Supplementary Table 1 and the mutation subtypes listed here do not have one-to-one correspondence, different models could result in the same mutation subtype. For example, if a one-locus detrimental model fits a mutation trajectory well, it can lead to “detrimental; without epistasis”. However, in the case that a two-loci detrimental-to-detrimental epistatic model fits best, but the inferred values of $f_{b|A}$ and $f_{ab|a}$ of this model are very close, it can also lead to the same mutation subtype. In model selection, to improve the accuracy of predicting epistasis, the “establishment” mutation subtype serves as quality control for "detrimental; with positive epistasis" or "detrimental to beneficial". When either of above-mentioned subtypes fits well with a mutation, if the inferred initial mutation-selection balance of this mutation is smaller than a given threshold (set to be $10^{-5}$), such a mutation would be classified as "establishment" instead.

**Supplementary Table 4. Performance of estimating start passage number in simulation.** We employ two methods of estimating the start passage number, neutral-mutation-based method and probabilistic method, and evaluate them by simulation. In the simulation, the composition of the simulated genomes is complex. Just as the simulation in Supplementary Fig. 18, the genome length is set to be 1000, and the numbers of beneficial mutation of large fitness are range from 1 to 5 with mutation rates being $10^{-4}$. For each 100 loci, their fitness/epistasis are set to be 1, 0.99, 0.98, 0.95, 0.9, 1.01, 1.02, 1.05, 0.8 to 1, and 1 to 0.8, with their mutation rates being indicated in the table. The data before the actual start passage numbers (1-8) are removed and then being estimated by two methods. The neutral methods would unable to estimate the start passage when the mutation rate is low ($\leq 10^{-5}$), thus these parts are not shown in the table.

**Supplementary Table 5. The number of inferred cases for each mutation subclass with different start passage numbers.** The actual or effective start passage of the real data is unknown but is a global parameter for the mutation subclass inference. Using a neutral-mutation-based method and a probabilistic method (Methods), we determine the start passage as 5. In this table, we show the number of inferred case with different start passage numbers, and marked the inferred start passage number (5) with bold font.

**Supplementary Table 6. Mutation subtype counting according to mutation type.** Mutation types are listed according to the magnitude of their mutation rates (figure 3 in Acevedo *et al.*[28]).

**a,** Counting by major classifications. **b,** Counting for positive and negative epistasis. **c,** Detailed counting for non-epistasis subtypes of different fitness types, where *o* stands for the one-loci model of this fitness type, *ci* is the clonal interference version, and *h* refers to hitchhiking.

**Supplementary Table 7. Information of each mutation examples in Fig. 3.** The meanings of symbols in "Estimation" are the same as these in Fig. 3.

**Supplementary Table 8. Performance of the model selection procedure in simulation with mutation rate $10^{-4}$ for predicting epistasis.** This table shows the performance of predicting epistasis for simulations in Supplementary Figs. 18 and 19. False positive rate is defined as the fraction of non-epistasis mutations being predicted with epistasis, and true positive rate is defined as the fraction of mutation with epistasis being correctly predicted. **a,** False positive rate of simulations without epistasis. **b,** True positive rate of simulations with epistasis.

**Supplementary Table 9. Performance of the model selection procedure in simulation with mutation rate $10^{-5}$ for predicting epistasis.** All simulation settings are same with these in Supplementary Figs. 18 and 19 except the mutation rate. **a,** False positive rate of simulations without epistasis. **b,** True positive rate of simulations with epistasis.

**Supplementary Table 10. Performance of the model selection procedure in simulation with mutation rate $10^{-6}$ for predicting epistasis.** All simulation settings are same with these in Supplementary Figs. 18 and 19 except the mutation rate. **a,** False positive rate of simulations without epistasis. **b,** True positive rate of simulations with epistasis.

**Supplementary Table 11. Performance of the model selection procedure for beneficial mutations in simulation.** In the simulations, the genome length is set to be 20, the mutation rate is $10^{-4}$, and the numbers of beneficial mutation with large fitness vary (1-5). **a,** False positive of simulations without epistasis. **b,** True positive of simulations with epistasis. **a,b,** The number of mutation being inferred as epistasis is given before the slash, and the total mutation number is given after the slash. The simulation results suggest that even when the supply for beneficial mutation is only few tens, clonal interference could mask almost all epistasis related to beneficial mutation, if their $f_{b|A}$ and $f_{ab|a}$ are larger or equal than 1.

**Supplementary Table 12. Performance of the model selection procedure in simulation for estimating mutation rate.** This table shows the inferred mutation rates in same simulations of Supplementary Figs. 18 and 19.

**Supplementary Table 13. Performance of the model selection procedure in simulation for estimating $f_{b|A}$ and $f_{ab|a}$.** This table shows the inferred $f_{b|A}$ and $f_{ab|a}$ in same simulations of Supplementary Figs. 18 and 19.

**Supplementary Table 14. Counting of mutation with flat trajectory and "other" subtype in simulations.** This tables shows the fractions of mutation with flat trajectory and "other" subtype in three groups of simulations with varying mutation rates ($10^{-4}$: the simulation is same as that in Supplementary Fig. 18; $10^{-5}$: Supplementary Table 9; and $10^{-6}$: Supplementary Table 10). The mutation rates for all beneficial mutations with large fitness are still $10^{-4}$, but these of other mutations are given as indicated in this table. Multiple factors, including fitness, mutation rate, and clonal interference, could contribute to these fractions.

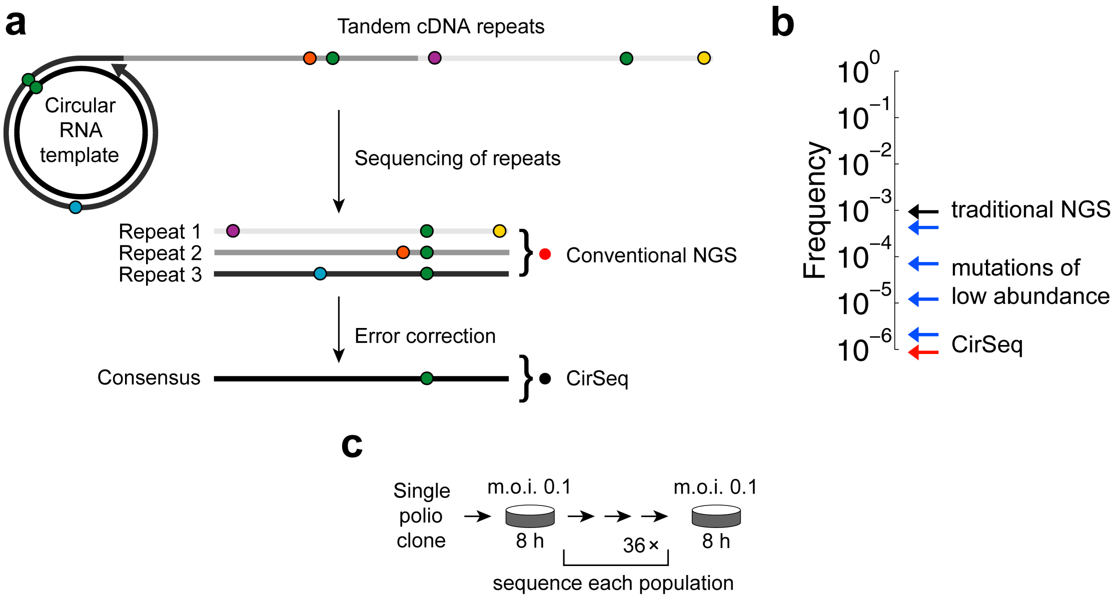


Supplementary Figure 1

**a**

| Abundance | Locus $k$ | Average relative fitness |
|---|---|---|
| $z_k(T)$ | ***w*** | $f^r_{z_k}(T)$ |
| | $\mu_k^- \; \uparrow\downarrow \; \mu_k^+$ | |
| $y_k(T)$ | ***m*** | $f^r_{y_k}(T)$ |

**b**

| Total abundance of 2nd locus | Abundance | Fitness | Average relative fitness |
|---|---|---|---|
| $z_2(T) = [\mathrm{ww}] + [\mathrm{mw}]$ | [ww] | $f_{ww}$ | $f^r_{z_2}(T) = \dfrac{[\mathrm{ww}] f_{ww} + [\mathrm{mw}] f_{mw}}{[\mathrm{ww}] + [\mathrm{mw}]}$ |
| | [mw] | $f_{mw}$ | |
| $y_2(T) = [\mathrm{wm}] + [\mathrm{mm}]$ | [wm] | $f_{wm}$ | $f^r_{y_2}(T) = \dfrac{[\mathrm{wm}] f_{wm} + [\mathrm{mm}] f_{mm}}{[\mathrm{wm}] + [\mathrm{mm}]}$ |
| | [mm] | $f_{mm}$ | |

**c**

| Model: name | Model: illustration | Parameter no.: fitness | Parameter no.: mutation | Parameter no.: interaction | Application |
|---|---|---|---|---|---|
| Full model | ***w*** $\mu_k^- \; \uparrow\downarrow \; \mu_k^+$ ***m*** | $2^L$ | 2L | | |
| One-locus models | w $\mu_k^- \; \uparrow\downarrow \; \mu_k^+$ m | 1 | 1 | 0 | Locus-independent<br>1. beneficial mutations<br>2. neutral mutations<br>3. detrimental mutations |
| Two-loci models | w $\mu_a^- \; \uparrow\downarrow \; \mu_a^+$ m w $\mu_b^- \; \uparrow\downarrow \; \mu_b^+$ m | 2 | 2 | 1 | Interactions between<br>1. beneficial-beneficial<br>2. beneficial-detrimental<br>~~3. detrimental-detrimental~~<br>Clonal interference<br>Hitchhiking |

Supplementary Figure 2

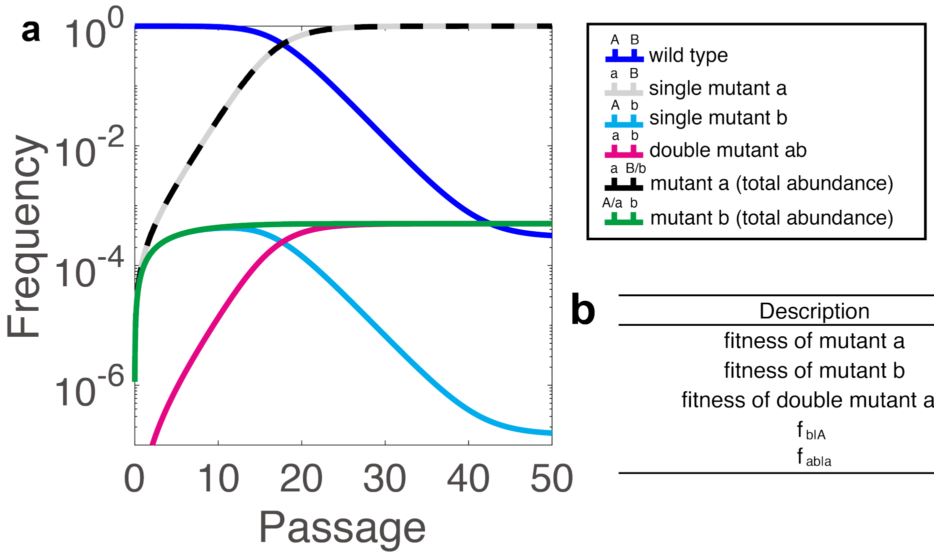


**b**

| Description | Fitness |
|---|---|
| fitness of mutant a | 1.5 |
| fitness of mutant b | 0.8 |
| fitness of double mutant ab | 1.2 |
| $f_{b\|A}$ | 0.8/1 |
| $f_{ab\|a}$ | 1.2/1.5 |

Supplementary Figure 3

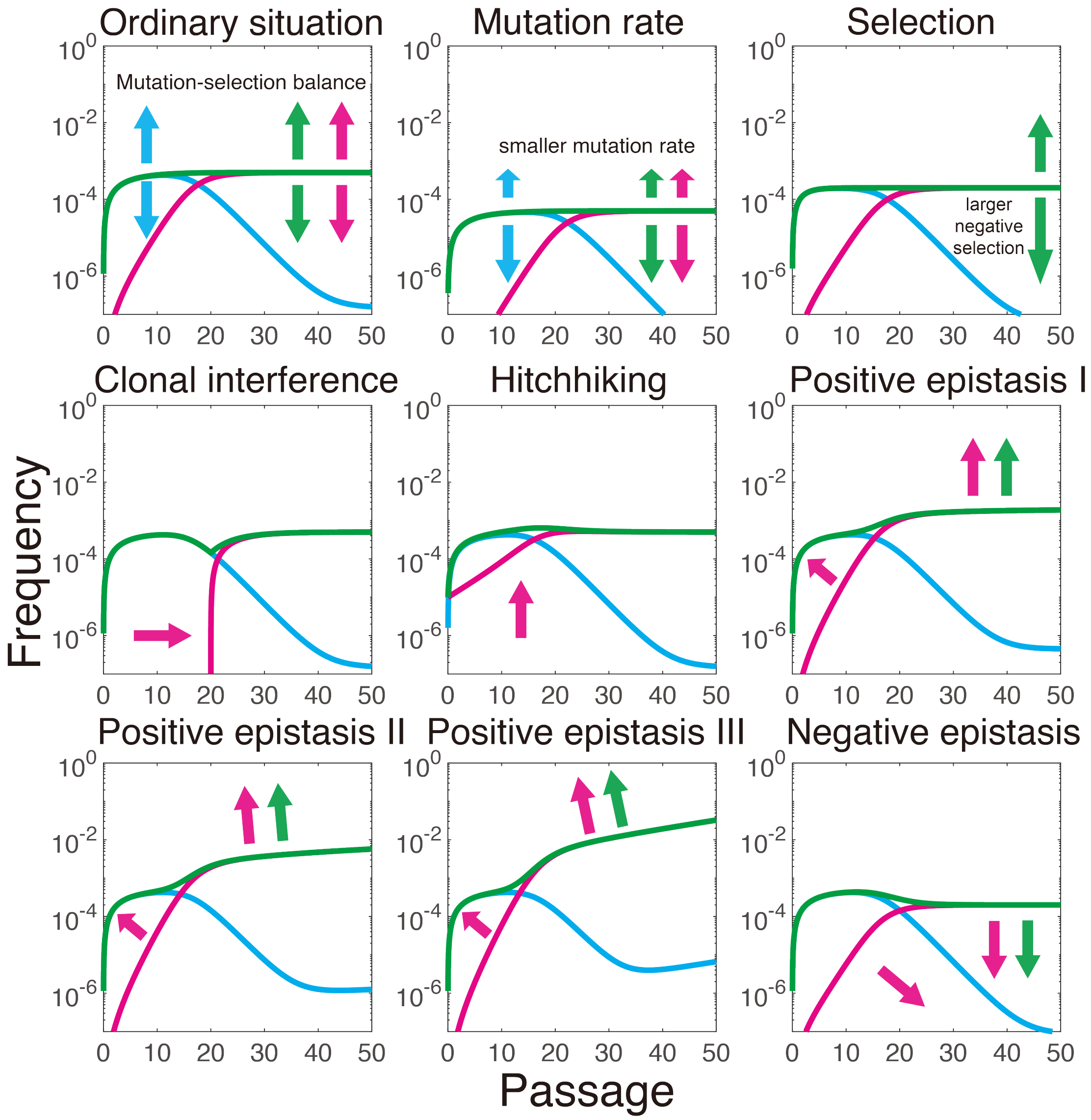


Supplementary Figure 4

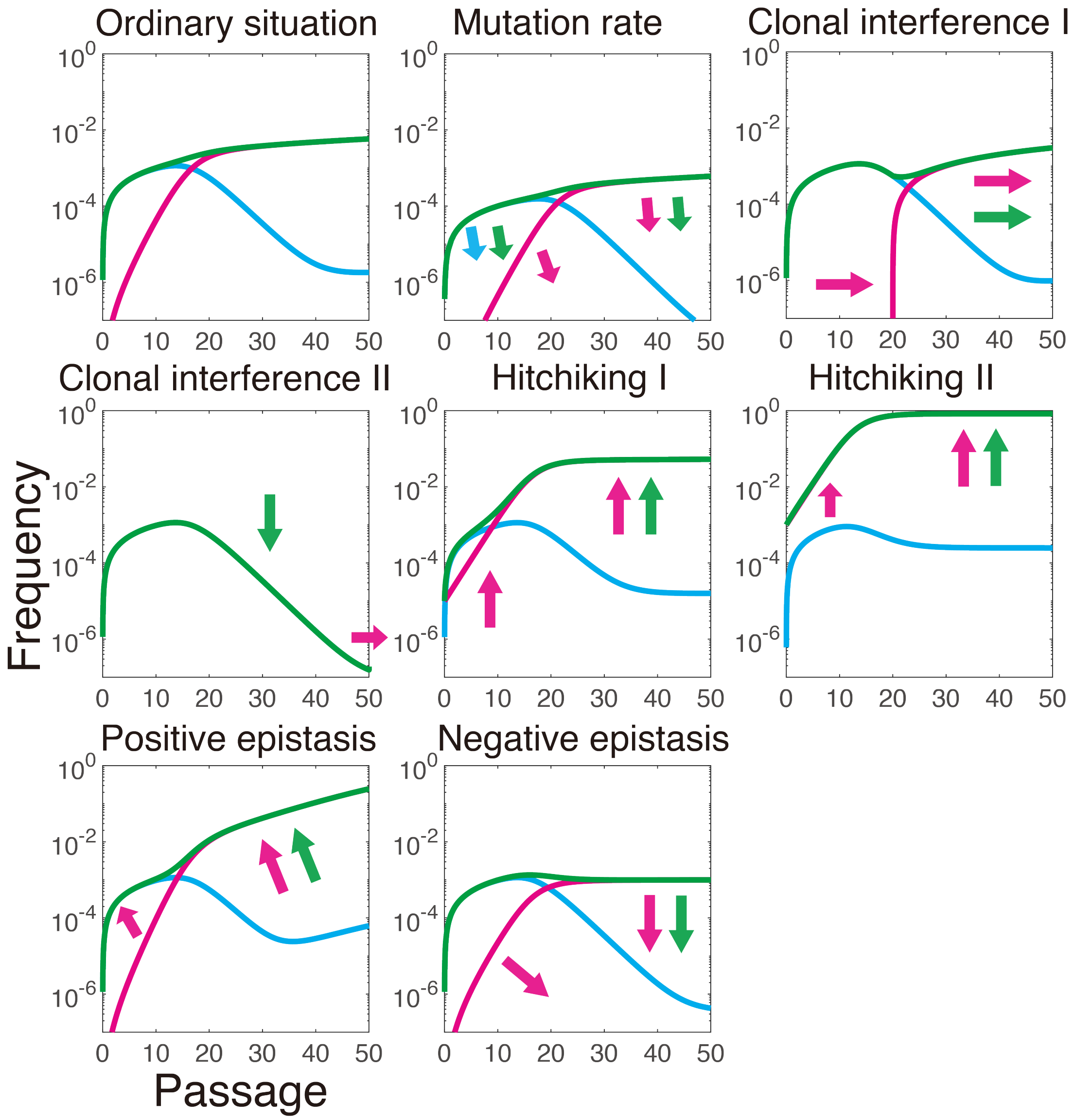


Supplementary Figure 5

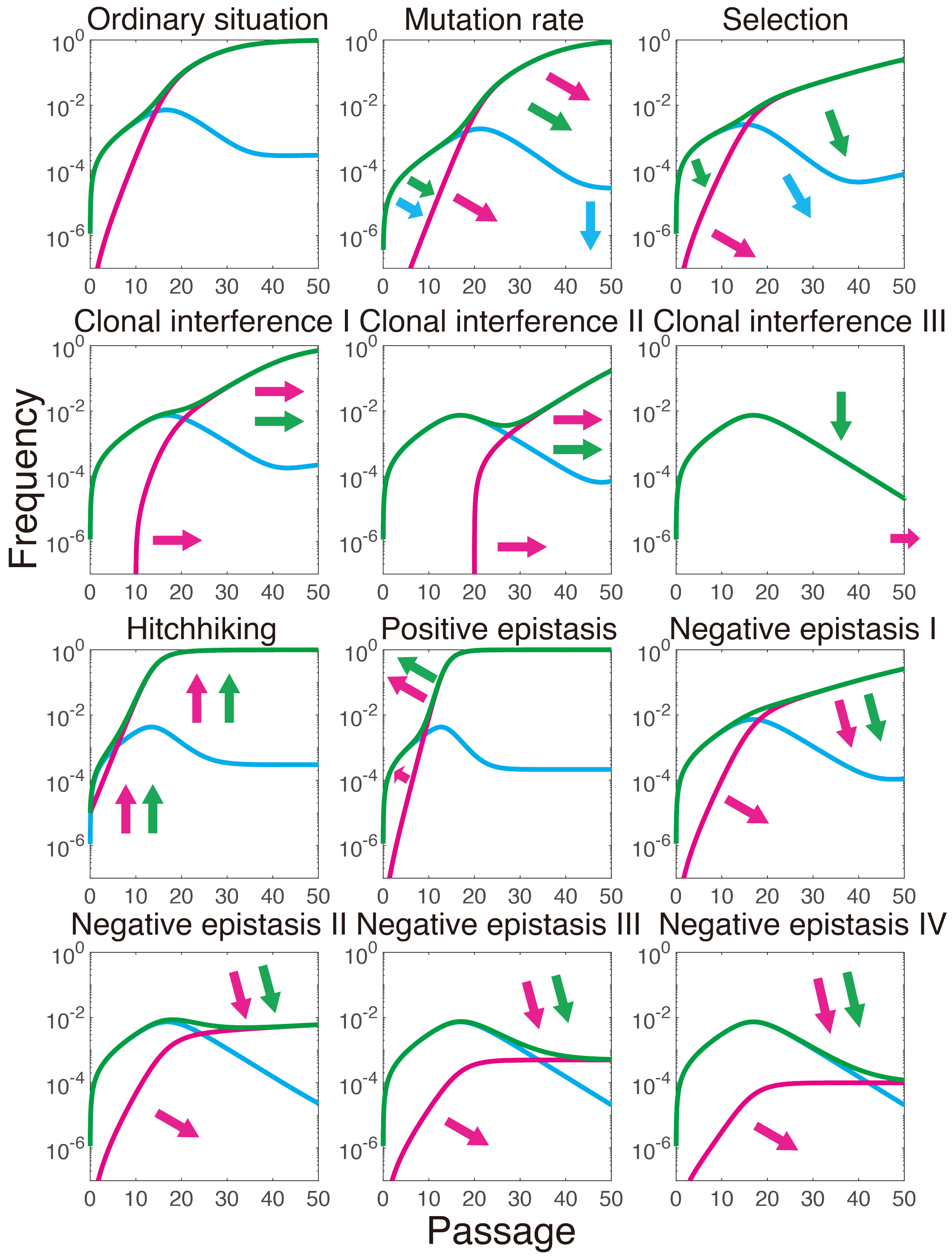


Supplementary Figure 6

**a**

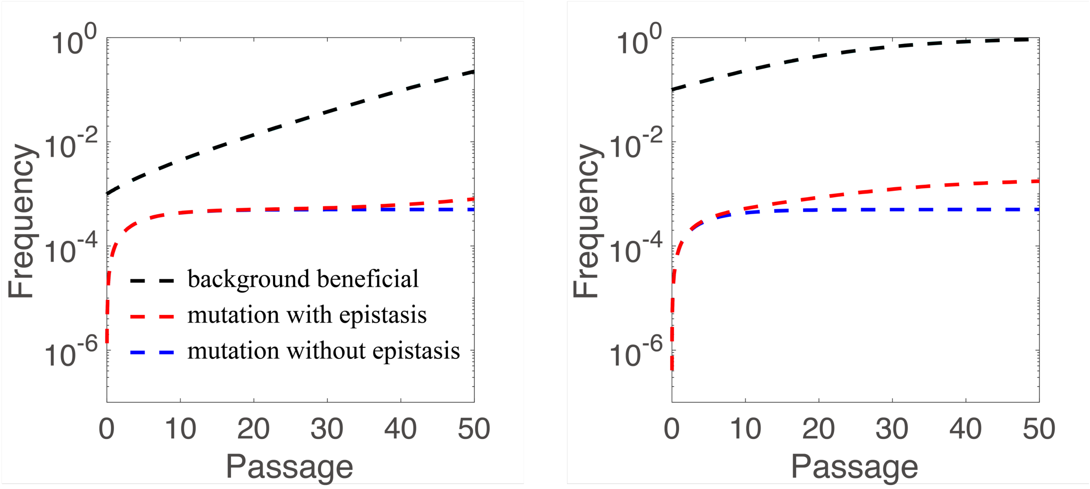


**b**

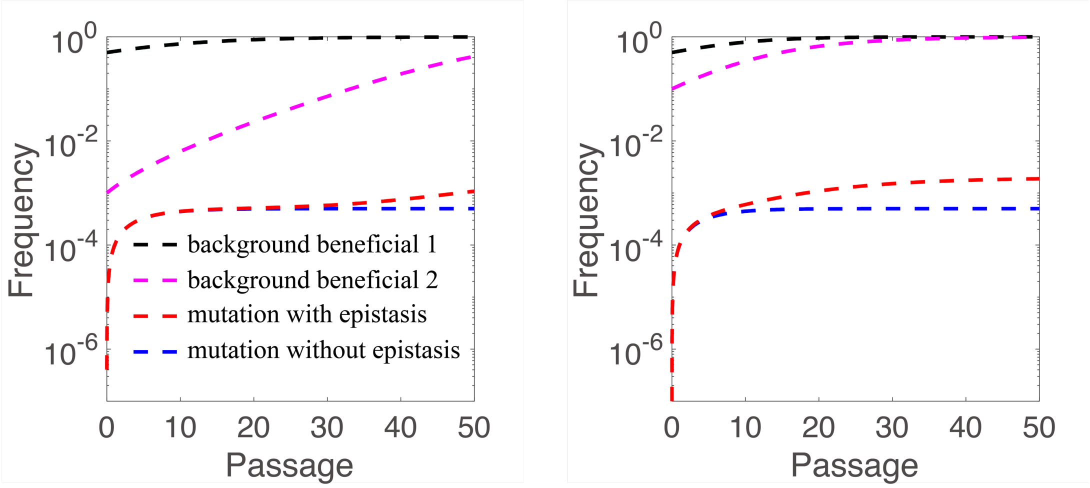


Supplementary Figure 7

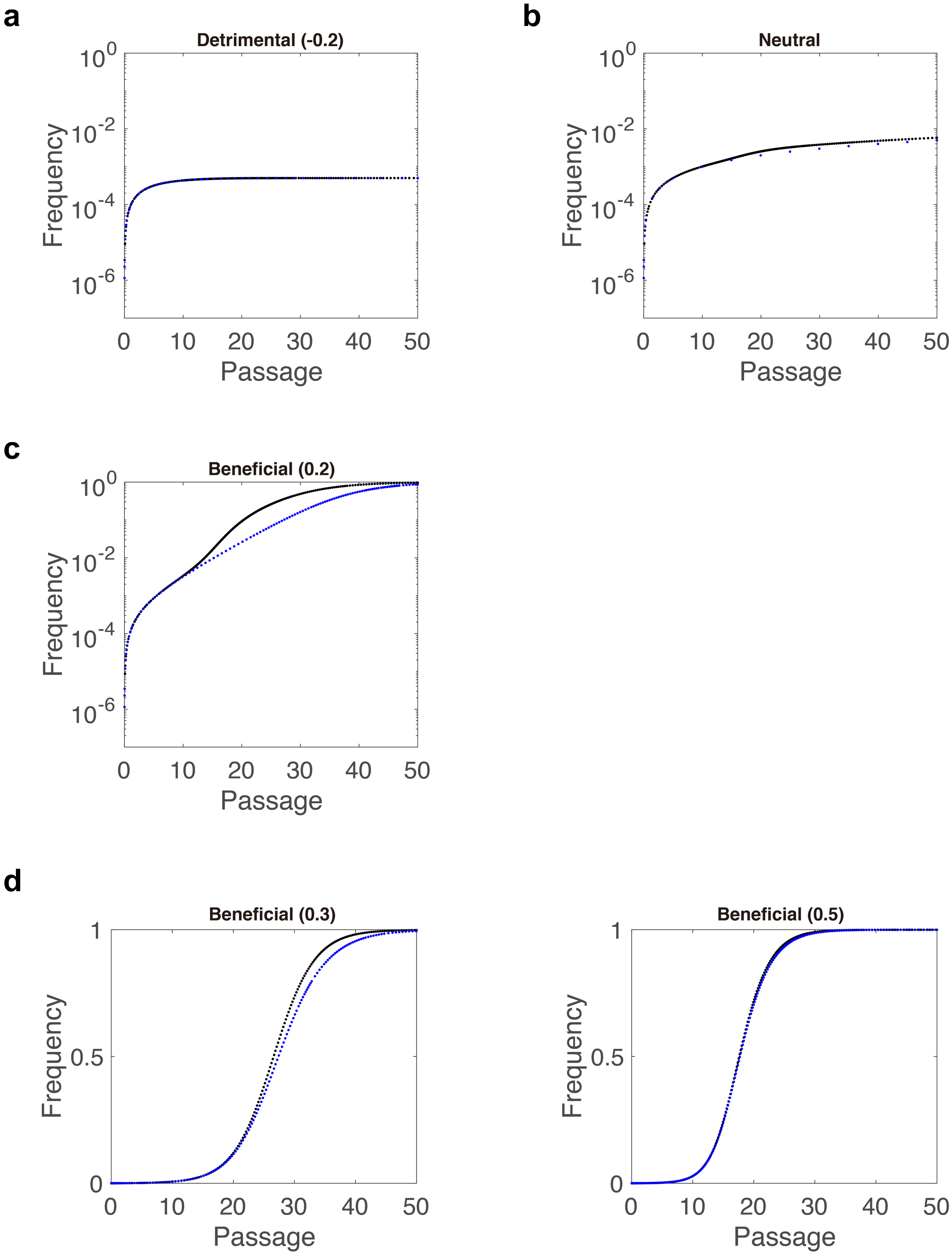


Supplementary Figure 8

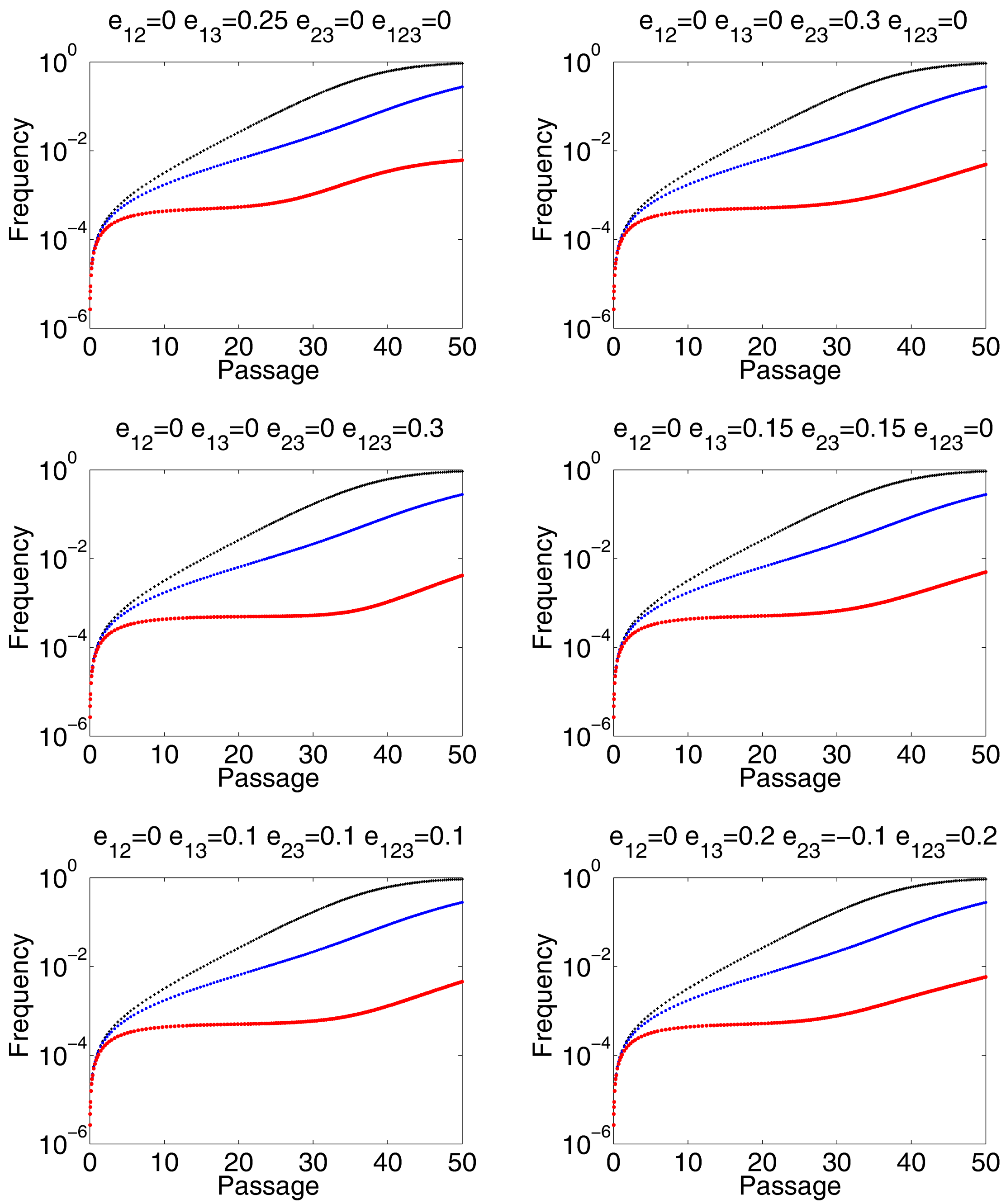


Supplementary Figure 9

**a** **Trajectories of beneficial mutation with or without epistasis are vulnerable to clonal interference**

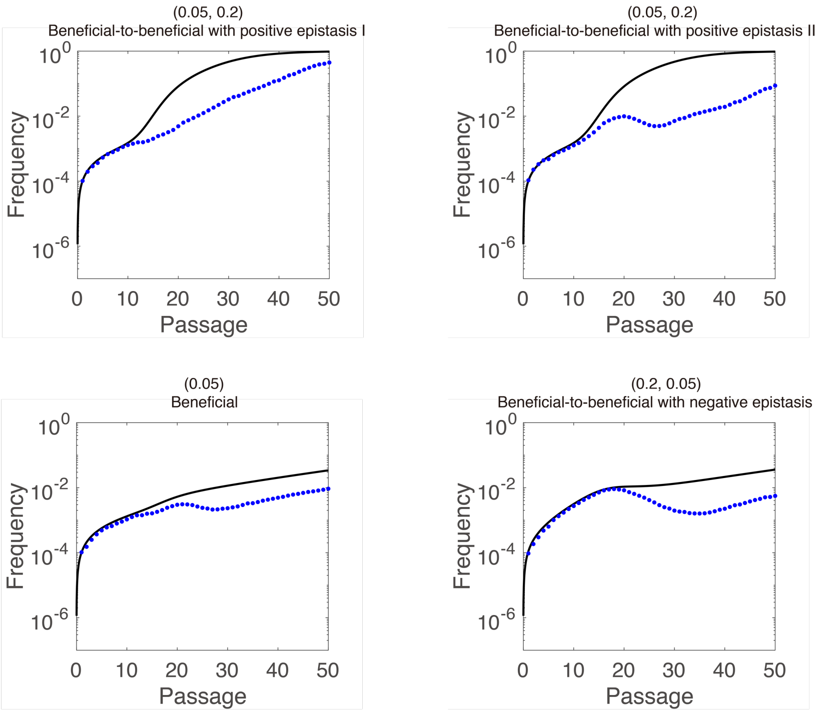


**b** **Trajectories of beneficial mutations with epistasis can be similar to these without epistasis**

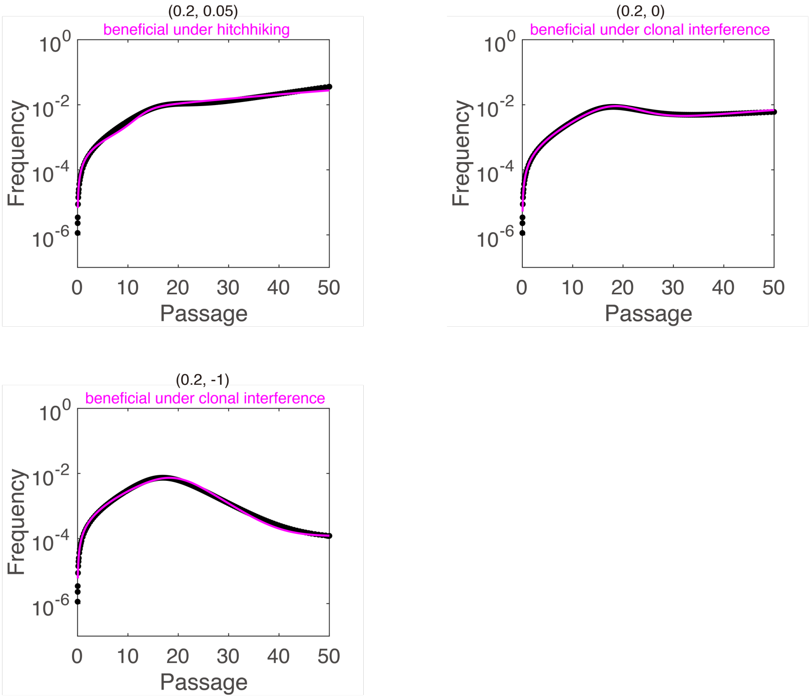


**c** **Trajectories of beneficial mutations without epistasis can be similar to these with epistasis**

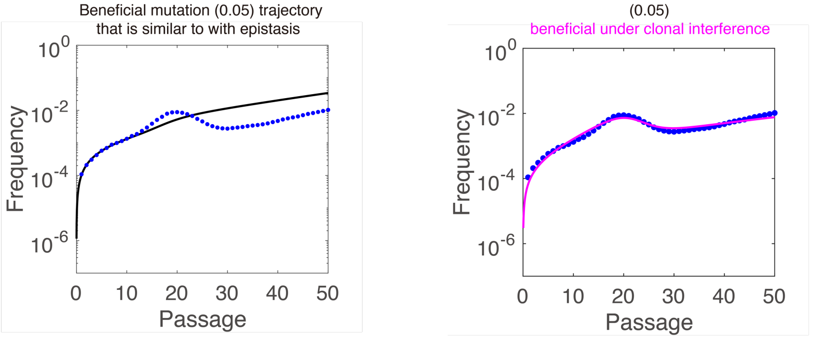

Supplementary Figure 10

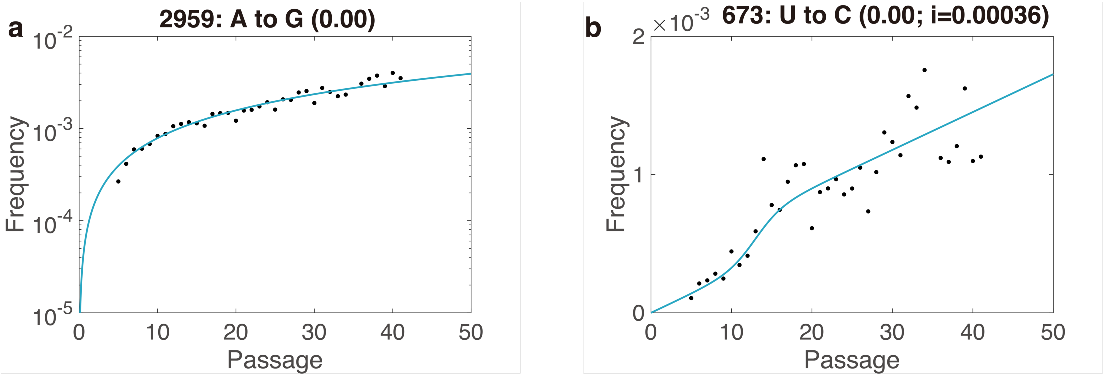


Supplementary Figure 11

a

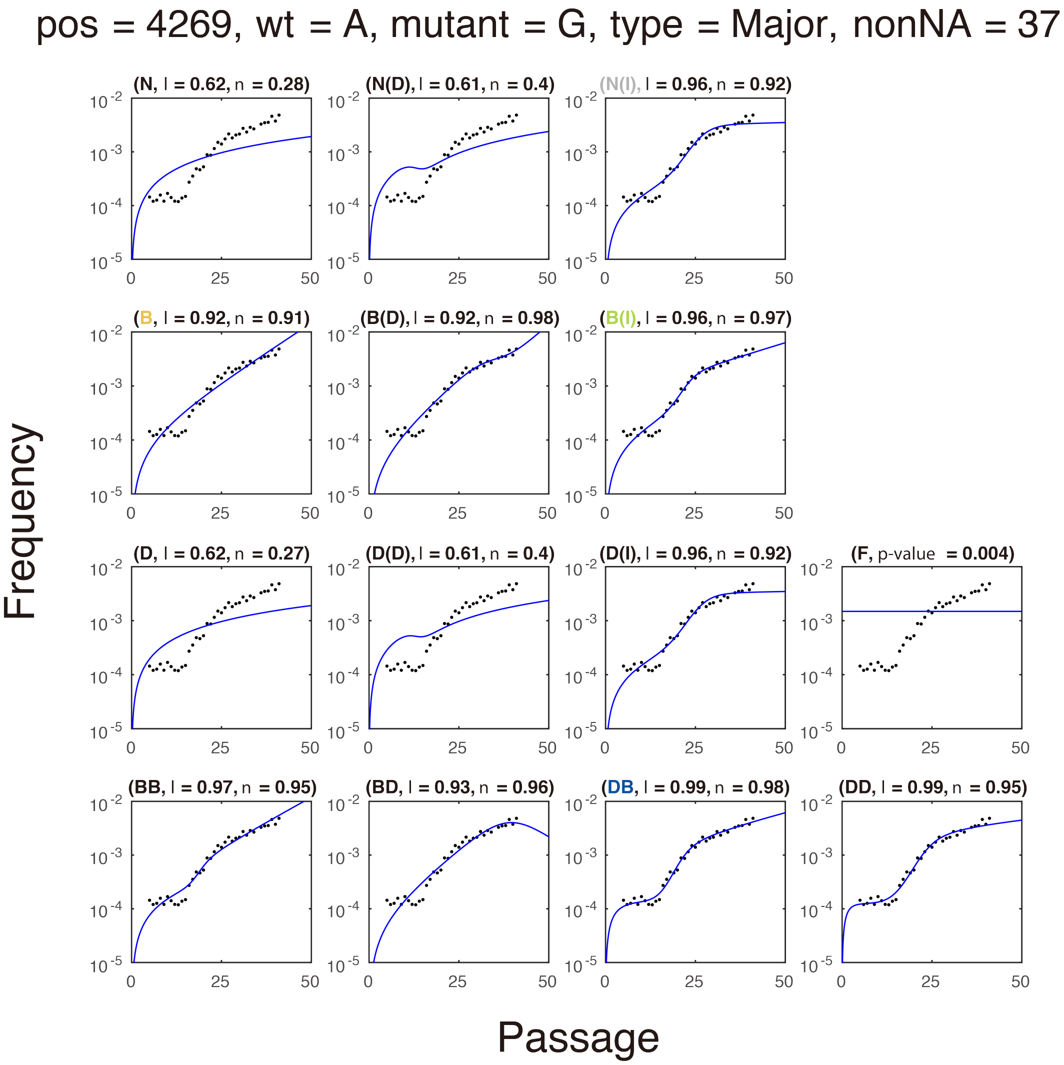


b

| Model level | Model | Estimation | average R2 | BIC |
|---|---|---|---|---|
| 2 | B | 0.10 | 0.92 | -65.82 |
| 3 | N(I) | i2=0.0028 | 0.94 | -81.29 |
| 4 | B(I) | 0.05 (i2=0.00045) | 0.97 | -77.83 |
| 5 | DB | -0.33 to 0.03 | 0.98 | -120.68 |

Supplementary Figure 12

pos = 2683, wt = A, mutant = G, type = Major, nonNA = 37

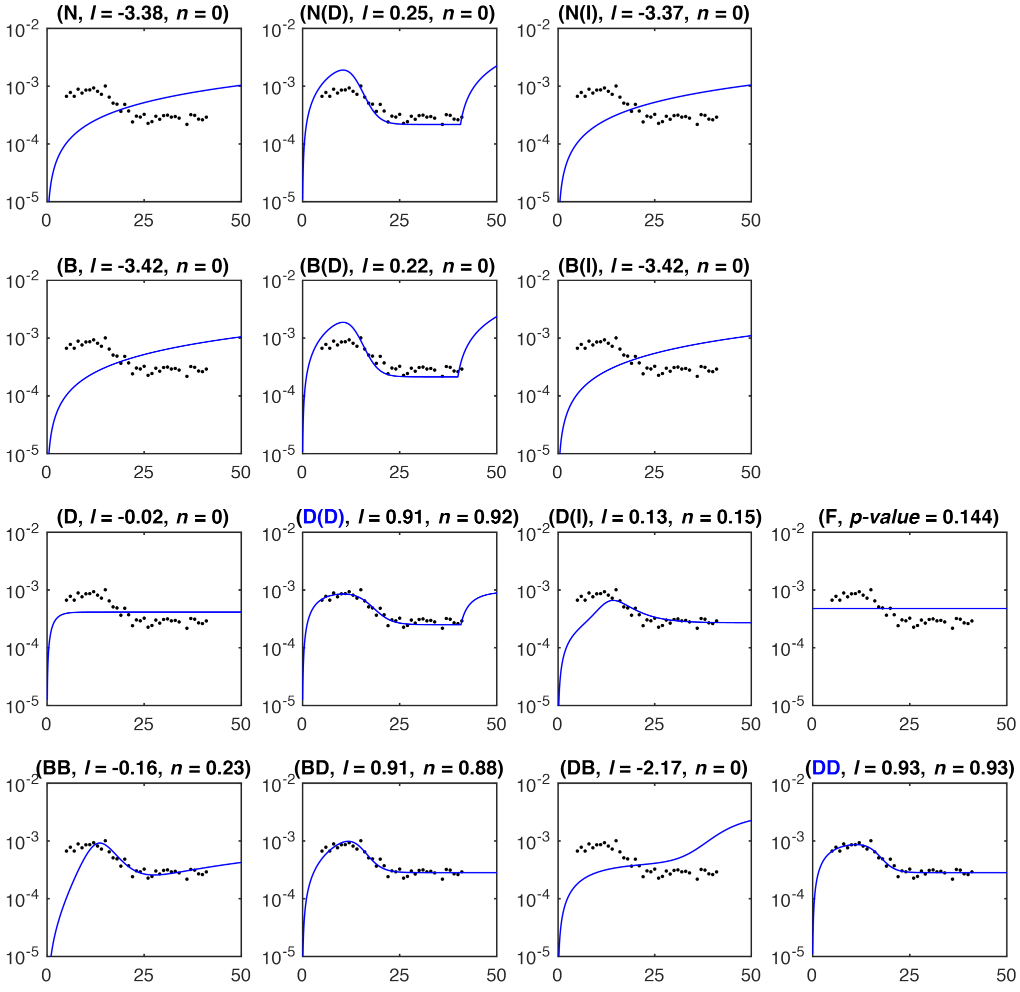


Supplementary Figure 13

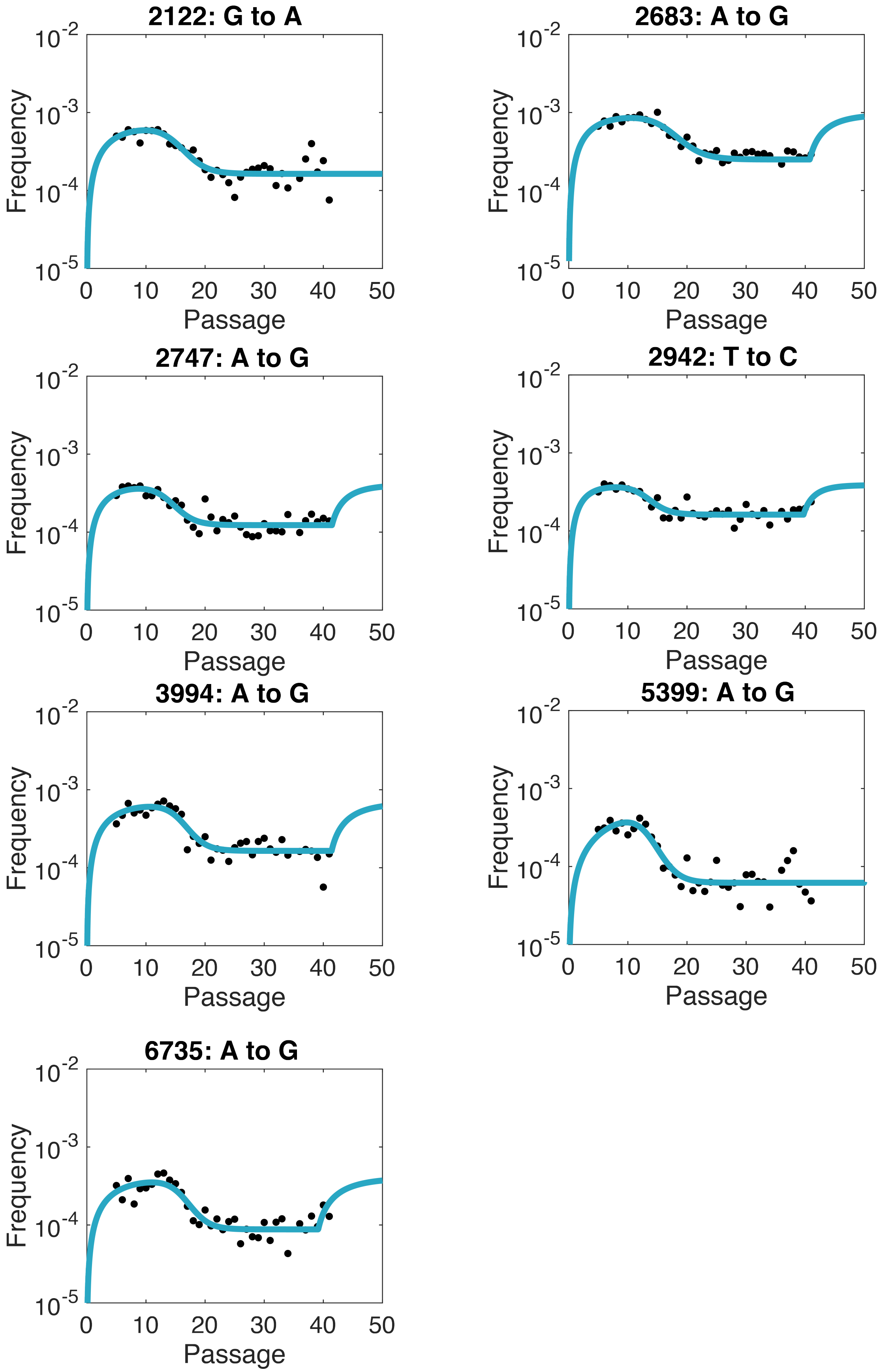
2122: G to A
2683: A to G
2747: A to G
2942: T to C
3994: A to G
5399: A to G
6735: A to G
Frequency
Passage
10^-2
10^-3
10^-4
10^-5
0
10
20
30
40
50

Supplementary Figure 14

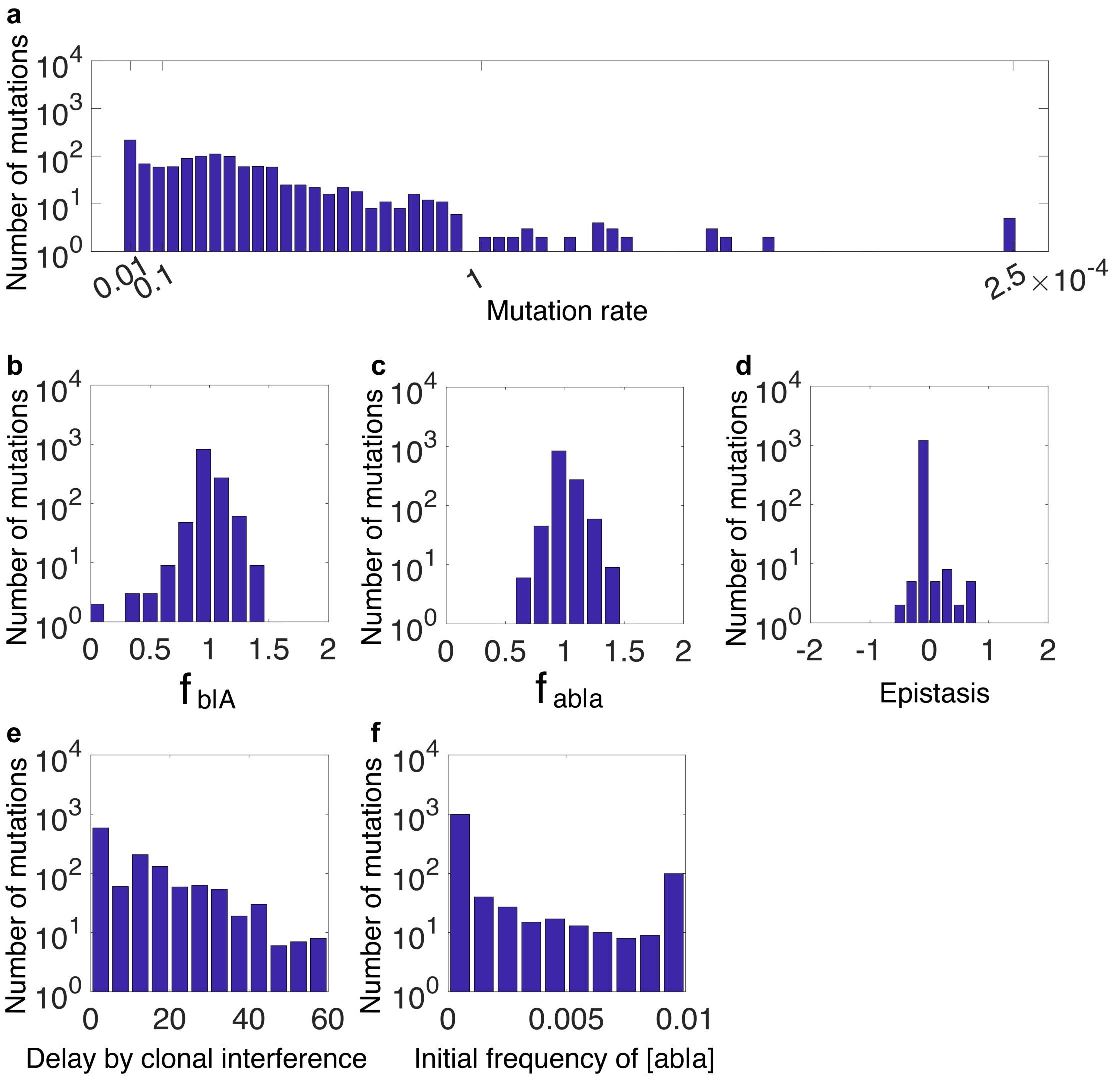


Supplementary Figure 15

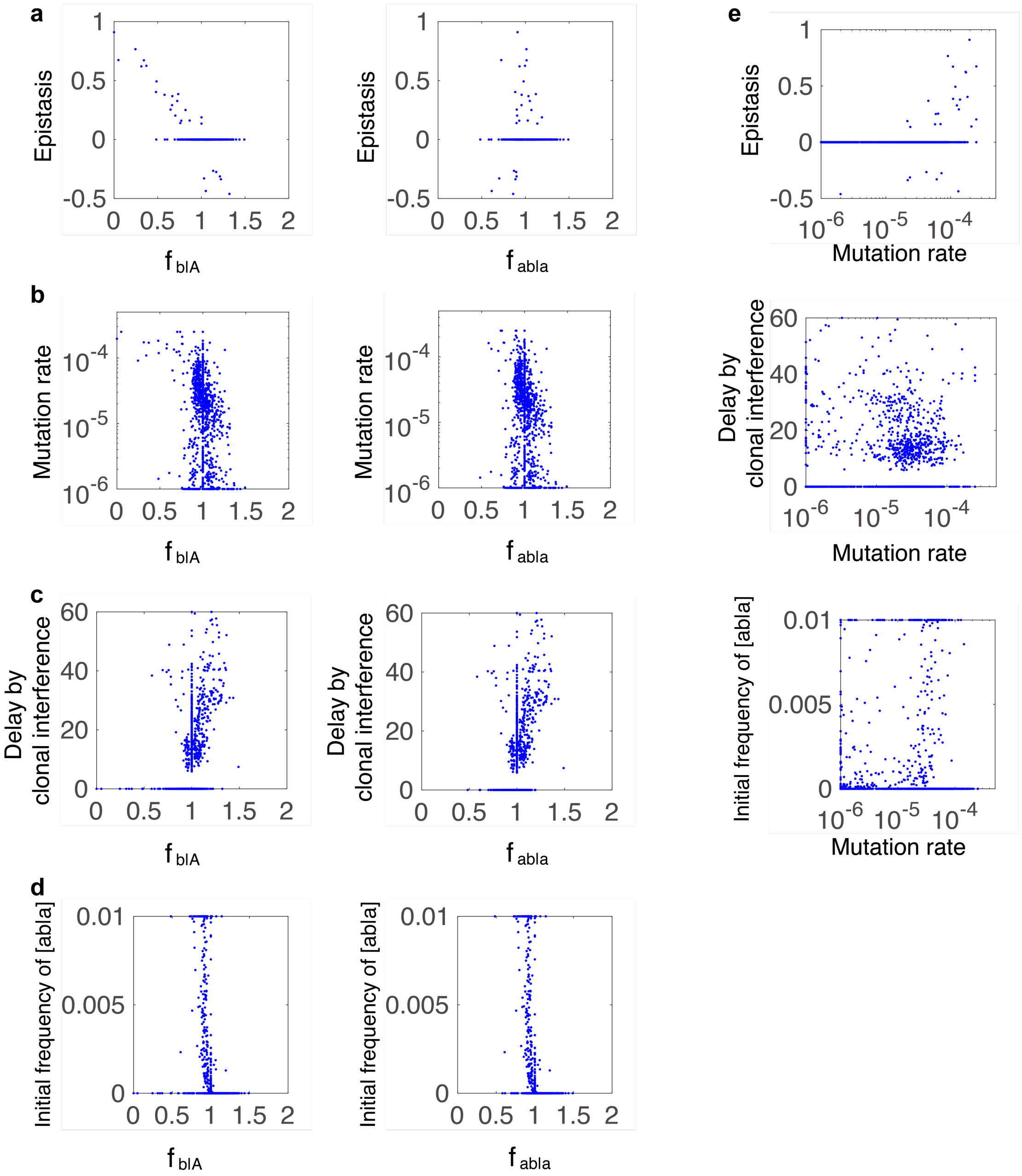


Supplementary Figure 16

**a**

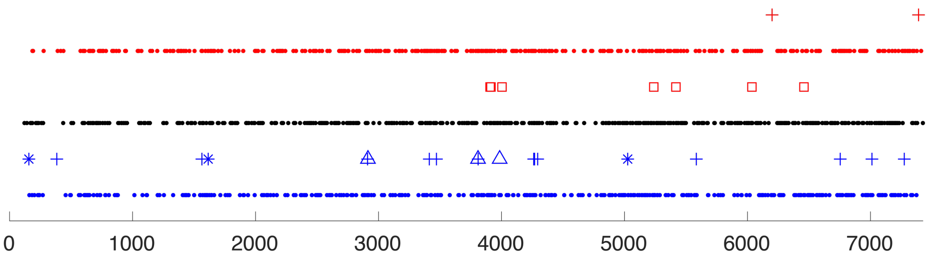


**b**

| FDR matrix | 5' | VP4 | VP2 | VP3 | VP1 | 2A | 2B | 2C | 3A | VPG | 3C | 3D | 3' |
|---|---|---|---|---|---|---|---|---|---|---|---|---|---|
| Beneficial mutation | 1.00E+00 | 1.00E+00 | 1.00E+00 | 1.00E+00 | 7.79E-01 | 1.52E-01 | 2.83E-02 | 1.00E+00 | 2.28E-01 | 3.83E-01 | 1.00E+00 | 4.11E-01 | 1.00E+00 |
| Neutral mutation | 1.00E+00 | 1.00E+00 | 1.00E+00 | 1.00E+00 | 9.57E-01 | 7.79E-01 | 2.44E-01 | 1.00E+00 | 1.59E-01 | 1.00E+00 | 1.00E+00 | 2.00E-04 | 1.00E+00 |
| Detrimental mutation | 1.00E+00 | 1.00E+00 | 1.00E+00 | 1.00E+00 | 1.00E+00 | 7.79E-01 | 9.96E-02 | 4.11E-01 | 9.96E-02 | 1.00E+00 | 1.00E+00 | 2.95E-02 | 1.00E+00 |
| Mutation with flat trajectory | 1.00E+00 | 2.95E-02 | 2.95E-02 | 5.24E-03 | 7.24E-05 | 1.00E+00 | 1.00E+00 | 7.79E-01 | 1.00E+00 | 1.00E+00 | 1.00E+00 | 1.00E+00 | 1.00E+00 |

Supplementary Figure 17

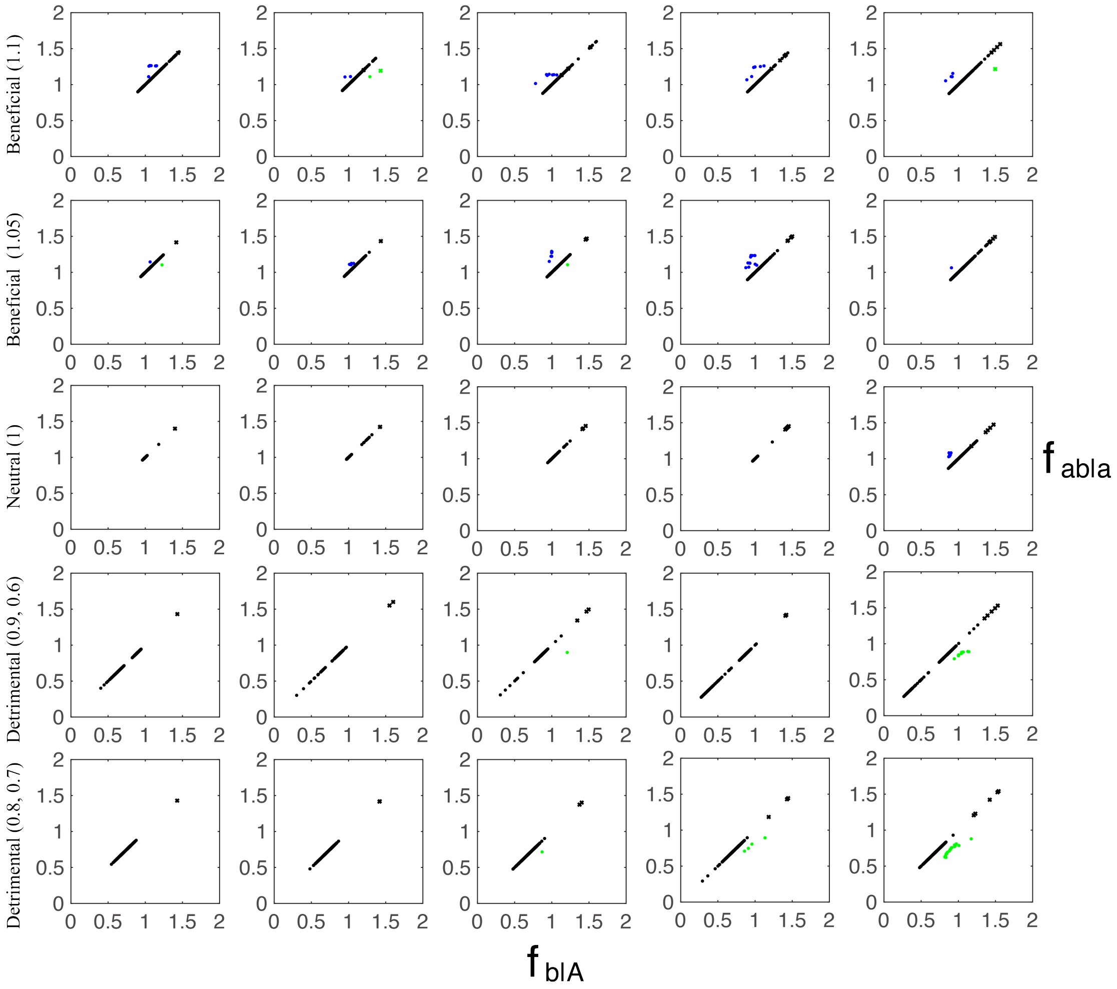


Supplementary Figure 18

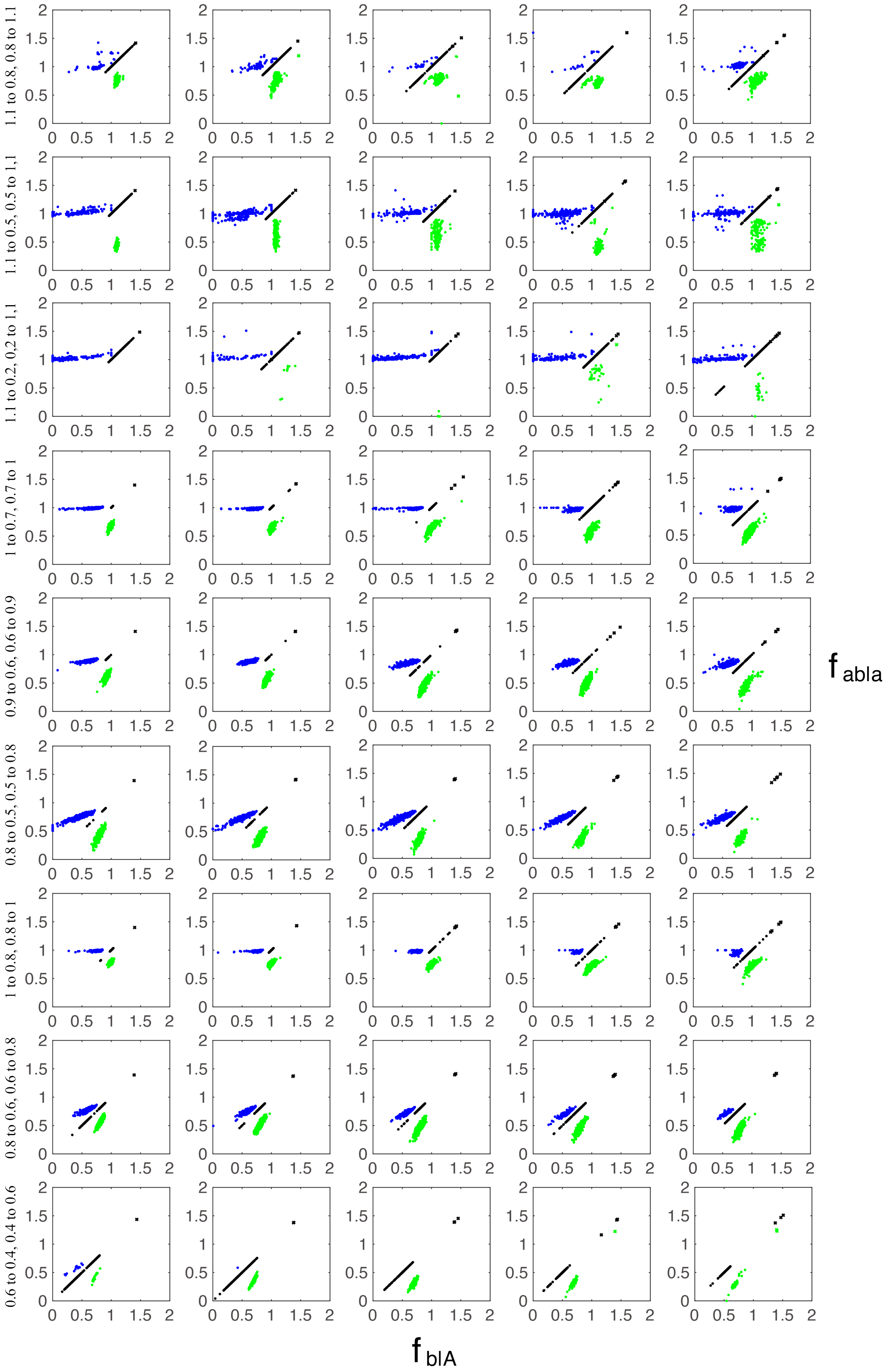
1.1 to 0.8, 0.8 to 1.1
1.1 to 0.5, 0.5 to 1.1
1.1 to 0.2, 0.2 to 1.1
1 to 0.7, 0.7 to 1
0.9 to 0.6, 0.6 to 0.9
0.8 to 0.5, 0.5 to 0.8
1 to 0.8, 0.8 to 1
0.8 to 0.6, 0.6 to 0.8
0.6 to 0.4, 0.4 to 0.6
$f_{abla}$
$f_{blA}$

Supplementary Figure 19

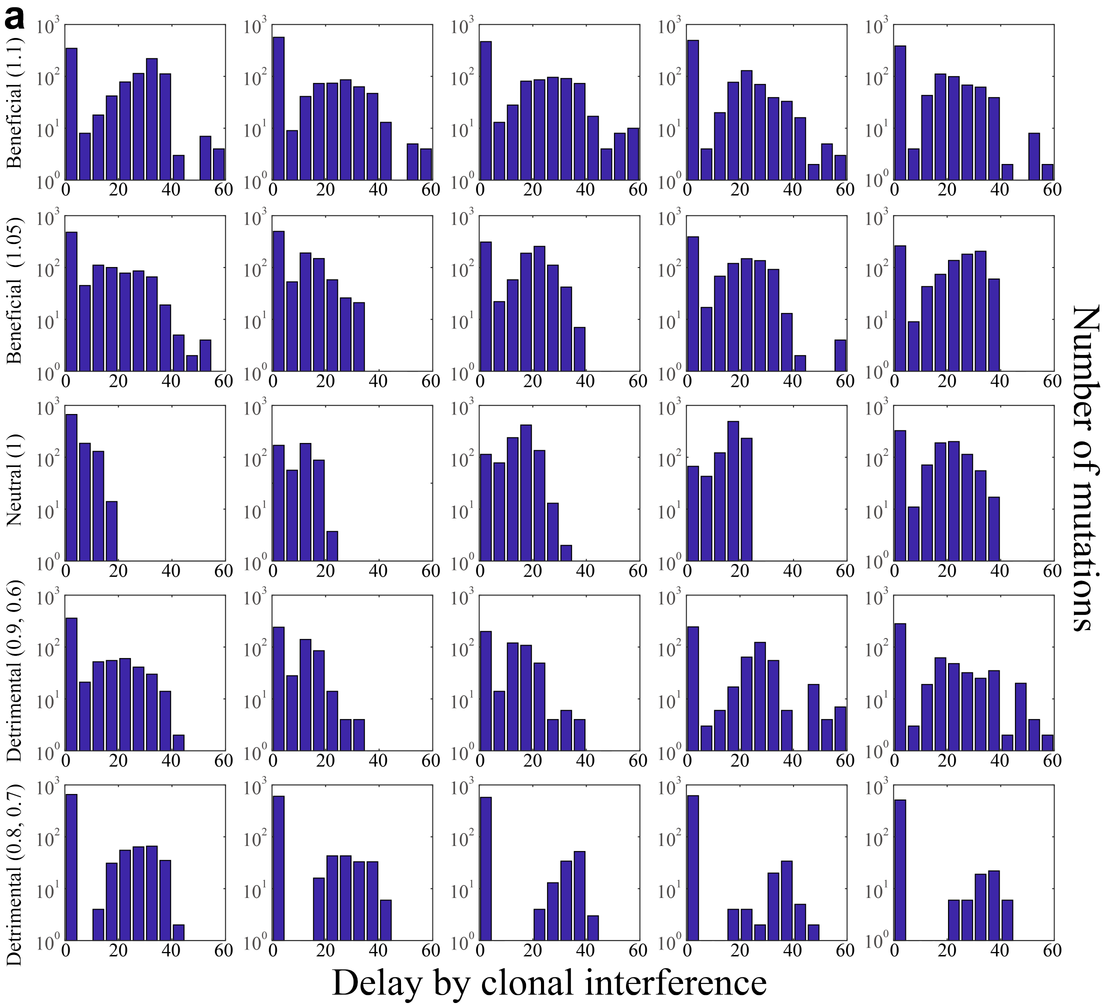


**b**

| Beneficial mutation number | 1 | 2 | 3 | 4 | 5 |
|---|---|---|---|---|---|
| Beneficial mutation (1.1) | 18.9 (15.5) | 11.0 (14.2) | 14.5 (15.6) | 11.5 (14.0) | 13.1 (13.8) |
| Beneficial mutation (1.05) | 11.0 (12.4) | 8.3 (9.3) | 14.7 (11.0) | 14.1 (12.8) | 19.5 (13.2) |
| Neutral mutation (1) | 3.3 (4.9) | 9.1 (6.8) | 14.4 (6.4) | 16.6 (5.6) | 14.8 (11.6) |
| Detrimental mutation (0.9) | 8.5 (10.4) | 7.9 (7.9) | 10.2 (9.0) | 14.8 (13.7) | 11.5 (13.4) |
| Detrimental mutation (0.8) | 7.2 (12.3) | 6.5 (11.8) | 5.8 (12.7) | 3.5 (10.4) | 3.8 (10.7) |
| Detrimental mutation (0.7) | 8.8 (13.7) | 6.3 (13.1) | 4.4 (12.0) | 3.6 (11.4) | 2.9 (10.0) |
| Detrimental mutation (0.6) | 11.9 (15.8) | 7.9 (17.2) | 6.1 (17.4) | 26.5 (25.6) | 24.8 (24.7) |

Supplementary Figure 20

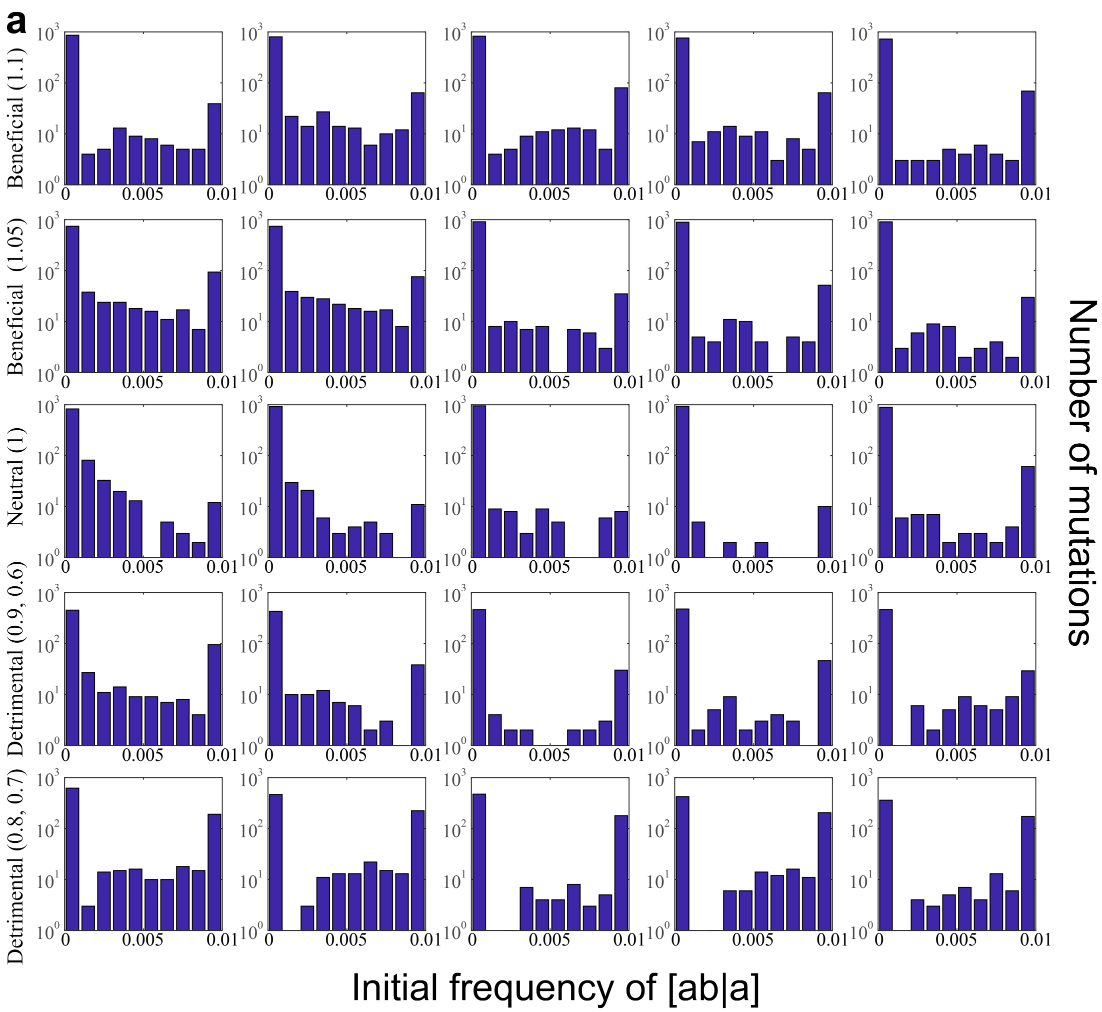


**b**

| Beneficial mutation number | 1 | 2 | 3 | 4 | 5 |
|---|---|---|---|---|---|
| Beneficial mutation (1.1) | 6.9e-04 (2.3e-03) | 1.2e-03 (2.8e-03) | 1.2e-03 (3.0e-03) | 1.1e-03 (2.8e-03) | 1.0e-03 (2.9e-03) |
| Beneficial mutation (1.05) | 1.6e-03 (3.2e-03) | 1.5e-03 (3.0e-03) | 5.7e-04 (2.1e-03) | 7.4e-04 (2.4e-03) | 4.8e-04 (1.9e-03) |
| Neutral mutation (1) | 5.6e-04 (1.5e-03) | 3.4e-04 (1.4e-03) | 2.5e-04 (1.3e-03) | 1.5e-04 (1.1e-03) | 7.6e-04 (2.5e-03) |
| Detrimental mutation (0.9) | 1.3e-03 (2.8e-03) | 9.8e-04 (2.6e-03) | 7.1e-04 (2.5e-03) | 1.1e-03 (3.0e-03) | 1.1e-03 (2.9e-03) |
| Detrimental mutation (0.8) | 1.7e-03 (3.4e-03) | 3.0e-03 (4.2e-03) | 2.6e-03 (4.2e-03) | 3.5e-03 (4.5e-03) | 3.8e-03 (4.6e-03) |
| Detrimental mutation (0.7) | 3.8e-03 (4.7e-03) | 4.5e-03 (4.9e-03) | 3.4e-03 (4.8e-03) | 3.6e-03 (4.8e-03) | 2.7e-03 (4.4e-03) |
| Detrimental mutation (0.6) | 4.9e-03 (4.9e-03) | 4.7e-03 (5.1e-03) | 1.3e-03 (3.5e-03) | 5.1e-04 (2.2e-03) | nan |

Supplementary Figure 21

| Level | Models | Abbreviation | Type | [a] | [b|A] | [ab|a] | Parameter No. |
|---|---|---|---|---|---|---|---|
| 1 | Neutral | N | One-locus | (4) | | | 1 |
| 2 | Beneficial | B | One-locus | (5) | | | 2 |
| 2 | Detrimental | D | One-locus | (6) | | | 2 |
| 3 | Neutral (clonal interference) | N(D) | Two-loci | (5) | (4) | (4) | 4 |
| 3 | Neutral (hitchhiking) | N(I) | Two-loci | (5) | (4) | (4) | 4 |
| 4 | Beneficial (clonal interference) | B(D) | Two-loci | (5) | (5) | (5) | 5 |
| 4 | Beneficial (hitchhiking) | B(I) | Two-loci | (5) | (5) | (5) | 5 |
| 4 | Detrimental (clonal interference) | D(D) | Two-loci | (5) | (6) | (6) | 5 |
| 4 | Detrimental (hitchhiking) | D(I) | Two-loci | (5) | (6) | (6) | 5 |
| 5 | Beneficial→Beneficial | BB | Two-loci | (5) | (5) | (5) | 5 |
| 5 | Beneficial→Detrimental | BD | Two-loci | (5) | (5) | (6) | 5 |
| 5 | Detrimental→Beneficial | DB | Two-loci | (5) | (6) | (5) | 5 |
| 5 | Detrimental→Detrimental | DD | Two-loci | (5) | (6) | (6) | 5 |
| 6 | Mutation with flat trajectory | F | | | | | 1 |

Supplementary Table 1

| Parameter | Lower bound | Upper bound | Initial value | Model type | Annotation |
|---|---|---|---|---|---|
| $\mu_k$ | 1.0E-06 | 2.5E-04 | 1.0E-06 | one-locus | mutation rate |
| $f_k$ | 1.001 | 1.6 | 1.05 | one-locus | fitness of a beneficial mutation |
| | 0 | 0.999 | 0.8 | one-locus | fitness of a detrimental mutation |
| $\mu_a$ | 1.0E-05 | 2.5E-04 | 1.0E-05 | two-loci | mutation rate of mutant *a* |
| $f_a$ | 1.25 | 1.6 | 1.25 | two-loci | fitness of mutant *a* |
| $\mu_b$ | | same as $\mu_k$ | | two-loci | mutation rate of mutant *b* |
| $f_{b\|A}$ | | same as $f_k$ | | two-loci | relative fitness of mutant b with genetic background A |
| $f_{ab\|a}$ | | same as $f_k$ | | two-loci | relative fitness of mutant b with genetic background a |
| *delay* | 0 | 60 | 0 | two-loci | delay by clonal interference |
| $[ab\|a]_0$ | 0 | 0.01 | 0 | two-loci | initial frequency of [ab\|a] by hitchhiking |

Supplementary Table 2

<table>
<tr><th>Mutation subtype</th><th>Description</th></tr>
<tr><td>beneficial; without epistasis</td><td rowspan="3">Mutations that not under epistasis, clonal interference, or hitchhiking.</td></tr>
<tr><td>neutral; without epistasis</td></tr>
<tr><td>detrimental; without epistasis</td></tr>
<tr><td>beneficial; without epistasis (clonal interference)</td><td rowspan="6">Mutations under clonal interference or hitchhiking.</td></tr>
<tr><td>beneficial; without epistasis (hitchhiking)</td></tr>
<tr><td>neutral; without epistasis (clonal interference)</td></tr>
<tr><td>neutral; without epistasis (hitchhiking)</td></tr>
<tr><td>detrimental; without epistasis (clonal interference)</td></tr>
<tr><td>detrimental; without epistasis (hitchhiking)</td></tr>
<tr><td>beneficial; with positive epistasis</td><td rowspan="2">Beneficial mutations with epistasis. In both subtypes, $f_{b|A}$ and $f_{ab|a}$ are larger than 1.</td></tr>
<tr><td>beneficial; with negative epistasis</td></tr>
<tr><td>beneficial to detrimental</td><td rowspan="4">The first word describes $f_{b|A}$, and the last word describes $f_{ab|a}$.</td></tr>
<tr><td>neutral to beneficial</td></tr>
<tr><td>neutral to detrimental</td></tr>
<tr><td>detrimental to beneficial</td></tr>
<tr><td>compensated mutation</td><td>Initially a detrimental mutation, becomes neutral under epistasis.</td></tr>
<tr><td>detrimental; with positive epistasis</td><td rowspan="2">Detrimental mutations with epistasis. In both subtypes, $f_{b|A}$ and $f_{ab|a}$ are smaller than 1.</td></tr>
<tr><td>detrimental; with negative epistasis</td></tr>
<tr><td>establishment</td><td>Mutations with very low frequencies in the initial phase, which raised in the later phase not because of epistasis but the establishment of double mutation.</td></tr>
<tr><td>mutation with flat trajectory</td><td>Mutations with no initial accumulation phase but only flat trajectories.</td></tr>
<tr><td>other</td><td>Mutations that do not belong to any of previous subtypes.</td></tr>
</table>

Supplementary Table 3

| | | Neutral mutation method | | | | | Probabilistic method | | | | |
|---|---|---|---|---|---|---|---|---|---|---|---|
| Mutation rate | Beneficial mutation number | 1 | 2 | 3 | 4 | 5 | 1 | 2 | 3 | 4 | 5 |
| $10^{-4}$ | Start passage (1) | 1.9 | 2.4 | 1.7 | 4.2 | 4.7 | 1 | 1 | 1 | 1 | 1 |
| | Start passage (2) | 3 | 3.5 | 2.7 | 5.6 | 6 | 2 | 2 | 2 | 2 | 2 |
| | Start passage (3) | 4.2 | 4.6 | 3.7 | 6.8 | 7.3 | 3 | 3 | 3 | 3 | 5 |
| | Start passage (4) | 5.4 | 5.7 | 4.5 | 8 | 8.4 | 4 | 4 | 4 | 4 | 7 |
| | Start passage (5) | 6.5 | 6.7 | 5.2 | 9.3 | 9.4 | 6 | 5 | 4 | 6 | 9 |
| | Start passage (6) | 7.5 | 7.7 | 5.9 | 10.4 | 9.9 | 6 | 7 | 5 | 8 | 10 |
| | Start passage (7) | 8.5 | 8.7 | 6.5 | 11.3 | 10.3 | 7 | 8 | 6 | 9 | 11 |
| | Start passage (8) | 9.4 | 9.5 | 6.8 | 12.2 | 10.2 | 7 | 9 | 7 | 11 | 11 |
| $5 \times 10^{-5}$ | Start passage (1) | 1.3 | 2.2 | 1.7 | 1.3 | 5 | 1 | 1 | 1 | 1 | 1 |
| | Start passage (2) | 2.3 | 3.3 | 2.7 | 2.1 | 6.2 | 2 | 2 | 2 | 2 | 2 |
| | Start passage (3) | 3.3 | 4.3 | 3.4 | 2.7 | 7.1 | 3 | 3 | 3 | 3 | 3 |
| | Start passage (4) | 4.3 | 5.3 | 4.1 | 3.4 | 7.7 | 4 | 4 | 4 | 4 | 6 |
| | Start passage (5) | 5.2 | 6 | 4.6 | 3.9 | 8.1 | 5 | 6 | 5 | 6 | 8 |
| | Start passage (6) | 6.1 | 6.8 | 5.1 | 4.3 | 8.2 | 6 | 7 | 6 | 7 | 9 |
| | Start passage (7) | 6.8 | 7.4 | 5.5 | 4.7 | 8.2 | 7 | 8 | 7 | 9 | 10 |
| | Start passage (8) | 7.6 | 7.9 | 5.7 | 4.8 | 8.3 | 8 | 9 | 9 | 10 | 11 |
| $10^{-5}$ | Start passage (1) | | | | | | 1 | 1 | 1 | 1 | 1 |
| | Start passage (2) | | | | | | 2 | 2 | 2 | 2 | 2 |
| | Start passage (3) | | | | | | 3 | 3 | 4 | 3 | 5 |
| | Start passage (4) | | | | | | 4 | 4 | 6 | 5 | 6 |
| | Start passage (5) | | | | | | 4 | 5 | 7 | 6 | 8 |
| | Start passage (6) | | | | | | 5 | 5 | 8 | 7 | 8 |
| | Start passage (7) | | | | | | 5 | 6 | 9 | 8 | 9 |
| | Start passage (8) | | | | | | 6 | 7 | 10 | 9 | 10 |
| $5 \times 10^{-6}$ | Start passage (1) | | | | | | 1 | 1 | 1 | 1 | 1 |
| | Start passage (2) | | | | | | 2 | 2 | 2 | 2 | 2 |
| | Start passage (3) | | | | | | 3 | 2 | 3 | 4 | 3 |
| | Start passage (4) | | | | | | 3 | 3 | 4 | 4 | 4 |
| | Start passage (5) | | | | | | 5 | 5 | 6 | 5 | 5 |
| | Start passage (6) | | | | | | 6 | 6 | 6 | 6 | 6 |
| | Start passage (7) | | | | | | 7 | 6 | 7 | 7 | 7 |
| | Start passage (8) | | | | | | 8 | 7 | 8 | 8 | 8 |
| $10^{-6}$ | Start passage (1) | | | | | | 1 | 1 | 1 | 1 | 1 |
| | Start passage (2) | | | | | | 1 | 1 | 1 | 1 | 1 |
| | Start passage (3) | | | | | | 1 | 1 | 1 | 1 | 2 |
| | Start passage (4) | | | | | | 2 | 2 | 2 | 2 | 2 |
| | Start passage (5) | | | | | | 2 | 2 | 2 | 3 | 2 |
| | Start passage (6) | | | | | | 3 | 3 | 3 | 3 | 3 |
| | Start passage (7) | | | | | | 3 | 4 | 4 | 4 | 4 |
| | Start passage (8) | | | | | | 3 | 5 | 5 | 5 | 5 |

### Supplementary Table 4

| Start passage | 1 | 2 | 3 | 4 | **5** | 6 | 7 | 8 |
|---|---|---|---|---|---|---|---|---|
| neutral; without epistasis | 19 | 35 | 50 | 53 | **75** | 87 | 95 | 93 |
| neutral; without epistasis (delay) | 110 | 180 | 277 | 320 | **317** | 265 | 223 | 184 |
| neutral; without epistasis (hitchhiking) | 43 | 58 | 67 | 75 | **87** | 115 | 144 | 162 |
| beneficial; without epistasis | 28 | 44 | 58 | 74 | **83** | 89 | 105 | 118 |
| beneficial; without epistasis (delay) | 83 | 125 | 160 | 208 | **262** | 298 | 333 | 358 |
| beneficial; without epistasis (hitchhiking) | 3 | 5 | 6 | 9 | **9** | 13 | 15 | 18 |
| detrimental; without epistasis | 76 | 111 | 93 | 82 | **67** | 55 | 30 | 23 |
| detrimental; without epistasis (delay) | 94 | 135 | 120 | 93 | **65** | 53 | 39 | 34 |
| detrimental; without epistasis (hitchhiking) | 200 | 217 | 230 | 233 | **233** | 238 | 243 | 229 |
| beneficial; with negative epistasis | 1 | 0 | 0 | 0 | **0** | 0 | 0 | 0 |
| beneficial to detrimental | 2 | 8 | 9 | 9 | **7** | 2 | 2 | 1 |
| neutral to beneficial | 1 | 1 | 2 | 2 | **2** | 1 | 1 | 2 |
| neutral to detrimental | 2 | 0 | 0 | 1 | **0** | 0 | 0 | 0 |
| detrimental to beneficial | 5 | 7 | 4 | 2 | **3** | 0 | 0 | 0 |
| compensated mutation | 75 | 61 | 42 | 24 | **13** | 9 | 6 | 8 |
| detrimental; with positive epistasis | 52 | 25 | 15 | 7 | **3** | 1 | 1 | 1 |
| detrimental; with negative epistasis | 0 | 1 | 1 | 0 | **0** | 1 | 2 | 2 |
| establishment | 4 | 5 | 5 | 5 | **4** | 5 | 5 | 6 |
| mutation with flat trajectory | 13353 | 13204 | 13120 | 13077 | **13052** | 13051 | 13041 | 13036 |
| other | 6880 | 6809 | 6772 | 6757 | **6749** | 6748 | 6746 | 6756 |

Supplementary Table 5

**a**

| mutation type | no epistais | epistasis | mutation with flat trajectory | other |
|---|---|---|---|---|
| C -> U | 71 | 4 | 1567 | 79 |
| G -> A | 90 | 3 | 579 | 1015 |
| U -> C | 361 | 9 | 1054 | 348 |
| A -> G | 403 | 11 | 1484 | 294 |
| G -> U | 14 | 0 | 468 | 1199 |
| C -> A | 37 | 0 | 899 | 749 |
| G -> C | 8 | 0 | 860 | 787 |
| U -> A | 73 | 1 | 1229 | 426 |
| A -> U | 76 | 0 | 1472 | 615 |
| C -> G | 11 | 0 | 974 | 130 |
| A -> C | 37 | 0 | 1357 | 679 |
| U -> G | 17 | 0 | 1109 | 428 |
| total | 1198 | 28 | 13052 | 6749 |

**b**

| mutation type | positive epistasis | negative epistasis |
|---|---|---|
| C -> U | 4 | 0 |
| G -> A | 3 | 0 |
| U -> C | 4 | 5 |
| A -> G | 10 | 1 |
| G -> U | 0 | 0 |
| C -> A | 0 | 0 |
| G -> C | 0 | 0 |
| U -> A | 0 | 1 |
| A -> U | 0 | 0 |
| C -> G | 0 | 0 |
| A -> C | 0 | 0 |
| U -> G | 0 | 0 |
| total | 21 | 7 |

**c**

| | beneficial | | | neutral | | | detrimental | | |
|---|---|---|---|---|---|---|---|---|---|
| mutation type | o | ci | h | o | ci | h | o | ci | h |
| C -> U | 2 | 21 | 1 | 7 | 15 | 4 | 3 | 4 | 14 |
| G -> A | 4 | 33 | 0 | 2 | 30 | 2 | 0 | 5 | 14 |
| U -> C | 16 | 67 | 3 | 32 | 126 | 12 | 33 | 24 | 48 |
| A -> G | 21 | 58 | 2 | 32 | 135 | 19 | 31 | 26 | 79 |
| G -> U | 0 | 5 | 0 | 0 | 0 | 4 | 0 | 1 | 4 |
| C -> A | 4 | 12 | 0 | 0 | 3 | 5 | 0 | 0 | 13 |
| G -> C | 3 | 1 | 0 | 0 | 0 | 1 | 0 | 0 | 3 |
| U -> A | 14 | 22 | 0 | 0 | 3 | 18 | 0 | 0 | 16 |
| A -> U | 12 | 22 | 2 | 1 | 2 | 13 | 0 | 0 | 24 |
| C -> G | 0 | 3 | 0 | 0 | 2 | 0 | 0 | 2 | 4 |
| A -> C | 5 | 13 | 1 | 1 | 0 | 7 | 0 | 1 | 9 |
| U -> G | 2 | 5 | 0 | 0 | 1 | 2 | 0 | 2 | 5 |
| total | 83 | 262 | 9 | 75 | 317 | 87 | 67 | 65 | 233 |

Supplementary Table 6

| Locus | Coding region | Mutation | Codon mutation | Protein mutation | Mutation type | Estimation |
|---|---|---|---|---|---|---|
| 2959 | VP1 | A to G | GUA to GUG | V160V | synonymous | 0.00 |
| 2650 | VP1 | U to C | CCU to CCC | P57P | synonymous | 0.00 (d=12.2) |
| 673 | 5' | U to C | UCC to CCC | S225P | nonsynonymous | 0.00 (i=0.00036) |
| 6254 | 3D | A to G | AUC to GUC | I90V | nonsynonymous | 0.05 |
| 3173 | VP1 | G to U | GCA to UCA | A232S | nonsynonymous | 0.49 (d=7.4) |
| 2065 | VP3 | U to C | CUU to CUC | L100L | synonymous | 0.04 (d=16.4) |
| 5712 | 3C | C to A | ACU to AAU | T92N | nonsynonymous | 0.21 (d>50) |
| 3942 | 2B | C to U | ACU to AUU | T37I | nonsynonymous | 0.14 (i=0.01) |
| 2509 | VP1 | U to C | AUU to AUC | I10I | synonymous | -0.03 |
| 1552 | VP2 | G to A | CGG to CGA | R198R | synonymous | -0.05 (d=14.4) |
| 3814 | 2A | A to G | GAA to GAG | E143E | synonymous | -0.08 (i=0.0082) |
| 7213 | 3D | C to U | AAC to AAU | N409N | synonymous | mutation with flat trajectory |
| 6036 | 3D | U to C | AUA to ACA | I17T | nonsynonymous | 0.21 to -0.10 |
| 3985 | 2B | A to G | CUA to CUG | L51L | synonymous | -0.52 to -0.12 |
| 4269 | 2C | A to G | AAA to AGA | K49R | nonsynonymous | -0.33 to 0.03 |
| 5026 | 2C | C to U | UAC to UAU | Y301Y | synonymous | -0.27 to 0.12 |

Supplementary Table 7

**a**

| Beneficial mutation number | 1 | 2 | 3 | 4 | 5 |
|---|---|---|---|---|---|
| Beneficial mutation (1.1) | 0.008 | 0.003 | 0.007 | 0.008 | 0.004 |
| Beneficial mutation (1.05) | 0.002 | 0.005 | 0.008 | 0.013 | 0.001 |
| Neutral mutation (1) | 0.000 | 0.000 | 0.000 | 0.000 | 0.006 |
| Detrimental mutation (0.9) | 0.000 | 0.000 | 0.002 | 0.000 | 0.016 |
| Detrimental mutation (0.8) | 0.000 | 0.000 | 0.002 | 0.006 | 0.028 |
| Detrimental mutation (0.7) | 0.000 | 0.000 | 0.000 | 0.002 | 0.016 |
| Detrimental mutation (0.6) | 0.000 | 0.000 | 0.000 | 0.000 | 0.000 |

**b**

| Beneficial mutation number | 1 | 2 | 3 | 4 | 5 |
|---|---|---|---|---|---|
| Negative epistasis (1.1 to 0.8) | 0.914 | 0.452 | 0.660 | 0.494 | 0.687 |
| Positive epistasis (0.8 to 1.1) | 0.130 | 0.156 | 0.064 | 0.034 | 0.244 |
| Negative epistasis (1.1 to 0.5) | 0.361 | 0.347 | 0.298 | 0.393 | 0.271 |
| Positive epistasis (0.5 to 1.1) | 0.320 | 0.398 | 0.300 | 0.430 | 0.326 |
| Negative epistasis (1.1 to 0.2) | 0.000 | 0.004 | 0.008 | 0.062 | 0.048 |
| Positive epistasis (0.2 to 1.1) | 0.366 | 0.174 | 0.458 | 0.332 | 0.360 |
| Negative epistasis (1 to 0.7) | 1.000 | 1.000 | 0.996 | 0.879 | 0.774 |
| Positive epistasis (0.7 to 1) | 0.926 | 0.516 | 0.430 | 0.464 | 0.316 |
| Negative epistasis (0.9 to 0.6) | 1.000 | 1.000 | 0.909 | 0.950 | 0.772 |
| Positive epistasis (0.6 to 0.9) | 0.890 | 0.802 | 0.798 | 0.828 | 0.730 |
| Negative epistasis (0.8 to 0.5) | 0.988 | 0.936 | 0.581 | 0.675 | 0.432 |
| Positive epistasis (0.5 to 0.8) | 0.982 | 0.882 | 0.782 | 0.696 | 0.528 |
| Negative epistasis (1 to 0.8) | 0.996 | 1.000 | 0.994 | 0.929 | 0.816 |
| Positive epistasis (0.8 to 1) | 0.464 | 0.344 | 0.270 | 0.208 | 0.292 |
| Negative epistasis (0.8 to 0.6) | 0.848 | 0.988 | 0.976 | 0.885 | 0.822 |
| Positive epistasis (0.6 to 0.8) | 0.856 | 0.720 | 0.440 | 0.272 | 0.178 |
| Negative epistasis (0.6 to 0.4) | 0.042 | 0.197 | 0.308 | 0.147 | 0.091 |
| Positive epistasis (0.4 to 0.6) | 0.026 | 0.002 | 0.000 | 0.000 | 0.000 |

Supplementary Table 8

**a**

| Beneficial mutation number | 1 | 2 | 3 | 4 | 5 |
|---|---|---|---|---|---|
| Beneficial mutation (1.1) | 0.003 | 0.003 | 0.010 | 0.006 | 0.011 |
| Beneficial mutation (1.05) | 0.000 | 0.001 | 0.003 | 0.005 | 0.004 |
| Neutral mutation (1) | 0.001 | 0.003 | 0.004 | 0.004 | 0.008 |
| Detrimental mutation (0.9) | 0.002 | 0.000 | 0.000 | 0.006 | 0.004 |
| Detrimental mutation (0.8) | 0.000 | 0.004 | 0.002 | 0.002 | 0.002 |
| Detrimental mutation (0.7) | 0.000 | 0.000 | 0.000 | 0.000 | 0.002 |
| Detrimental mutation (0.6) | 0.000 | 0.000 | 0.000 | 0.000 | 0.000 |

**b**

| Beneficial mutation number | 1 | 2 | 3 | 4 | 5 |
|---|---|---|---|---|---|
| Negative epistasis (1.1 to 0.8) | 0.465 | 0.622 | 0.290 | 0.488 | 0.283 |
| Positive epistasis (0.8 to 1.1) | 0.088 | 0.128 | 0.156 | 0.164 | 0.106 |
| Negative epistasis (1.1 to 0.5) | 0.383 | 0.444 | 0.133 | 0.123 | 0.093 |
| Positive epistasis (0.5 to 1.1) | 0.596 | 0.458 | 0.326 | 0.238 | 0.220 |
| Negative epistasis (1.1 to 0.2) | 0.056 | 0.114 | 0.010 | 0.002 | 0.016 |
| Positive epistasis (0.2 to 1.1) | 0.396 | 0.534 | 0.288 | 0.380 | 0.332 |
| Negative epistasis (1 to 0.7) | 0.433 | 0.376 | 0.336 | 0.393 | 0.202 |
| Positive epistasis (0.7 to 1) | 0.236 | 0.194 | 0.178 | 0.182 | 0.214 |
| Negative epistasis (0.9 to 0.6) | 0.076 | 0.064 | 0.064 | 0.065 | 0.048 |
| Positive epistasis (0.6 to 0.9) | 0.200 | 0.142 | 0.112 | 0.066 | 0.048 |
| Negative epistasis (0.8 to 0.5) | 0.006 | 0.008 | 0.006 | 0.010 | 0.010 |
| Positive epistasis (0.5 to 0.8) | 0.006 | 0.002 | 0.000 | 0.000 | 0.000 |
| Negative epistasis (1 to 0.8) | 0.267 | 0.235 | 0.181 | 0.131 | 0.176 |
| Positive epistasis (0.8 to 1) | 0.094 | 0.120 | 0.068 | 0.068 | 0.060 |
| Negative epistasis (0.8 to 0.6) | 0.006 | 0.002 | 0.004 | 0.008 | 0.004 |
| Positive epistasis (0.6 to 0.8) | 0.010 | 0.002 | 0.000 | 0.002 | 0.002 |
| Negative epistasis (0.6 to 0.4) | 0.000 | 0.000 | 0.000 | 0.000 | 0.000 |
| Positive epistasis (0.4 to 0.6) | 0.000 | 0.000 | 0.000 | 0.000 | 0.000 |

Supplementary Table 9

**a**

| Beneficial mutation number | 1 | 2 | 3 | 4 | 5 |
|---|---|---|---|---|---|
| Beneficial mutation (1.1) | 0.002 | 0.002 | 0.003 | 0.001 | 0.002 |
| Beneficial mutation (1.05) | 0.001 | 0.001 | 0.001 | 0.001 | 0.001 |
| Neutral mutation (1) | 0.000 | 0.001 | 0.001 | 0.000 | 0.000 |
| Detrimental mutation (0.9) | 0.000 | 0.000 | 0.000 | 0.000 | 0.000 |
| Detrimental mutation (0.8) | 0.000 | 0.000 | 0.000 | 0.000 | 0.000 |
| Detrimental mutation (0.7) | 0.000 | 0.000 | 0.000 | 0.000 | 0.000 |
| Detrimental mutation (0.6) | 0.000 | 0.000 | 0.000 | 0.000 | 0.000 |

**b**

| Beneficial mutation number | 1 | 2 | 3 | 4 | 5 |
|---|---|---|---|---|---|
| Negative epistasis (1.1 to 0.8) | 0.010 | 0.002 | 0.000 | 0.004 | 0.002 |
| Positive epistasis (0.8 to 1.1) | 0.000 | 0.000 | 0.004 | 0.000 | 0.000 |
| Negative epistasis (1.1 to 0.5) | 0.006 | 0.006 | 0.002 | 0.000 | 0.002 |
| Positive epistasis (0.5 to 1.1) | 0.002 | 0.000 | 0.002 | 0.000 | 0.000 |
| Negative epistasis (1.1 to 0.2) | 0.008 | 0.000 | 0.000 | 0.000 | 0.000 |
| Positive epistasis (0.2 to 1.1) | 0.000 | 0.000 | 0.000 | 0.000 | 0.000 |
| Negative epistasis (1 to 0.7) | 0.000 | 0.002 | 0.006 | 0.000 | 0.000 |
| Positive epistasis (0.7 to 1) | 0.000 | 0.000 | 0.000 | 0.000 | 0.000 |
| Negative epistasis (0.9 to 0.6) | 0.000 | 0.000 | 0.000 | 0.000 | 0.000 |
| Positive epistasis (0.6 to 0.9) | 0.000 | 0.000 | 0.000 | 0.000 | 0.000 |
| Negative epistasis (0.8 to 0.5) | 0.000 | 0.000 | 0.000 | 0.000 | 0.000 |
| Positive epistasis (0.5 to 0.8) | 0.000 | 0.000 | 0.000 | 0.000 | 0.000 |
| Negative epistasis (1 to 0.8) | 0.004 | 0.000 | 0.000 | 0.000 | 0.000 |
| Positive epistasis (0.8 to 1) | 0.000 | 0.000 | 0.000 | 0.000 | 0.000 |
| Negative epistasis (0.8 to 0.6) | 0.000 | 0.000 | 0.000 | 0.000 | 0.000 |
| Positive epistasis (0.6 to 0.8) | 0.000 | 0.000 | 0.000 | 0.000 | 0.000 |
| Negative epistasis (0.6 to 0.4) | 0.000 | 0.000 | 0.000 | 0.000 | 0.000 |
| Positive epistasis (0.4 to 0.6) | 0.000 | 0.000 | 0.000 | 0.000 | 0.000 |

Supplementary Table 10

**a**

| Beneficial mutation number | 1 | 2 | 3 | 4 | 5 |
|---|---|---|---|---|---|
| Beneficial mutation (1.1) | 0/19 | 0/18 | 0/17 | 0/16 | 0/15 |
| Beneficial mutation (1.05) | 0/19 | 0/18 | 0/17 | 1/16 | 0/15 |

**b**

| Beneficial mutation number | 1 | 2 | 3 | 4 | 5 |
|---|---|---|---|---|---|
| Negative epistasis (1.2 to 1.05) | 0/9 | 0/8 | 0/7 | 0/6 | 0/5 |
| Positive epistasis (1.05 to 1.2) | 0/10 | 0/10 | 0/10 | 0/10 | 0/10 |
| Negative epistasis (1.2 to 1) | 0/9 | 0/8 | 0/7 | 0/6 | 0/5 |
| Positive epistasis (1 to 1.2) | 0/10 | 1/10 | 2/10 | 1/10 | 0/10 |
| Negative epistasis (1.3 to 1.15) | 0/9 | 0/8 | 0/7 | 0/6 | 0/5 |
| Positive epistasis (1.15 to 1.3) | 1/10 | 0/10 | 0/10 | 0/10 | 0/10 |
| Negative epistasis (1.4 to 1.15) | 0/9 | 0/8 | 2/7 | 0/6 | 0/5 |
| Positive epistasis (1.15 to 1.4) | 0/10 | 1/10 | 0/10 | 1/10 | 0/10 |
| Negative epistasis (1.45 to 1.15) | 1/9 | 0/8 | 2/7 | 1/6 | 0/5 |
| Positive epistasis (1.15 to 1.45) | 0/10 | 1/10 | 0/10 | 1/10 | 1/10 |

Supplementary Table 11

| Beneficial mutation number | 1 | 2 | 3 | 4 | 5 |
|---|---|---|---|---|---|
| Beneficial mutation (1.1) | 8.20E-05 | 9.30E-05 | 1.00E-04 | 9.40E-05 | 1.10E-04 |
| Beneficial mutation (1.05) | 1.10E-04 | 9.20E-05 | 9.50E-05 | 1.00E-04 | 9.70E-05 |
| Neutral mutation (1) | 9.00E-05 | 8.60E-05 | 9.00E-05 | 9.40E-05 | 9.50E-05 |
| Detrimental mutation (0.9) | 9.70E-05 | 9.90E-05 | 1.00E-04 | 9.80E-05 | 9.90E-05 |
| Detrimental mutation (0.8) | 9.70E-05 | 9.90E-05 | 9.90E-05 | 9.00E-05 | 9.00E-05 |
| Detrimental mutation (0.7) | 9.70E-05 | 9.30E-05 | 9.00E-05 | 8.30E-05 | 8.30E-05 |
| Detrimental mutation (0.6) | 8.80E-05 | 8.70E-05 | 1.00E-04 | 1.30E-04 | 1.20E-04 |
| Negative epistasis (1.1 to 0.8) | 8.30E-05 | 1.10E-04 | 8.40E-05 | 9.60E-05 | 8.80E-05 |
| Positive epistasis (0.8 to 1.1) | 4.90E-05 | 5.80E-05 | 5.70E-05 | 6.30E-05 | 6.60E-05 |
| Negative epistasis (1.1 to 0.5) | 1.00E-04 | 9.80E-05 | 9.40E-05 | 9.80E-05 | 1.00E-04 |
| Positive epistasis (0.5 to 1.1) | 4.90E-05 | 5.70E-05 | 4.70E-05 | 6.80E-05 | 5.50E-05 |
| Negative epistasis (1.1 to 0.2) | 8.50E-05 | 9.80E-05 | 8.20E-05 | 9.60E-05 | 9.90E-05 |
| Positive epistasis (0.2 to 1.1) | 4.30E-05 | 3.70E-05 | 4.90E-05 | 4.70E-05 | 4.90E-05 |
| Negative epistasis (1 to 0.7) | 9.40E-05 | 9.30E-05 | 8.50E-05 | 9.50E-05 | 1.10E-04 |
| Positive epistasis (0.7 to 1) | 9.10E-05 | 7.20E-05 | 6.70E-05 | 7.10E-05 | 6.60E-05 |
| Negative epistasis (0.9 to 0.6) | 8.50E-05 | 8.50E-05 | 1.00E-04 | 8.90E-05 | 1.00E-04 |
| Positive epistasis (0.6 to 0.9) | 8.80E-05 | 8.30E-05 | 8.90E-05 | 8.60E-05 | 9.10E-05 |
| Negative epistasis (0.8 to 0.5) | 1.00E-04 | 9.90E-05 | 1.20E-04 | 1.10E-04 | 1.10E-04 |
| Positive epistasis (0.5 to 0.8) | 1.10E-04 | 9.80E-05 | 1.00E-04 | 9.40E-05 | 9.20E-05 |
| Negative epistasis (1 to 0.8) | 9.30E-05 | 8.90E-05 | 9.00E-05 | 9.80E-05 | 9.60E-05 |
| Positive epistasis (0.8 to 1) | 8.10E-05 | 7.70E-05 | 7.50E-05 | 7.30E-05 | 7.80E-05 |
| Negative epistasis (0.8 to 0.6) | 9.60E-05 | 9.40E-05 | 9.80E-05 | 1.00E-04 | 1.10E-04 |
| Positive epistasis (0.6 to 0.8) | 9.50E-05 | 9.30E-05 | 8.90E-05 | 8.50E-05 | 8.50E-05 |
| Negative epistasis (0.6 to 0.4) | 1.20E-04 | 1.10E-04 | 1.10E-04 | 1.10E-04 | 1.10E-04 |
| Positive epistasis (0.4 to 0.6) | 7.00E-05 | 7.30E-05 | 7.50E-05 | 8.60E-05 | nan |

Supplementary Table 12

| | $f_{blA}$ | | | | | $f_{abla}$ | | | | |
|---|---|---|---|---|---|---|---|---|---|---|
| Beneficial mutation number | 1 | 2 | 3 | 4 | 5 | 1 | 2 | 3 | 4 | 5 |
| Beneficial mutation (1.1) | 1.06 | 1.03 | 1.03 | 1.02 | 1.01 | 1.06 | 1.03 | 1.03 | 1.02 | 1.01 |
| Beneficial mutation (1.05) | 1.03 | 1.03 | 1.03 | 1.00 | 1.02 | 1.03 | 1.03 | 1.03 | 1.00 | 1.02 |
| Neutral mutation (1) | 1.00 | 1.00 | 1.00 | 1.00 | 0.97 | 1.00 | 1.00 | 1.00 | 1.00 | 0.97 |
| Detrimental mutation (0.9) | 0.89 | 0.88 | 0.86 | 0.86 | 0.85 | 0.89 | 0.88 | 0.86 | 0.86 | 0.85 |
| Detrimental mutation (0.8) | 0.78 | 0.75 | 0.74 | 0.75 | 0.74 | 0.78 | 0.75 | 0.74 | 0.75 | 0.73 |
| Detrimental mutation (0.7) | 0.68 | 0.66 | 0.65 | 0.66 | 0.65 | 0.68 | 0.66 | 0.65 | 0.66 | 0.64 |
| Detrimental mutation (0.6) | 0.62 | 0.58 | 0.48 | 0.41 | 0.40 | 0.62 | 0.58 | 0.48 | 0.41 | 0.40 |
| Negative epistasis (1.1 to 0.8) | 1.10 | 1.03 | 1.09 | 1.07 | 1.08 | 0.78 | 0.86 | 0.85 | 0.88 | 0.83 |
| Positive epistasis (0.8 to 1.1) | 1.02 | 0.99 | 0.95 | 0.94 | 0.92 | 1.05 | 1.03 | 0.95 | 0.95 | 0.99 |
| Negative epistasis (1.1 to 0.5) | 1.07 | 1.06 | 1.07 | 1.06 | 1.05 | 0.83 | 0.90 | 0.93 | 0.79 | 0.92 |
| Positive epistasis (0.5 to 1.1) | 0.86 | 0.80 | 0.87 | 0.74 | 0.82 | 1.05 | 1.02 | 1.03 | 1.02 | 1.01 |
| Negative epistasis (1.1 to 0.2) | 1.09 | 1.07 | 1.11 | 1.07 | 1.06 | 1.09 | 1.07 | 1.10 | 1.04 | 1.02 |
| Positive epistasis (0.2 to 1.1) | 0.77 | 0.84 | 0.69 | 0.72 | 0.73 | 1.05 | 1.04 | 1.03 | 1.03 | 1.02 |
| Negative epistasis (1 to 0.7) | 0.99 | 1.01 | 1.01 | 0.98 | 0.93 | 0.68 | 0.65 | 0.65 | 0.63 | 0.60 |
| Positive epistasis (0.7 to 1) | 0.73 | 0.86 | 0.88 | 0.86 | 0.89 | 1.00 | 1.00 | 1.00 | 0.99 | 0.98 |
| Negative epistasis (0.9 to 0.6) | 0.92 | 0.93 | 0.87 | 0.91 | 0.87 | 0.62 | 0.58 | 0.50 | 0.52 | 0.50 |
| Positive epistasis (0.6 to 0.9) | 0.66 | 0.70 | 0.68 | 0.67 | 0.66 | 0.90 | 0.89 | 0.87 | 0.87 | 0.86 |
| Negative epistasis (0.8 to 0.5) | 0.80 | 0.80 | 0.73 | 0.78 | 0.73 | 0.44 | 0.42 | 0.47 | 0.46 | 0.53 |
| Positive epistasis (0.5 to 0.8) | 0.48 | 0.53 | 0.51 | 0.56 | 0.56 | 0.76 | 0.75 | 0.72 | 0.73 | 0.73 |
| Negative epistasis (1 to 0.8) | 0.99 | 1.00 | 1.00 | 0.99 | 0.99 | 0.79 | 0.77 | 0.75 | 0.72 | 0.74 |
| Positive epistasis (0.8 to 1) | 0.90 | 0.92 | 0.94 | 0.94 | 0.91 | 1.00 | 0.99 | 0.99 | 0.99 | 0.97 |
| Negative epistasis (0.8 to 0.6) | 0.78 | 0.81 | 0.80 | 0.77 | 0.77 | 0.57 | 0.53 | 0.48 | 0.46 | 0.45 |
| Positive epistasis (0.6 to 0.8) | 0.63 | 0.65 | 0.68 | 0.70 | 0.70 | 0.78 | 0.77 | 0.76 | 0.75 | 0.74 |
| Negative epistasis (0.6 to 0.4) | 0.40 | 0.45 | 0.55 | 0.52 | 0.53 | 0.38 | 0.38 | 0.43 | 0.46 | 0.49 |
| Positive epistasis (0.4 to 0.6) | 0.66 | 0.63 | 0.60 | 0.50 | nan | 0.68 | 0.63 | 0.60 | 0.50 | nan |

Supplementary Table 13

| | | Mutation with flat trajectory | | | | | Other | | | | |
|---|---|---|---|---|---|---|---|---|---|---|---|
| | beneficial mutation number | 1 | 2 | 3 | 4 | 5 | 1 | 2 | 3 | 4 | 5 |
| | Beneficial mutation (1.1) | 0.009 | 0.000 | 0.004 | 0.002 | 0.072 | 0.036 | 0.017 | 0.021 | 0.106 | 0.104 |
| | Beneficial mutation (1.05) | 0.000 | 0.000 | 0.000 | 0.003 | 0.003 | 0.005 | 0.002 | 0.004 | 0.006 | 0.023 |
| | Neutral mutation (1) | 0.000 | 0.000 | 0.000 | 0.000 | 0.002 | 0.000 | 0.000 | 0.000 | 0.045 | 0.010 |
| $10^{-4}$ | Detrimental mutation (0.9) | 0.000 | 0.002 | 0.004 | 0.008 | 0.034 | 0.000 | 0.000 | 0.000 | 0.010 | 0.004 |
| | Detrimental mutation (0.8) | 0.006 | 0.012 | 0.064 | 0.060 | 0.180 | 0.000 | 0.000 | 0.004 | 0.008 | 0.008 |
| | Detrimental mutation (0.7) | 0.170 | 0.428 | 0.564 | 0.544 | 0.664 | 0.000 | 0.000 | 0.000 | 0.000 | 0.000 |
| | Detrimental mutation (0.6) | 0.730 | 0.962 | 0.984 | 0.882 | 0.894 | 0.000 | 0.000 | 0.000 | 0.000 | 0.000 |
| | Beneficial mutation (1.1) | 0.000 | 0.001 | 0.001 | 0.015 | 0.050 | 0.003 | 0.008 | 0.013 | 0.029 | 0.075 |
| | Beneficial mutation (1.05) | 0.000 | 0.000 | 0.014 | 0.010 | 0.072 | 0.000 | 0.000 | 0.014 | 0.039 | 0.133 |
| | Neutral mutation (1) | 0.002 | 0.010 | 0.003 | 0.133 | 0.234 | 0.000 | 0.002 | 0.016 | 0.037 | 0.072 |
| $10^{-5}$ | Detrimental mutation (0.9) | 0.331 | 0.464 | 0.509 | 0.615 | 0.651 | 0.004 | 0.004 | 0.034 | 0.030 | 0.053 |
| | Detrimental mutation (0.8) | 0.846 | 0.926 | 0.915 | 0.942 | 0.873 | 0.014 | 0.006 | 0.010 | 0.016 | 0.046 |
| | Detrimental mutation (0.7) | 0.978 | 0.990 | 0.986 | 0.972 | 0.948 | 0.010 | 0.006 | 0.004 | 0.020 | 0.028 |
| | Detrimental mutation (0.6) | 0.996 | 0.992 | 0.996 | 0.988 | 0.980 | 0.004 | 0.006 | 0.002 | 0.006 | 0.018 |
| | Beneficial mutation (1.1) | 0.014 | 0.025 | 0.006 | 0.004 | 0.014 | 0.260 | 0.485 | 0.384 | 0.683 | 0.688 |
| | Beneficial mutation (1.05) | 0.031 | 0.019 | 0.012 | 0.011 | 0.022 | 0.437 | 0.521 | 0.531 | 0.672 | 0.819 |
| | Neutral mutation (1) | 0.042 | 0.032 | 0.014 | 0.011 | 0.019 | 0.681 | 0.754 | 0.822 | 0.812 | 0.898 |
| $10^{-6}$ | Detrimental mutation (0.9) | 0.018 | 0.032 | 0.030 | 0.042 | 0.032 | 0.954 | 0.938 | 0.962 | 0.933 | 0.952 |
| | Detrimental mutation (0.8) | 0.060 | 0.052 | 0.038 | 0.069 | 0.089 | 0.936 | 0.940 | 0.960 | 0.929 | 0.911 |
| | Detrimental mutation (0.7) | 0.118 | 0.132 | 0.132 | 0.152 | 0.150 | 0.880 | 0.868 | 0.868 | 0.848 | 0.850 |
| | Detrimental mutation (0.6) | 0.252 | 0.290 | 0.298 | 0.304 | 0.338 | 0.748 | 0.710 | 0.700 | 0.696 | 0.662 |

Supplementary Table 14
